# Clonal memory of cell division in humans diverges between healthy haematopoiesis and acute myeloid leukaemia

**DOI:** 10.1101/2025.06.24.660535

**Authors:** A Donada, G Hermange, T Tocci, A Midoun, G Prevedello, L Hadj Abed, D Dupré, W Sun, I Milo, S Tenreira Bento, C Pospori, A Innes, C Willekens, J Vargaftig, D Michonneau, C Lo Celso, N Servant, KR Duffy, H Isambert, PH Cournède, L Laplane, L Perié

## Abstract

Clonal memory, a cellular property inherited across at least two divisions, has emerged as a key driver of cell heterogeneity. To uncover its roles in human haematopoiesis, we developed high-resolution *ex vivo* tools that track both division and fate commitment of individual primary human haematopoietic stem and progenitor cells (HSPCs). We show that human HSPCs display a clonal memory of division, as cells descending from the same ancestor cell divide synchronously over multiple generations. In parallel, HSPCs inherit a clonal memory of fate commitment, independently of lineage identity. Both forms of clonal memory persist over at least two divisions, across different HSPC commitment stages and cell culture conditions. In contrast, malignant haematopoiesis exhibits lower synchronicity, revealing a disruption of clonal memory in leukemic cells. Epigenetic remodelling using a bromodomain inhibitor partially restores the clonal memory in division in leukemic HSPCs, highlighting the plasticity of this trait and its potential for therapeutic modulation. Our findings position clonal memory as a key regulator of human haematopoietic stem cell behaviour. Demonstrating that clonal memory can be modulated opens new avenues for tuning cell heterogeneity in healthy and pathological tissues.

## Introduction

Stem cells, and how they proliferate, play a key role in tissue development, homeostasis, and regeneration. However, not all stem cells divide or die at the same rate^1^. These proliferation differences impact the cellular composition of tissues and thereby tissue formation and renewal over time. In cancer, changes in cell division directly affect the fitness of cancer cells, leading to their clonal expansion, increased malignant phenotypes and poorer prognosis^2^. Currently, we still know little about what drives differences in stem cell proliferation in healthy or diseased tissues.

Recently, clonal memory has emerged as a driver of stem cell heterogeneity^3–7^. Clonal memory is defined as a cellular property inherited over at least two successive rounds of cell division. Despite its importance in stem cell heterogeneity, the role of clonal memory in proliferation heterogeneity is poorly understood. Additionally, fundamental questions about clonal memory remain, including how long it persists over time (e.g. hours^8^, days^9^, months^10^) and its long-term functional impact. Although clonal memory is an intrinsic property, whether and how extrinsic factors (e.g. microenvironment, soluble factors) could modulate it remains unknown. Modulating clonal memory is crucial as it could promote the stability required for homeostasis by transmitting cellular traits across multiple cell generations. However, this stability could come at the expense of adaptability to environment changes. Thus, understanding how clonal memory contributes to heterogeneity in stem cell division is critical to determining its role in balancing stability and adaptability in both homeostasis and disease.

Haematopoiesis is a good model to address the role of clonal memory. It is a hierarchically organized differentiating system that generates over 13 different cell types across tens of divisions^11^. HSPCs are a heterogeneous population with diverse division and differentiation abilities^12^. In mice, clonal memory of differentiation partly contributes to this heterogeneity. Indeed, HSPCs show heterogeneous differentiation biases that are maintained and transmitted throughout cell division and differentiation^13^, as shown by single-cell lineage tracing^3^ and serial transplantations^10^. However, most clonal memory studies have focused on differentiation, while its influence on HSPC proliferation^14^, in particular in human cells, remains unexplored. Moreover, as differentiation tends to be coupled with division^15,16^ and both properties are central to stem cell biology, the clonal memory of both processes needs to be studied together. Critical for this aim is the ability to simultaneously track division and differentiation within individual cell lineages, using lineage tracing tools such as the MultiGen assay^14,17–19^.

In pathological human haematopoiesis, evidence for clonal memory of division and fate in HSPCs remains largely indirect. While stem cell bone marrow transplantation induces cell proliferation to repopulate the recipient bone marrow, the observation that stem cells accumulate mutations at similar rates in both recipient and donor support the hypothesis that stem cell proliferation is an intrinsic properties^20–22^. In clonal haematopoiesis, known genetic drivers account only for approximately 20% of all age-related clonal expansions, indicating that while genetic mutations contribute to proliferation heterogeneity, other intrinsic and extrinsic factors are at play^23^. In humanized mouse models, cellular memory was demonstrated to impact the heterogeneity in clonal fitness for otherwise genetically identical acute myeloid leukaemia (AML) stem cells, shaping their clonal evolution^3^, and leading to differences in treatment resistance^24,25^. However, whether clonal memory can be modulated in leukaemia and leveraged for therapeutic benefit remains unexplored.

Here, we address whether clonal memory plays a role in human haematopoiesis, by studying clonal memory in primary human HSPCs during healthy and malignant haematopoiesis, focusing on division. We developed several tools to follow division, differentiation, and lineage of individual human HSPC *ex vivo*. Using live-cell microscopy, we establish that human HSPCs display a clonal memory of cell division over at least two division and three cell generations, across different HSPCs commitment stages and several culture conditions. Further, we demonstrate that there are two distinct inherited clonal memory properties, one for division and one for fate commitment. Counterintuitively, we find that HSPCs in malignant AML haematopoiesis show altered clonal memory in division, with a decreased level of synchronicity within cell families. By manipulating clonal memory of division through inhibition of epigenetic remodelling in AML cells, we observe increased clonal memory but no changes in the total number of divisions, suggesting that clonal memory is regulated independently of the number of divisions. Altogether, we demonstrate the existence of distinct clonal memories that govern division and lineage commitment in healthy human primary HSPCs. Furthermore, we show that while the clonal memory of division is altered in AML, it remains amenable to modulation. This ability to modulate clonal memory opens new avenues for tuning cell heterogeneity.

## Results

### Clonal memory of division in human HSPC families

To measure the heterogeneity in division in human HSPCs and their descendant cells, we analysed three types of HSPCs, via single-cell live-cell imaging. We isolated hundreds of immature progenitors, defined as CD90^+^ (Lin^−^CD34^+^CD38^−^CD90^+^CD45RA^−^), CD90^-^ (Lin^−^ CD34^+^CD38^−^CD90^−^CD45RA^−^), and haematopoietic progenitor cells (HPCs; Lin^−^CD34^+^CD38^+^) from umbilical cord blood and adult bone marrow from unrelated donors **(Extended Data Fig. 1a).** We then cultured single HSPCs for 96 hours in two serum-free media: one to promote cell division and differentiation towards the myeloid and erythroid lineages (Diff)^26^, and one to maintain the immature CD34^+^ phenotype (gene therapy medium, GT)^27^. Analysing 1973 cell families, each composed of the descendants of a single HSPC, we manually extracted the time necessary to complete the first three divisions from more than 5000 cells **(Fig. 1a)**. First, we observed differences of proliferation between cell types, with HPCs dividing faster and forming bigger colonies at 96 hours than CD90^+^ or CD90^-^, for multiple donors in both culture media **(Fig. 1b-c, Extended Data Fig. 2m)**. Past the first division **(Extended Data Fig. 2a-b)**, the time to divide shortened not only for CD90^+^ and CD90^-^ **(Extended Data Fig. 2d-e, 2g-h),** as previously described^28^, but also for HPCs. Therefore, cellular quiescence alone was not responsible for this shortened division time, as HPCs are not quiescent^29,30^ **(Extended Data Fig. 2j-k)**. Additionally, cells of the same type displayed heterogenous division times, as previously reported^31^. The heterogeneity was of similar amplitude between cell types (i.e., CD90^+^s, CD90^-^s, and HPCs) and culture media, although adult bone marrow HSPCs **(Extended Data Fig. 2c, 2f, 2i, 2l)** displayed more heterogeneity than foetal cord blood HSPCs.

**Figure 1.**
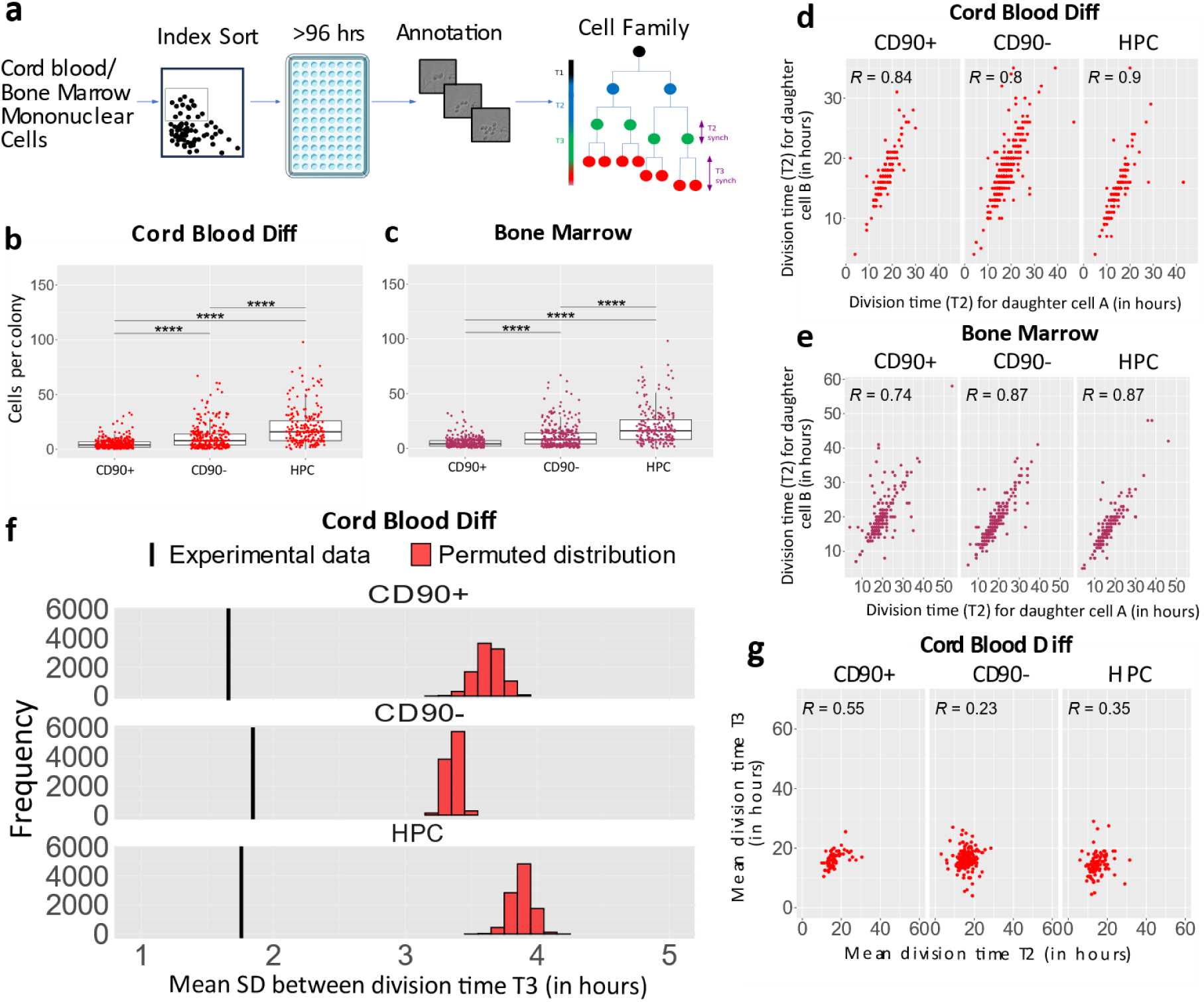
Division in synchronicity within cell family. **a)** Overview of the live cell imaging experimental workflow: individual progenitors were index sorting via FACS in 96-well tissue plates and kept in culture for at least 96 hours. Division dynamics were manually annotated. **b)** Colony size, measured as number of light-refringent cells in the well at T = 96 hours, for foetal cord blood HSPCs expanded in Diff medium, in 7 independent experiments (n = 166 for CD90^+^, n = 369 for CD90^-^ and n = 157 for HPC). **c)** Colony size, measured as number of light-refringent cells in the well at T = 96 hours, for bone marrow HSPCs, in 8 independent experiments (n = 356 for CD90^+^, n = 328 for CD90^-^ and n = 222 for HPCs). Dunn post-hoc test with Hochberg correction, ****p < 1×10^-4^. **d-e)** Correlation in division time (expressed in hours) between two sister cells in generation 1, for **d)** foetal HSPCs isolated from cord blood and expanded in Diff medium (7 independent experiments, n = 149 for CD90^+^, n = 335 for CD90^-^ and n = 144 for HPC), and **e)** adult bone marrow cells (8 independent experiments, n = 255 for CD90^+^, n = 269 for CD90^-^ and n = 205 for HPC). Spearman’s rank correlation coefficient R. **f)** Mean SD of third division time (T3) of cells derived from the same ancestor cells (black line), and permuted families (red histograms), for HSPCs from foetal cord blood and expanded in Diff medium. Only families with four recorded T3 divisions were used. Seven independent experiments; n = 65 for CD90^+^, n = 170 for CD90^-^ and n = 99 for HPC), p values < 1×10^-4^. **g)** Correlation in division time (expressed in hours) between the mean division time T2 and the mean division time T3, for HSPCs isolated from foetal cord blood and expanded in Diff medium. Spearman’s rank correlation coefficient R.

**Figure 2.**
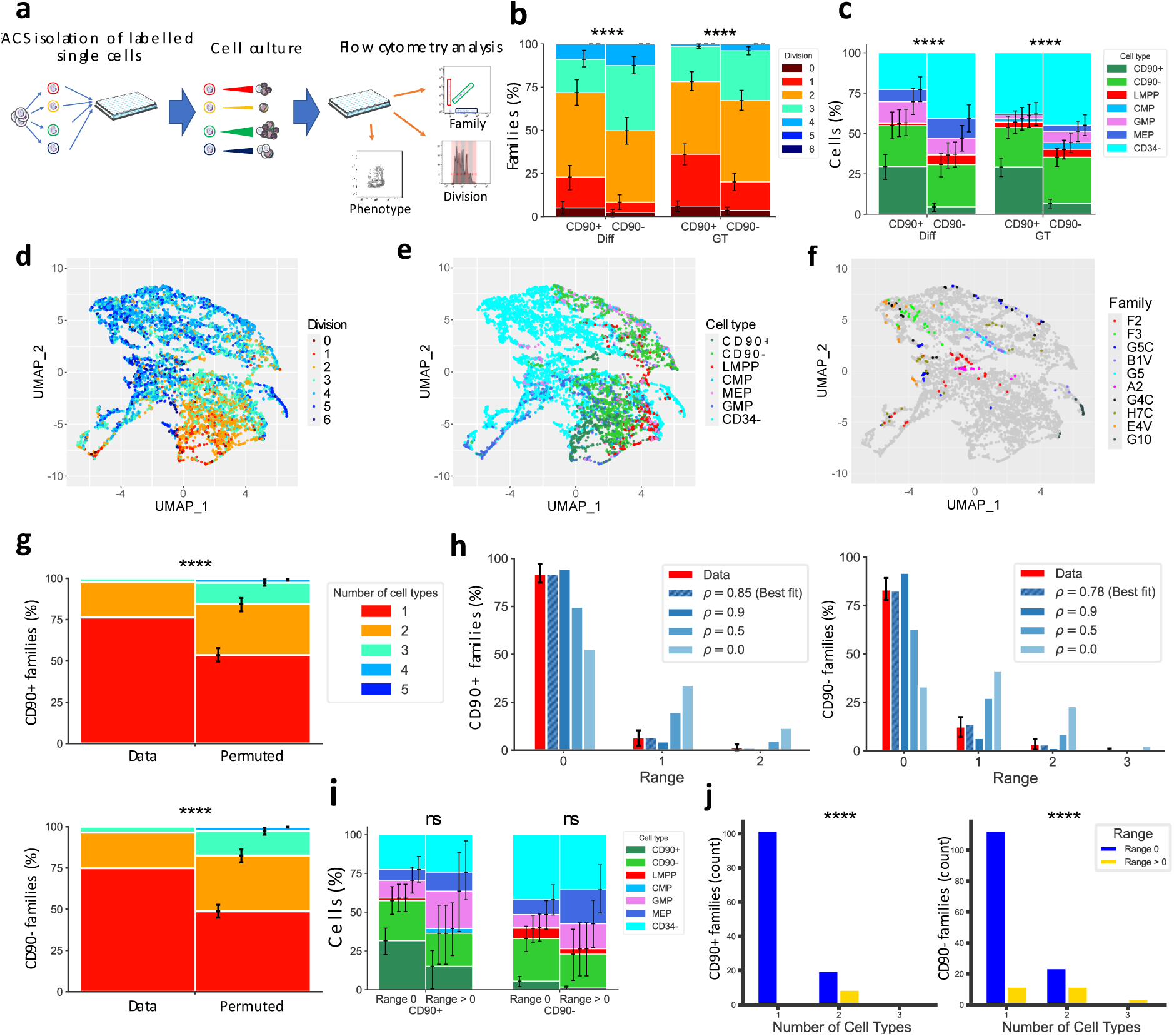
Clonal memory of fate and division in human HSPCs. **a)** MultiGen experimental workflow: HSPCs stained with four combinations of CFSE and CTV were sorted in 96-wells tissue plates: four cells per well, each one from a different CTV/CFSE combination. After 72 and 96 hours in culture, plates were analysed by flow cytometry. **b)** Distribution of the maximum division reached by individual families, derived from CD90^+^ and CD90^-^ and expanded in two media (Diff and GT) after 72 hours (7 independent experiments, n = 235 for CD90^+^ GT, n = 248 for CD90^-^ GT, n = 135 for CD90^+^ Diff, n = 167 for CD90^-^ Diff). **c)** Proportions of cell types per families derived from CD90^+^ and CD90^-^, cultured in the GT and Diff media for 72 hours. **d-f)**, UMAP projection of 6148 individual cells from 13 independent experiments, colour coded by **d)** division (based on CTV/CFSE fluorescence), **e)** surface markers-based phenotype and **f)** family (displaying the ten largest families). **g)** Distribution of the number of cell types per family, compared with 250,000 permutations of the experimental data (CD90^+^ and CD90^-^ in Diff medium, after 72 hours). **h)** Percentage of families per division range, defined as the difference between divisions within each family (range = 0 means that all the cells in the family divided the same number of times). The red bar corresponds to the experimental dataset (135 CD90^+^ and 167 CD90^-^ families, in Diff medium for 72 hours). The dashed blue bar corresponds to the best fit to the experimental data of the correlation coefficient in division ρ. The filled blue bars are simulations parametrized by the indicated ρ value. **i)** Proportions of cell types per families in division range = 0 or > 0, derived from CD90^+^ or CD90^-^ cultured in the Diff media for 72 hours. **j)** Number of progenitor subsets per family, for families with range = 0 or > 0. For the panels **b, c, g, h** and **i** the error bars indicate 95% confidence intervals calculated via basic bootstrap. The p values are calculated using permutation testing, **** p < 1×10^-4^.

To assess the existence of clonal memory of division, we next explored the heterogeneity in division times within cell families. To this end, we measured the difference in division timing between cells in the same generation: sister cells (cells sharing the same mother cell) and cousin cells (cells sharing the same grandmother cell). Cells within a family divided synchronously **(Fig. 1d),** as sister cell division times were highly correlated for all tested progenitors (CD90^+^s, CD90^-^s, and HPCs) and culture media, accordingly to the calculated Spearman ranked correlation coefficient **(Extended Data Fig. 3a)**. Furthermore, cousin cells also divided synchronously **(Fig. 1f)**, with the four division for a given set of cousin cells displaying a lower standard deviation compared to permuted families composed of unrelated cells. This synchronicity in division was present in all HSPC types, irrespective of their origin **(Fig. 1e, Extended Data Fig. 3c)** or culture media **(Extended Data Fig. 3b)**, suggesting a clonal memory of division across commitment stages. However, cell division time across generations correlated weakly **(Fig. 1g),** as in other cell systems. This was similar for each HSPC type, irrespective of culture media **(Extended Data Fig. 3d-e, 3g-h, 3j)** or cell origin **(Extended Data Fig. 3f, 3i, 3k)**. In summary, we observed a significant and conserved intragenerational synchrony in cell division timing, over different HSPC commitment stages, and across different culture conditions.

**Figure 3.**
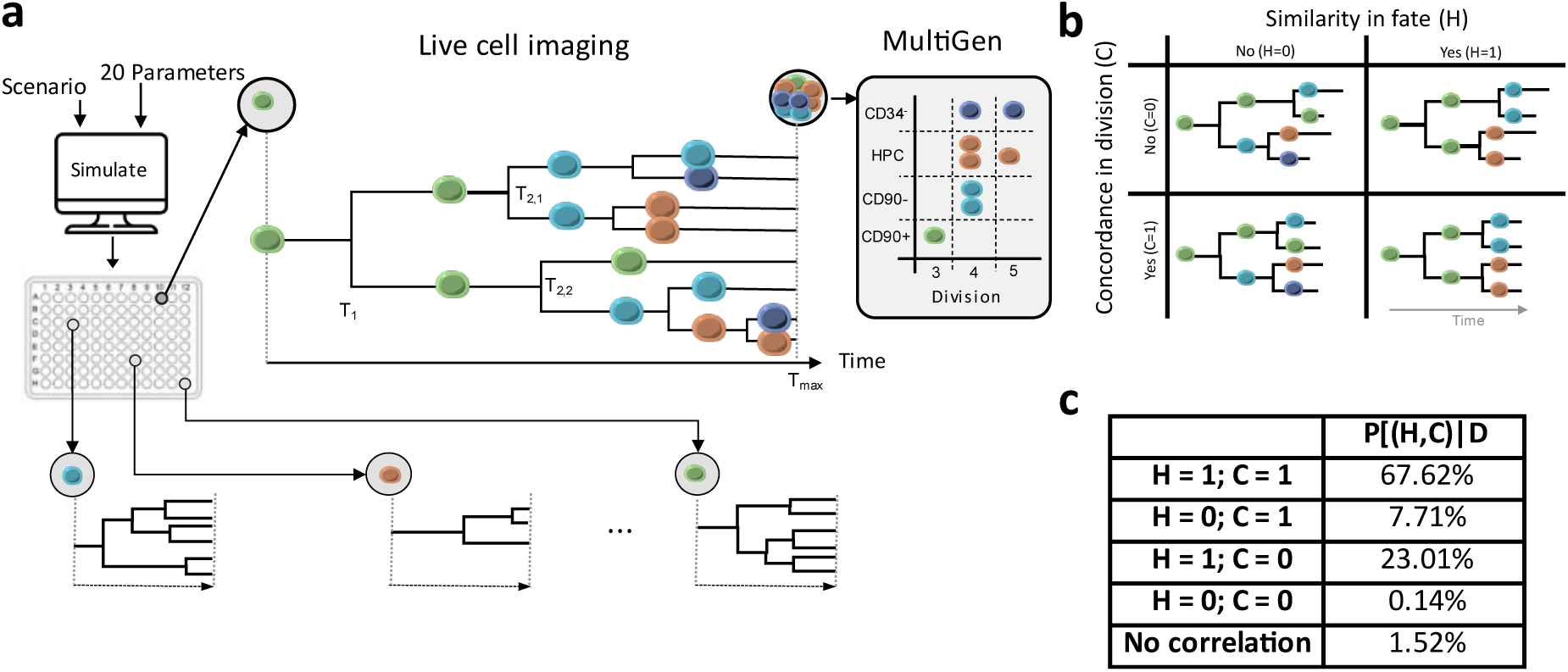
Two distinct clonal memories of division and lineage commitment. **a)** Overview of the modelling strategy: proliferation and differentiation for each well were simulated by agent-based modelling for different scenarios. For each simulation, simulated data were extracted to be compared with both the live-cell and MultiGen datasets. From the live-cell imaging datasets, the experimental parameters used were the values of time of 1^st^ (T_1_), 2^nd^ (T_2,1_ and T_2,2_) and 3^rd^ divisions (T_3,1_ to T_3,4_) and the colony size at T_max_. From the MultiGen datasets, the relative percentages of cell types and the associated cell divisions were used. Green, cyan, orange and dark blue correspond to CD90^+^, CD90^-^, HPC, and CD34^−^ types, respectively. **b)** The four distinct modelling scenarios tested, based on whether there is similarity in fate (H=1) or not (H=0), and whether there is concordance in division (C=1) or not (C=0). **c)** Value of the posterior probability of each scenario ℙ[(*H*, *C*) |𝒟], given the experimental observations 𝒟, after running the ABC (Approximate Bayesian Computation) procedure. We included also a fifth modelling scenario assuming no cell memory of cell division and cell differentiation and time-to-divide distributions that are independent of the dividing cell type.

### Clonal memory of division and lineage commitment in human HSPCs

To determine if human HSPCs display a clonal memory of lineage commitment, we performed the MultiGen assay^14,17,18^ to track simultaneously the cell family, the number of cell divisions and the phenotype of each cell produced by individual HSPCs **(Fig. 2a)**. We analysed 1634 individual CD90^+^- and CD90^-^-derived families, from human cord blood, after 72 hours and 96 hours and using both Diff and GT media. Following the MultiGen assay, we seeded 4 individual progenitors per each well, all labelled with a different division tracer colour, to allow the correct identification of the cells sharing the same ancestry. We assigned the progeny phenotype using an antibody panel that identifies six different HSPC subtypes **(Extended Data Fig. 1b)**. At the population level, we observed that CD90^-^ families divided and differentiated more often than CD90^+^ families **(Fig. 2b-c, Extended Data Fig. 4a-b)**, as expected from their position in the haematopoietic hierarchy. Consistent with previous observations^14^, cells that divided more were also more likely to be more differentiated than the ones that divided less **(Extended Data Fig. 4c-j)**. Moreover, we observed only a minority of cells retaining their initial phenotype (CD90^+^ or CD90^-^) after the first two divisions, confirming that most cells undergo differentiation instead of self-renewal. The vast majority of immature HSPCs (i.e., CD90^+^s and CD90^-^s) were associated with fewer divisions, while committed progenitors and precursors emerged with more divisions **(Fig. 2d-e)**. We observed substantial heterogeneity in lineage commitments and divisions between families of the same progenitor cell type **(Fig. 2c, Extended Data Fig. 4k-r)**, in line with the current state of the art^32^. Therefore, our experimental set-up recapitulates the heterogeneity of the lineage commitment and division found in human HSPCs.

Next, we explored the heterogeneity in lineage commitment within cell families. Cells from the same family consistently co-localized on the UMAP representation of the HSPC subsets detected via flow cytometry **(Fig. 2f).** Most of the families displayed only one subset **(Fig. 2g),** despite the detection of multiple cell types in our culture system and the multipotent nature^33^ of CD90^+^ and CD90^-^. Permutation testing revealed significant intrafamilial similarities in lineage commitment, more than expected if each cell was committing independently of the sister and cousin cells. This observation was independent of the ancestor cell type, the timing of the analysis, and the culture condition **(Extended Data Fig. 5a-c)**. We concluded that a clonal memory of lineage commitment in human HSPCs is present over different HSPC commitment stages and across culture conditions.

We also observed clonal memory of division with the MultiGen assay, consistent with our results using live-imaging microscopy **(Fig. 1)**. For each family, we compiled a metric called “range”, which is the difference in the number of divisions performed by cells sharing the same ancestor. Most dividing families displayed high concordance in division: for 786 CD90^+^ and 848 CD90^-^ families, all cells within the same family had the same division number (range = 0) after 72 and 96 hours. Only 25% of families were spread over more than one generation (range > 1), over all our datasets **(Fig. 2h, Extended Data Fig. 5d-f)**. As cell death and sampling effects could potentially make families look more concordant in division, we developed an improved version of the previously published mathematical model^14^ to account for sampling and cell death. We did not detect cell death during the first 96 h, using live-imaging data, and we estimated a recovery rate of 0.6 **(Extended Data Fig. 1c)**. We fitted the model to the family distribution over range, to estimate the correlation in division progression decisions among cells within a family. High correlation coefficients (70–85%) **(Fig. 2h)** resulted in the best fit to the measured familial ranges, demonstrating that cell families progress through division together. We used permutation testing to assess if families sharing the same microwell would influence each other, for their properties of cell division and differentiation. We did not see such an effect, as families sharing the same well retained both different division times and cell type composition **(Extended Data Table 1).** These results suggest the existence of a cell-intrinsic clonal memory of division and lineage commitment over several divisions.

### Two distinct clonal memories for division and lineage commitment within cell families

While we observed clonal memory in division and lineage commitment within HSPC families, we also observed that lineage commitment progresses with division. Therefore, these factors could influence each other: for example, the concordance in division being a consequence of lineage commitment progressing with division or vice-versa. In this line of thought, while concordant and discordant families showed the same diversity of cell subsets **(Fig. 2i, Extended Data Fig. 5g-i)**, individual discordant families (range > 0) displayed more progenitor subsets than concordant families **(Fig. 2j, Extended Data Fig. 5j-l)**. We next tested whether the presence of one clonal memory of division or lineage commitment or both would best explain the similarity in lineage commitment and concordance in division. To this end, we applied a stochastic modelling framework with approximate Bayesian computation (ABC)^34–36^ to the live-cell imaging and the MultiGen data combined. Briefly, we built a generic model of the HSPCs division and differentiation over 96 h **(Fig. 3a)** starting from one cell of a known phenotype in a well and considering four possible phenotypes: CD90^+^, CD90^-^, HPC, or CD34^−^. In the model cells divided at random times, according to log-normal probability distributions^37,38^ for each cell type. Differentiation only happened after division^39^ and de-differentiation was not allowed. We tested distinct modelling scenarios, whereby cells from the same family could be similar (or not) in lineage commitment and/or concordant in division **(Fig. 3b)**. In all scenarios, symmetric and asymmetric division were allowed. In the scenarios where we modelled the similarity in lineage commitment, we introduced - as a parameter - the strength of this similarity in lineage commitment between daughter cells. To compare how the different modelling scenarios fitted the entire dataset, we used a distance that aggregated the derivation of many summary statistics. We ran simulations with different parametrizations of the models and then compared the posterior probability of those models. We found that the models lacking similarity in commitment within the cell family were inconsistent with the data. The best model, with the higher posterior probability, displayed both similarity in commitment and concordance in division for cells from the same family. The model in which cells from the same family were only similar in commitment but not concordant in division had a lower posterior probability **(Fig. 3c).** As a control, we tested if a minimal scenario with no correlation between division and differentiation and no clonal memory could explain the data, we assumed that the probability distributions of the time to divide was identical across cell types. Unsurprisingly, this scenario had a lower posteriori probability, making it unlikely to explain our data. Overall, these results demonstrated that two distinct clonal memories for division and lineage commitment underlie the similarity in lineage commitment and concordance in division.

### Clonal transcriptional programs of clonal memory in human HSPCs

To explore the transcriptional programs involved in the clonal memory of human HSPCs, we developed an experimental protocol integrating lineage tracing and molecular profiling with the division history of each cell. Briefly, we FACS-sorted individual CD90^-^s from two cord blood samples and cultured them *ex vivo* in the Diff media while recording their division history by live-cell imaging over 90 hours. We then re-isolated individual cells from randomly selected wells, using low throughput, multi-fluorescence image sorting^40^, and profiled each cell with high-coverage single-cell RNA sequencing (scRNA-seq) via SMART-seq3^41^ **(Fig. 4a)**. We obtained two multidimensional datasets containing information on division, family, and transcriptome for 142 individual cells from 30 families after QC and filtering **(Fig. 4b, Extended Data Fig. 6d, Extended Data Table 2)**. After supervised annotation using known marker genes, we identified several cell subsets at different stages of myelo-erythroid differentiation: immature progenitors (HSPCs), erythroid-megakaryocytic progenitors (EryP/MkP) and myelomonocytic committed progenitors (MyP and GraP) **(Fig. 4c-d, Extended Data Fig. 6e-g)**. These subsets globally matched the subsets we identified via CITE-seq (cellular indexing of transcriptomes and epitopes by sequencing) profiling of 8,580 cord blood HSPCs after 4 days in Diff media (**Extended Data Fig. 6a-c**).

**Figure 4.**
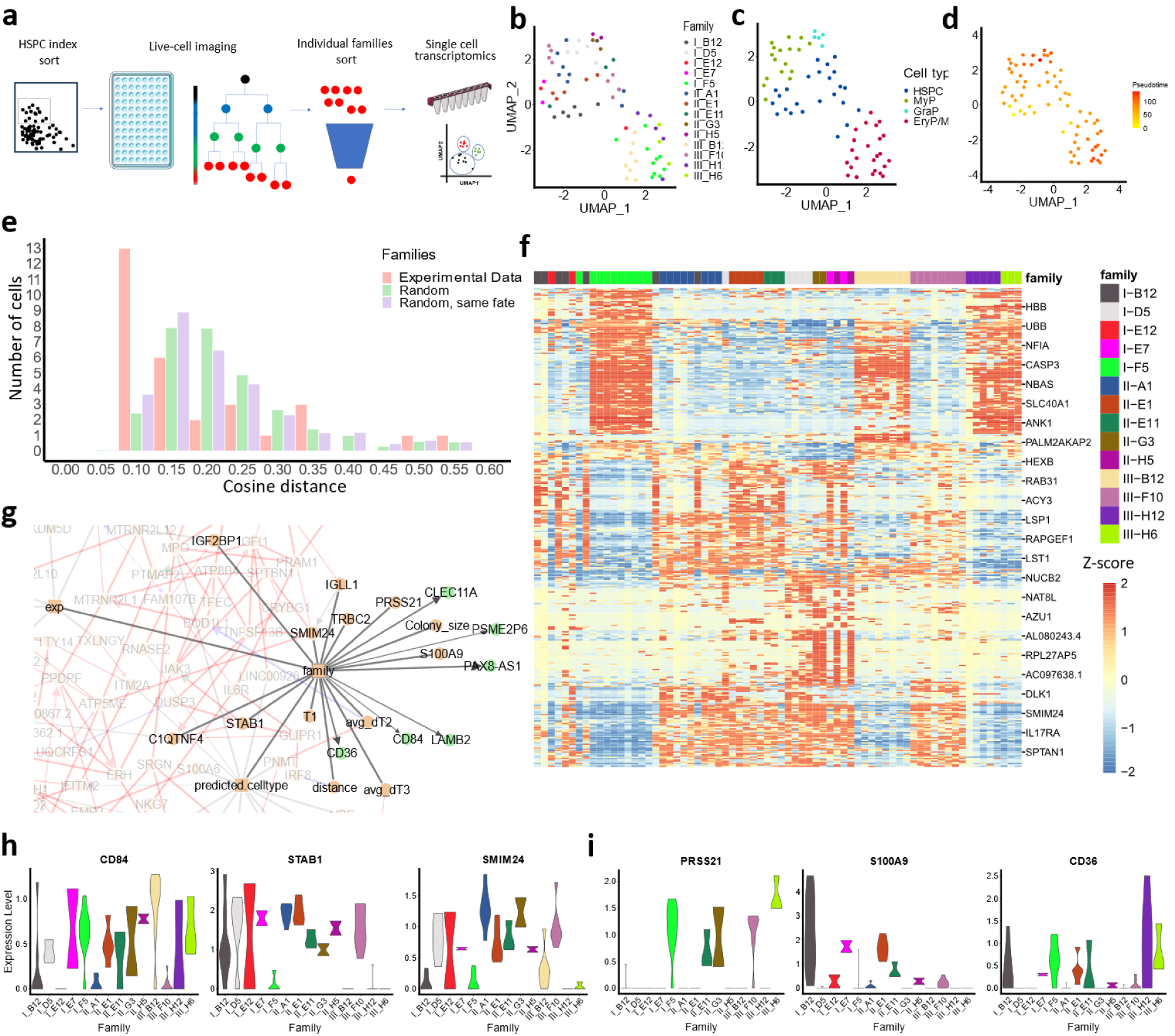
Clonal transcriptional programs of clonal memory in human HSPCs. **a)** Scheme of the experimental workflow: individual CD90^-^ were FACS-sorted in 96-well plates and live-imaged for 90 hours (Diff medium). Then, individual cells from 30 randomly selected families were re-sorted by image-based sorting into 384-wells plates and processed by SMART-Seq3. **b-c-d)** UMAP representations of a transcriptomic dataset (70 cells), colour-coded by **b)** family and **c)** annotated cell type, one experiment representative of two and **d)** pseudotime. **e)** Distribution of the gene expression median cosine distance for real cell families (red) and two permutation-generated distributions, either retaining (purple) or discarding (green) the cell type of the permuted cells. Wilcoxon rank-sum test, p value < 0.0035 for family/permutation discarding cell type and p value < 0.02 for family/permutation retaining cell type. **f)** Heatmap displaying the z-score of the 371 most variant genes (rows) between families. 24 randomly selected gene names are displayed on the right side. Cells (columns) are colour-coded according to their original family. **g)** Snapshot of the MIIC network generated using transcriptomic and divisional data, highlighting the features directly connected to Family. Variables in orange are causal regarding Family, those in green are consequential. Edges represent an association between two variables, while edges with an arrow represent a causal relationship. Red edges highlight positive association, blue edges represent negative associations, and grey edges represent categorical association with no sign. A green arrow indicates genuine causality, while a red, blue or grey arrow stands for putative causality (i.e. the presence of an unobserved common cause between the two variables cannot be ruled out). Full network with detailed legend is shown in Extended Data Figure 8a. **h-i)** Dataset A expression of a selection of “family”-associated genes accordingly to the MIIC analysis, displayed per individual family and organised as **h)** genes expressed ubiquitously and **i)** genes expressed only in a subset of families.

We next explored the similarity of cell families in the transcriptomics space. We observed that cell families co-localized within the same UMAP clusters, suggesting that cell from the same family share a similar transcriptome **(Fig. 4b-c, Extended Data Fig. 6d-e)**. To quantify the transcriptomic similarity between cells from the same family, we compared the median cosine distance^42^ distributions between the observed cell families and artificial families made of permutated cells. Cell families had lower median cosine distance than permutated families **(Fig. 4e)**, demonstrating that cells within a family are more transcriptionally similar than cells between families, in line with the MultiGen results. Moreover, this result held after permutating cells within the same cell type subsets **(Fig. 4e),** suggesting that transcripts associated with cell subsets alone were insufficient to explain this similarity. Thus, biological processes other than those associated with lineage commitment contributed to the similarities between the cells within a family.

To characterize the transcriptional programs associated with the familial similarity^9,43^, and therefore with clonal memory, we extracted the genes that varied the least within a family and the most between families **(Fig. 4f, Extended Data Fig. 6i)**. We could then identify transcriptional fingerprints specific for each family, contributing to the unique intrafamilial similarities beyond fate commitment. These signatures contained genes associated to many biological processes, including membrane trafficking (e.g., *RAB31, CAVIN2, VAMP8*), cell proliferation (e.g., *CASP3, IGFBP2*), cytoskeleton structure (e.g., *LESP1, ANK1, MYH10)*, cell signalling (e.g., *IL17RA, IL1RL1, LAT2*), and metabolism and biosynthesis (e.g., *DLK1, CALR, XBP1*) **(Fig. 4f; Extended Data Table 3)**. To identify common genes between families that contributed to clonal memory, we ran the Multivariate Information-based Inductive Causation (MIIC) causal network reconstruction algorithm^44^ on both transcriptomic datasets using the transcriptomes, the family and the division information from live imaging of each cell **(Fig. 4g; Extended Data Fig. 7a, Extended Data Table 4)**. The MIIC algorithm is a multivariate information-based network reconstruction method^45,46^ that reconstructs causal and non-causal graphical models from heterogeneous data. The MIIC analysis identified the variables “family” (sharing the same ancestry), “predicted cell type” and “distance” (transcriptomic similarity) as key hubs in the networks **(Extended Data Fig. 7b)**. Gene expression of *IGLL1* and surface markers like *CD36* and *CD84*, amongst others, were strongly associated with the “family” hub **(Fig. 4g)**. Notably, the transmembrane protein STAB1, which directly connected with the “family” hub, emerged as a central hub within the gene network, orchestrating interactions with a broad array of other genes **(Extended Data Fig. 7c)**. In contrast, variables associated with division properties, such as division time and colony size, were only directly connected to less than 5 genes, and were strongly related to the “family” variable **(Extended Data Fig. 7d**) as all these families were synchronous in division **(Fig. 1d-f)**. The limited number of genes associated with division-related properties suggests that the transcriptome is not a suitable proxy for these properties, or that the relevant genes are expressed at low levels and thus remain undetected. Finally, the genes associated to the variable “family” were not equally expressed by all families **(Fig. 4g-h, Extended Data Fig. 7e-f)**, suggesting the existence of different transcriptomic program between families. Some genes were expressed in all families but at different levels **(Fig. 4h)**, while others were specific to some families **(Fig. 4i)**. In summary, human HSPC families displayed heritable transcriptomes linked to several cellular processes, including cell commitment, that contributed to clonal memory.

### The clonal memory of division is disrupted in leukemic cells

AML is a proliferative disease with a differentiation blockade, resulting in a disproportionate production of blast cells. To test if this disruption affects the cellular clonal memory, we analysed primary CD34^+^ blasts from AML patients **(Extended Data Table 5),** that had been isolated from bone marrow or peripheral blood and cultured for at least 72 hours in Diff media supplemented with IL-1β^47^, using live-cell imaging and MultiGen. By live-cell imaging, the intragenerational correlations in the division time between daughter and cousin cells were diminished **(Fig. 5a, Extended Data Fig. 8a).** As expected, blasts displayed increased number of divisions **(Extended Data Fig. 8c)** and division time **(Extended Data Fig. 8d-f)** when compared with cells from healthy controls. The MultiGen assay showed that CD34^+^ AML cells possessed a lower clonal memory of division than healthy HSPCs, as the concordance in division ρ was lower than for control cells from healthy blood donors **(Fig. 5b).** Stratifying the analysis per ancestor cell types (i.e., CD90^+^s, CD90^-^s, HPCs), we measured clonal memory of division reduction for all cell types **(Extended Data Fig. 8b, 8i-j),** even if we did not observe significant changes in the differentiation profile **(Extended Data Fig. 8g)** or in the number of divisions over the 72 hours **(Extended Data Fig. 8h)**. We excluded the impact of genetic heterogeneity, as we measured synchronicity for sisters/cousin cells, all sharing the same ancestor genomic background. These results revealed that primary CD34^+^ blasts from AML patients had a lower synchronicity of division within families.

**Figure 5.**
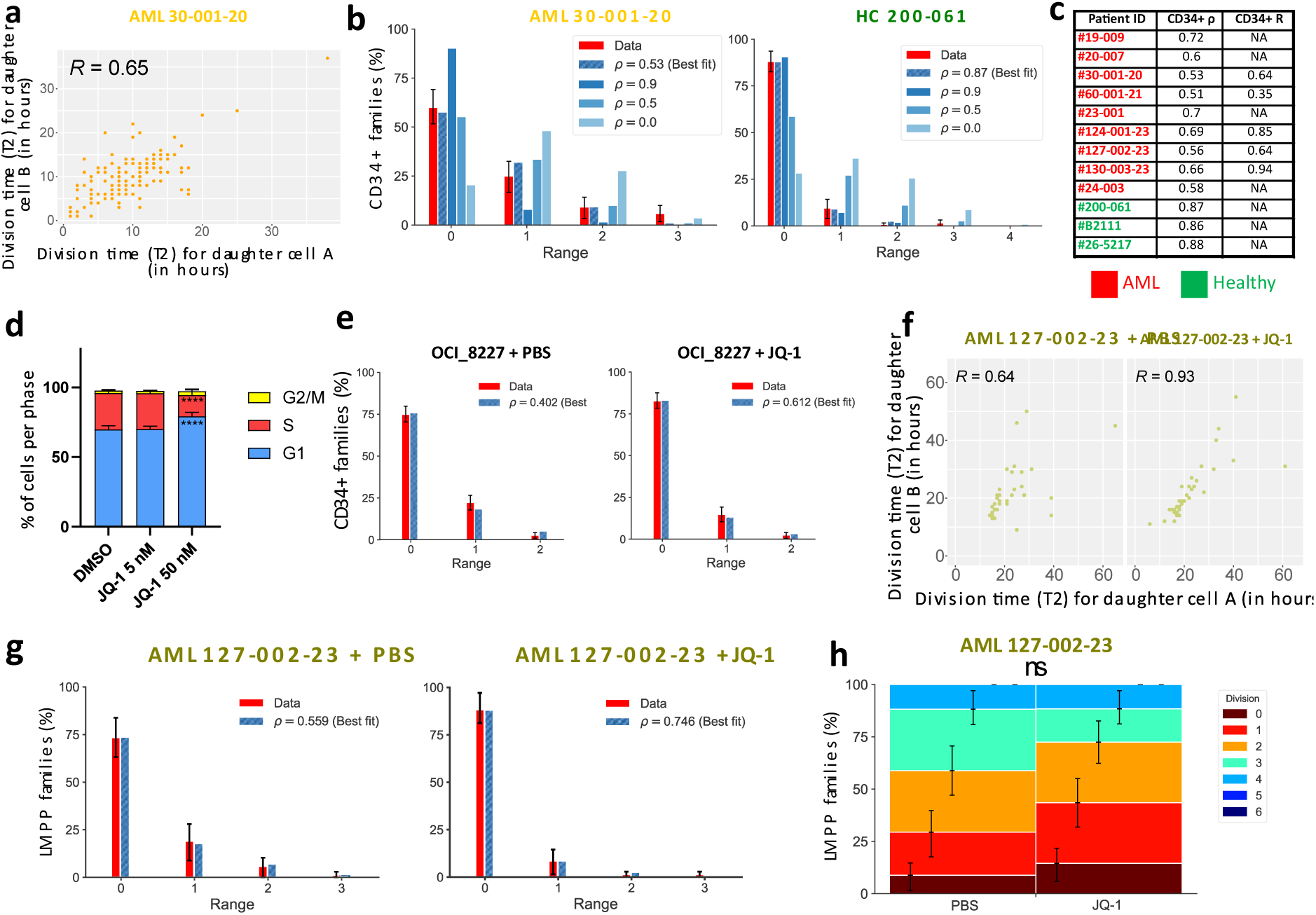
Reduced concordance in division of leukemic families can be restored. **a)** Correlation in division time (in hours) between sister cells after 1 division, for CD34^+^-derived families from AML patient #30-001-20. Spearman’s rank correlation coefficient R (n = 154). **b)** Percentage of CD34^+^-derived families per division range, for the same patient and a healthy control (HC, #200-061), after 72 hours. The red bar represents the data, the dashed blue bar represents the correlation coefficient ρ best fit, the filled blue bars represent simulations for different parametrization of ρ, n = 120. **c)** Summary of correlation coefficients as in **a)** and **b)** for 9 AML samples (red) and three healthy controls (green), analysed with live-cell imaging, MultiGen or both. NA for not available. **d)** Proportions of OCI_8227 cells in G1-, S- and G2/M-phase, after 72 hours exposure to JQ-1 (5 and 50 nM), 2-way ANOVA test error bars indicate SD, **** for p < 0.001, n = 3. **e)** Percentage of families per division range for OCI_8227 treated for 72h with PBS (control) or JQ-1 5 nM (treatment). The red bars correspond to the data (n = 250 and 327 respectively, four independent experiments), the dashed blue bar represents the ρ best fit. We used a recovery rate of 0.6 and a survival rate of 0.95. **f)** Correlation in division time (in hours) between sister cells after 1 division, for LMPP-derived families for patient #127-002-23 exposed to PBS and JQ-1 for 96 hours (n = 32 for both). Spearman’s rank correlation coefficient R. **g)** Percentage of families per division range for LMPP--derived families from patient #127-002-23, treated for 72h with sublethal doses of JQ-1 or PBS (n = 69 and n = 68 respectively). The red bars correspond to the data, the dashed blue bar represents the ρ best fit. **h)** Distribution of the maximum division reached by individual families treated for 72h with sublethal doses of JQ-1 and PBS (n = 69 and 68 respectively). For the panels **b, e, g** and **h** the error bars indicate 95% confidence intervals calculated via basic bootstrap. The p values are calculated using permutation testing, ns for p > 0.05.

We observed some variability in the synchronicity in division between patient samples **(Fig. 5c),** with best-fitted correlation coefficients of division within families varying between 53% and 72%. Nevertheless, this variability was uncorrelated to the patient mutation profile, clinical history **(Extended Data Table 5)** or the ability of the blasts to expand *ex vivo* **(Extended Data Fig. 8k-n)**. These findings demonstrated a reduction in the synchronicity of primary AML blast divisions, suggesting a disruption of the clonal memory that orchestrates cell division. Decreasing clonal memory could be a strategy for AML blasts to burst their heterogeneity in division and promote adaptability over stability.

### Clonal memory of division within AML HSPC families can be modulated

Finally, we assessed if the clonal memory in division was a plastic trait that can be modulated. As we observed that the clonal memory was independent of genetic mutations, we hypothesized an epigenetic mechanism as central to clonal memory^3,7,9,43^. To this end, we tested the effects of JQ-1, a pan-bromodomain and extra-terminal domain (BET) inhibitor^48^ on the clonal memory of AML cells.

First, we used a leukaemia stem cell (LSCs) model of differentiating AML^49,50^, the OCI_8227, to test the effect of JQ-1 on clonal memory **(Extended Data Fig. 9a)**. We selected the drug dose of 5 nM, as it induced little death and did not cause significant changes in cell cycle progression and cell division number **(Fig. 5d, Extended Data Fig. 9b)**. Without JQ-1, the OCI_8227 cells displayed a decreased clonal memory in division, similarly to primary CD34^+^ blasts, with fewer families in range 0 and low correlation coefficients of division within families. After exposure to JQ-1 (5 nM), the cell families that divided simultaneously increased by approximately 10%, and the best fitted correlation coefficient by 1.5-fold **(Fig. 5e).** We concluded that the clonal memory of cell division could be modulated, in this case increased with JQ-1. As the JQ-1 treatment affected the differentiation profile **(Extended Data Fig. 9c)** but not the total number of divisions **(Extended Data Fig. 9d)** or the clonogenicity **(Extended Data Fig. 9e),** we concluded that JQ-1 inhibition modulated only the division clonal memory but not the division capacity of cells, suggesting that these two processes are regulated by different mechanisms^51^.

We obtained similar results in primary CD34^+^ blasts from AML patients. JQ-1 treatment increased the synchronicity between sister cells during their second divisions **(Fig. 5e)** without significantly modifying their division time **(Extended Data Fig. 9f-j).** Using the MultiGen assay, JQ-1 treatment also increased the percentage of families in range 0 and the correlation coefficients of division within families **(Fig. 5g)** as compared to vehicle, without changing either the family progression through divisions **(Fig. 5h)** or their surface phenotype **(Extended Data Fig. 9j).** These results were confirmed using samples from two other patients **(Extended Data Fig. 9k-r)**. Thus, as the clonal memory in division increased *ex vivo* in primary AML CD34^+^ blasts when exposed to an agent acting on the cellular epigenetic landscape, we concluded that clonal memory can be modulated.

## Discussion

Our study revealed that human HSPCs display two distinct forms of clonal memory— one for division and one for fate commitment. These clonal memories were consistently observed across multiple progenitor types, for more than two successive divisions, and diverse culture conditions, suggesting that clonal memory is a fundamental and conserved feature of early haematopoiesis. To the best of our knowledge, this is the first demonstration of both forms of clonal memory in human HSPCs. Clonal memory of differentiation has been previously described in murine haematopoiesis^52,53^, and we previously identified clonal memory of division in mouse^14^. They also align with indirect *in vivo* evidence of proliferation inheritance in twin patients developing the same neoplasms^54,55^ and in donor and recipient of bone marrow transplants ^20–22^. Although clonal memory of division has been reported in other cell systems^56–58^, such memory has not yet been described in differentiating systems.

As differentiation and division are linked, both factors could influence each other and produce synchronicity or similarity in fate within cells without clonal memory. Using our mathematical model, we demonstrated that clonal memory is required to explain the observed synchrony and similarity in fate. In fact, the presence of two distinct forms of memory for division and differentiation explained best the data, raising the possibility that they are governed by separate regulatory mechanisms. This echoes findings in B cell differentiation, where division and differentiation are independently clonally inherited^59^. Whether these two memory types are independent or coordinated remains an open question.

Interestingly, the division time of cells from the same family across generations did not correlate. One might think that it is not the division time itself that is transferred across generations, but the ability to divide at the same time. This mechanism was observed accross cell type, suggesting that it could be independent from commitment, but we do not exclude the fact that cells sharing the same differentiating phenotype could also share this mechanism. Advanced experimental approaches allowing to assess simultaneously the differentiation status and the division times^60^ will help elucidate this point.

We also identified that transcriptional fingerprints, gene modules shared by each member of that family, extend beyond differentiation states^43^, implicating clonal identity in shaping gene expression. Proliferative parameters such as clone size and division timing are poorly captured at the transcriptomic level, indicating that these features may be regulated through epigenetic or post-transcriptional mechanisms. Our results confirm that clonal analysis by transcriptomics^3,4,56–58,61^ is an effective method when the cells analysed display a strong clonal memory (cells from the same family display high transcriptional similarity). This should be confirmed beforehand, as we could observe a non-negligible fraction of discordant, more heterogeneous families, which could bias the clonal analysis by transcriptomics.

The existence of clonal memory in healthy haematopoiesis represents a form of non-genetic inheritance under non-pathological conditions. For this reason, we use the term *family* rather than *clone* to describe cell genealogies independently of the underlying cellular genetics. Our work focused on early haematopoiesis, the stage at which initial commitment decisions are made. Further studies are needed to determine how long clonal memory persists through subsequent stages of haematopoietic differentiation.

Functionally, clonal memory of division may serve to maintain proliferative heterogeneity among haematopoietic stem cells, a feature critical for homeostasis and stress-induced haematopoiesis. Such heterogeneity could enable distinct cell families to contribute to steady-state haematopoiesis versus responses to environmental fluctuations (e.g. inflammation, blood loss). Unexpectedly, we observed a marked reduction in clonal memory of division in AML, which could be partially restored by treatment with the epigenetic modifier JQ-1. This demonstrates that clonal memory is not only regulated but can also be tuned pharmacologically. While AML is composed of genetically distinct clones, our data reveal that proliferation rates vary even within genetically defined families. This intra-clonal variability may reflect a strategy by which leukemic cells reduce memory to enhance adaptability. Whether this reduction is a dynamic, reversible process or a fixed trait of selected clones remains to be determined.

The ability to modulate clonal memory pharmacologically opens new therapeutic avenues. By shifting the balance from plasticity toward stability, it may be possible to reduce intra-tumour heterogeneity and sensitize malignant cells to treatment. More broadly, our findings position clonal memory as a key regulatory layer in stem cell biology, with implications for both normal development and disease.

### Limitations of study

To enhance the reproducibility of our findings, we employed several culture conditions, including at least two distinct media and multiple timepoints, to ensure the reproducibility of our observations. However, future studies should aim to better capture the influence of the microenvironment, which we could only partially addressed here. We prioritized the analysis of many individual families per sample, particularly for the pathological ones, and the inclusion of a representative, yet incomplete selection of AML samples. We acknowledge the small number of samples displayed, which are not necessarily recapitulating the genetic and phenotypic heterogeneity associated to AML. The magnitude of the decrease in cell division concordance, or the JQ-1-associated increases, differed between samples, underlying different sensitivities to JQ-1 and unknown biological determinants associated to cell division synchronicity. Furthermore, our study faced the classical technical challenges with primary AML samples, notably: (1) the limited capacity for *ex vivo* expansion of primary leukaemia samples, especially in the absence of supportive stromal cells, and (2) the inherent constraints of single-cell culture systems. Continued development of advanced culture platforms will enable to expand our analysis of primary AML cell proliferation and differentiation.

## Material and Methods

### Ethical approval, banking, and processing of human samples

De-identified umbilical cord blood units were collected accordingly to the guidelines defined by the Saint-Louis Hospital Umbilical Cord Blood biobank (authorization AC-2016-2759) and in accordance with the Declaration of Helsinki.

Human Bone Marrow samples were collected accordingly to the guidelines defined by the Saint-Louis Hospital Institutional Review board (IRB 00003888), authorization 21-799.

Human AML primary samples (bone marrow or peripheral blood) were obtained from patients with informed consent for sample collection and use in research were obtained accordingly to the IRB-approved procedures from Institut Curie (Saint-Cloud, France), Gustave Roussy (Department of Clinical Haematology and Drug Development Department DITEP, Villejuif, France) and Hammersmith Hospital (London, UK). A summary of the patient samples used for this study can be found in **Extended Data Table 5**.

Mononuclear Cells (MNC) were isolated via Ficoll density gradient. Human bone marrow cells were systematically processed fresh, while umbilical cord blood units and human AML primary samples underwent CD34-enrichment using immunomagnetic microbeads (MACS Miltenyi). Total MNCs or enriched CD34^+^ fractions were immediately processed or frozen in Fetal Bovine Serum (FBS, Eurobio Scientific) supplemented with 10% dimethyl sulfoxide (DMSO, Sigma) for further analysis.

### Fluorescence Activated Cell-Sorting (FACS) and single-cell isolation

Total MNCs or enriched CD34^+^ fractions were prepared for immunostaining as previously described^17^. The list of antibodies used with the respective fluorochromes can be found in Extended Data Table 7. Single-cell FACS was performed using BD Aria III or BD Fusion instruments (Becton Dickson), in 96-wells U-bottom tissue culture vessels (Falcon). Cells were isolated using Single Cell sorting precision when individually isolated or Purity precision when isolated in bulk. The sorted cells were indexed for all the fluorescence parameters, using the Index sorting function, to record the fluorescence levels for each marker. Data analysis was performed using the software FlowJo v10.1 (BD Biosciences).

### MultiGen cell dye labelling

The cell dye labelling step was performed as previously described^17^. Briefly, cells were stained with four combinations of 5-(and 6)-carboxyfluorescein diacetate succinimidyl ester (CFSE, ThermoFisher Scientific) and CellTrace Violet (CTV, ThermoFisher Scientific): CFSE only, CFSEhigh + CTVlow, CFSElow + CTVhigh and CTV only.

For umbilical cord blood HSPCs, the labelling was performed in two sequential steps, starting with the CFSE staining and concluding with the CTV staining. The final concentration used for the labelling are respectively CFSE 5 µM (CFSE only and CFSEhigh CTVlow), CFSE 2.5 µM (CFSElow CTVhigh), CTV 5 µM (CTV only and CFSElow CTVhigh) and CTV 2.5 µM (CFSEhigh CTVlow).

For OCI_8227 cells, the labelling was performed in a single step, preparing 2X solutions for each of the cell dye combination. The final concentration used for the labelling are respectively CFSE 2.5 µM (CFSE only and CFSEhigh CTVlow), CFSE 1.25 µM (CFSElow CTVhigh), CTV 2.5 µM (CTV only and CFSElow CTVhigh) and CTV 1.25 µM (CFSEhigh CTVlow).

For primary human AML cells and healthy adult peripheral blood, the labelling was performed in a single step, preparing 2X solutions for each of the cell dye combination. The final concentration used for the labelling are respectively CFSE 5 µM (CFSE only and CFSEhigh CTVlow), CFSE 2.5 µM (CFSElow CTVhigh), CTV 5 µM (CTV only and CFSElow CTVhigh) and CTV 2.5 µM (CFSEhigh CTVlow).

### Short-term liquid culture of primary cells

Progenitors isolated from umbilical cord blood or healthy bone marrow, either as single cells or in bulk, have been cultured in two types of cell culture medium, named Gene Therapy medium (GT)^27^ and Differentiation medium (Diff)^26^. Both media are based on StemPro-34 SFM (SP34, ThermoFisher Scientific) supplemented with Penicillin – Streptomycin 1X (Pen/Strep, ThermoFisher Scientific), L-Glutamine 2 mM (ThermoFisher Scientific) and human Low-Density Lipoprotein 500 ng/mL (LDL, Stem Cells). For GT medium we added the following cytokines: SCF 300 ng/mL (Peprotech), FLT3-ligand 300 ng/mL (FLT3l, Peprotech), TPO 100 ng/mL (Peprotech), IL-3 60 ng/mL (Peprotech). For Diff medium: SCF 100 ng/mL, TPO 100 ng/mL, IL-6 50 ng/mL (Peprotech), GM-CSF (Miltenyi) 25 ng/mL, FLT3l 20 ng/mL, EPO 1 U/mL (Janssen) and IL-3 10 ng/mL. Progenitors isolated from AML samples, both as single cells orbulks, were cultured in a slightly modified version of the Diff medium, containing IL-1β 50 ng/mL (Peprotech). A complete list of references for cytokines and small molecules is available in **Extended Data Table 6**.

### MultiGen flow cytometry assessment of cell division, cell differentiation and cell family

Flow cytometry experiments, both using umbilical cord blood progenitors, healthy and leukemic peripheral blood and 8227 cells, were performed as previously described^17^. The list of antibodies used with the respective fluorochromes can be found in **Extended Data Table 7**. Data analysis was performed using the software FlowJo v10.1 (BD Biosciences) and custom Python scripts. The recovery rate was experimentally measured following the same protocol, except that each well was seeded with a range of CTV-stained 8227 cells (4 to 16 cells). Immediately before the flow cytometry analysis, each well was manually assessed and the number of light-refringent cells assessed via an inverted microscope. The recovery rate was therefore calculated as a ratio between the value from the flow cytometry analysis (CTV+ cells in the Single Cells gate) and the microscope value (Extended Data Figure 1c-d).

### Time-lapse imaging

Haematopoietic progenitors from umbilical cord blood or adult bone marrow were, as previously mentioned, individually sorted via FACS in 96-wells U bottom plates (Falcon) and then continuously imaged for 96 hours, using an Incucyte Zoom system (Sartorius). Cell culture was performed using the same culture media previously described. Time-lapse experiments were conducted at 37 °C, 5% CO_2_, with time intervals of bright field image acquisition of 1-2 hours. Cell viability and cell division were assessed manually: high refractility, membrane integrity and cell size were the criteria used to assess viability, and cell death time was measured as the first image when visible blebbing could be observed, together with cell fragmentation, loss of circularity and lack of cell movement. The first cell division time was measured as the time between the beginning of the experiment and the time point at which two cells could be clearly distinguished. For the second and third divisions, the time was measured as the interval between the end of the former division and the time point at which two novel cells could be clearly distinguished and were adjacent in the same region. We stopped to measure the time for cell division after the third division, due to cell motility making impossible to reliably track each individual cell manually. For the analysis of the synchrony in division for the second divisions, we randomised the order of annotation of the two division times, to avoid any bias in the two distributions.

### Cell line culture and *ex vivo* treatments

OCI_8227 cells were maintained in culture for no more than eight weeks at the time. Cells were passaged every 5 days in T-25 or T-75 cell culture-grade flasks, at a cell concentration of 1.5-3 x 10^5^ cells/mL. Cells were cultured in StemSpan II (Stem Cells) supplemented with Pen/Strep 1x, SCF 50 ng/mL, TPO 50 ng/mL, FLT3l 50 ng/mL, G-CSF 10 ng/mL, IL-6 10 ng/mL and IL-3 10 ng/mL. 8227 cells were treated *ex vivo* using sublethal doses of JQ-1 (Sigma): the working solutions were prepared accordingly to the manufacturer’s instructions.

### Confidence Intervals for MultiGen analysis

The confidence intervals at 95% level were calculated via basic bootstrap^62^ with 250,000 bootstrap datasets. Following this procedure, each bootstrap dataset was constructed by sampling with replacement as many cellular families as were in the original data. The distribution of the statistics, each calculated from one bootstrap dataset, then provided a reference from which the confidence interval was derived. Formally, given the statistics *ϑ*^ calculated from the original data, and *ϑ*\*_(0.025)_ and *ϑ*\*_(0.975)_ percentiles, respectively derived from the bootstrapped distribution, the confidence interval of *ϑ*^ at 95% level was calculated as follows:

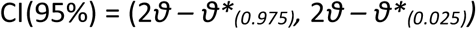

### Cell division range analysis

To quantify how correlated were the cell decisions to divide, for cells from the same generation and descending from the same ancestor, we employed a previously used stochastic model^14,18,19^, upgraded to account for cell death. Biologically, each cell reaching a certain generation have a chance of surviving that is independent from other cells. All surviving cells can then divide or stop dividing, with a given probability that is correlated between cells from the same family and generation. When sampling, cells that survive but divide less tend to be also less detected, simply because they are less abundant. Thus, it is important to take both survival and sampling into account when measuring the range of division within a family. In this model, cells have a certain probability to survive before dividing, as defined by the survival rate *r_surv_*. Cells have also a certain probability to be measured, defined by the recovery rate *r*. The correlation coefficient that links the cell division within the same family was fitted to the data and was used to evaluate the degree of correlation in division within families, otherwise called concordance in division. The model was directly parameterized by the data, apart from one variable that encapsulated the correlation in decision-making that fitted to the data. With *n* being the maximum generation recorded, let p_i_ ∈ [0, 1] for *i* = 0,…, *n* denoted the proportion of cells that divided from generation *i* to the next, which was determined from the data as follows. Set z_i_ the total number of cells recovered in generation *I*, the p_i_ was estimated by

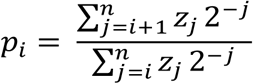

For *i* = 0,…, *n*-1 and by *p^_n_* = 0.

In this model, given *k_i_* the number of cells from a particular family that reach generation *i*, the number of cells that continued dividing to *i* + 1 followed a beta-binomial distribution with a number of trials that was a random variable *K_i_* (independent of all the others) and parameters *a_i_ = p_i_ (1 − ρ)/ρ*, and *b_i_ = (1 − p_i_)(1 − ρ)/ρ,* namely *β(K_i_, a_i_, b_i_),* where *ρ* ∈ [0, 1] was a free parameter.

To account for cell death, *K_i_* = B(*k_i_, r_surv_*) followed a binomial distribution with k_i_ that reached generation *i* and survived to divide. *r_surv_* was estimated from time-lapse microscopy experiments for each cell type and specified in the results section. Finally, one family was generated recursively by setting k_o_ = 1 and defining *k_i+1_ =* 2*β(K_i_, a_i_, b_i_)*. The recovery rate *r* was determined experimentally as previously described, and set at 0.6. In the model, we accounted therefore for the sampling effect by sub-sampling each cell from every artificially generated family with a probability of 0.6 and independently of all other cells. The beta-binomial model interpolated between surviving cells that decided to divide independently of each other (*ρ* = 0) and cells that were perfectly aligned in their decision to divide (*ρ* = 1). A value between 0 and 1 reflected the concordance level within each family in division, but, by design irrespective of the values of *p_i_*, that determined the population distribution among generations. Defining the family range as the difference between maximum and minimum generation for all recovered surviving cells per family, the best fit *p^* of *p* was determined to be the value from 0 to 1 that maximized the likelihood of recapitulating the family range distribution in the data. The 95% confidence interval for *p^* was calculated following the basic bootstrap procedure previously described, that was by sampling with replacement the cellular families and setting the maximum-likelihood estimation as *p^* the statistic to be calculated on the bootstrapped datasets. To assess the significance of *p^*, the statistical hypothesis H_0_ : ρ = 0 was tested against the alternative H_1_ : 0 < ρ ≤ 1 using the likelihood-ratio statistic

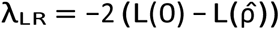

where *L*(*ρ*) was the log-likelihood of the cellular family ranges under the beta-binomial model with parameter *p*. The p-value for the test was derived from the asymptotic distribution of λ_LR_ that was a χ^2^ with 1 degree of freedom.

### Mathematical modelling and statistical inference of the HSPC proliferation and differentiation dynamics

The detailed description of the mathematical model, the parameter estimation and model selection procedure, and supplementary results are available in the Supplementary Information (SI).

#### Model assumptions

we considered a stochastic model, at the scale of a family, continuous in time (denoted by *t*), over a short period (*T*_*max*_ = 96h), and where each cell was characterized by a phenotype *p*, with four different possibilities: *p* ∈ 𝒫 = (CD90^+^, CD90^-^, HPC, CD34^-^), where 𝒫 was naturally provided with an order relation (e.g., a CD90^-^ being more differentiated than a CD90^+^). Therefore, we confounded 𝒫 with the set [1,2,3,4]. *p* = 3 would then correspond to an HPC. At *t* = 0, the process begun with a single cell of a known phenotype *p*_1_ **(Fig. 3a)**. The initial cell division occurred at a time T_1_, randomly drawn from the empirical distribution of observed first division times (of cells of type *p*_1_) (for further details, see **SI – B.3.1**). We treated separately the first division from the subsequent ones since we observed that the initial division typically exhibits a longer duration **(Extended Data Figure 2j-l)**. After division, two daughter cells were considered. No cell death was considered in our model, and each of the daughter cells was assigned a cell phenotype at least as differentiated as the mother cell (we assume no de-differentiation). The probability for a daughter cell to be of type *p = i*, given its mother’s type *p_m_ = a*, was denoted by *p*_*a*→*i*_ = ℙ[*p* = *i* | *p*_*m*_ = *a*]. Then, the cells pursued the proliferation process, with division times randomly sampled from Log-Normal distributions (ℒ𝒩) where the distribution parameters (*μ*_*p*_, *σ*_*p*_) varied according to the dividing cell’s phenotype: *T* ∼ℒ𝒩(*μ*_*p*_, *σ*_*p*_). We noted that a Gamma distribution would have been appropriate as well, as we describe in Supplementary Material B.3.2.

#### Possible scenarios

The generic model of HSPC proliferation and differentiation could be declined into several scenarios to explore different biological hypotheses **(Fig. 3b)**. These scenarios modify the interactions between sister cells, to reflect distinct biological hypotheses that we evaluated for their agreement with our experimental data. Scenarios were defined based on two criteria: homogeneity in fate (H) and concordance in division timing (C). Homogeneity (H) assesses whether sister cells tended to share the same fate (H=1) or not (H=0). Concordance (C) evaluated whether sister cells tended to divide simultaneously, with C = 1 for synchronised divisions and C = 0 for independent timing. Each scenario was identified by a pair of values for (H, C), creating a matrix of possibilities. The default scenario, characterized by H = 0 and C = 0, posited that sister cells were fully independent from each other (conditioning on their mother). This scenario served as a baseline, against which the implications of fate homogeneity and division concordance were assessed. When we hypothesized a homogeneity in fate (H = 1), we considered in our model a higher probability to have a symmetrical division compared to the scenario where H = 0. To do that, we introduced the parameter *ρ*_*H*_ ∈ [0,1] such that the probability of having two daughter cells of the same type *b*, given their mother’s type *a*, was equal to: 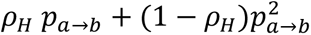. In the limiting case where *ρ*_*H*_ = 1, all divisions were symmetrical. The case *ρ*_*H*_ = 0 corresponded to the hypothesis H = 0 of absence of homogeneity in fate. When we hypothesised a concordance in division (C = 1), we considered that the division times of two sister cells were not independent anymore but correlated. Let us denote by T_i_ and T_j_ the division times of two daughter cells of type p_i_ and p_j_, respectively. Then, in our model, the division times of the two sister cells followed a bivariate log-normal distribution:

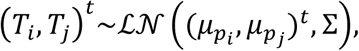

where the covariance matrix was expressed by: 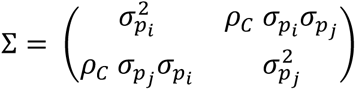

With parameter *ρ*_*C*_ ∈ [0,1] quantifying to which extent sister cells were concordant.

The limiting case *ρ*_*C*_ = 0 would correspond to the hypothesis C = 0, which was absence of concordance in division.

In addition to our four main scenarios, we also studied more complex ones, by considering that *ρH* would depend on the mother cell type (SI – B.2.3), or that there could be concordance in division between cousin cells, or that the concordance between sister cells would depend on their types (SI – B.3.4). We also tested a minimal – more parsimonious – scenario, still without concordance in division timing and without homogeneity in fate, but also without differentiating the distribution of the division time, according to the type of cell which divides. That is, in this minimal scenario, we assume that *μ*_4_ = *μ*_3_ = *μ*_2_ = *μ*_1_ and *σ*_4_ = *σ*_3_ = *σ*_2_ = *σ*_1_.

We denote by:

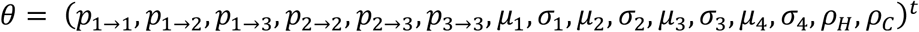

our parameter vector.

#### Comparing the model simulation with the experimental observations

To evaluate which scenario is the most likely, we compared the output of our model simulations to the experimental observations, defining a distance between the data and simulations. We denoted by 𝒟 our dataset (for further details about its definition, the inclusion and pre-treatment step, and the uncertainties in the observations, see **Supplemental Information – A**). 𝒟 is of size 1,740, meaning that we had the observations for 1740 different families, either from the MultiGen or the single-cell live imaging assay, for different initial conditions, and for different observation times. Because of the large size of our dataset, and the heterogeneity in the observations, defining a proper distance was challenging. Our methodology involved the derivation of an extensive set of summary statistics, denoted as S_i_, where *i* indexes the specific summary statistic within the set. Each summary statistic can be computed from experimental observations, in that case we noted: *Ŝ*_*i*_ = *S*_*i*_(𝒟).

Given a parameter vector θ, we simulated the model to generate a virtual dataset of equivalent size to the number of experimental observations, from which we calculated each theoretical (i.e., given by our model) summary statistic *S*_*i*_(*θ*). Summary statistics synthesised the information from the observations (or from the simulations) making comparisons possible. This was accompanied by a potential loss of information, which we first minimised by defining 340 summary statistics. Ideally, we aimed to find parameter values such that the simulations of the model lead to computed summary statistics as close as possible to the experimental ones, that is, find *θ*^∗^ such that, ∀*i* ∈ {1,…, 340}, *S*_*i*_(*θ*^∗^) ≈ *Ŝ*_*i*_.

However, given the number of summary statistics considered, and the potential redundancy and high correlation among them, we refined our approach using Principal Component Analysis (PCA), thus reducing the dimensionality of our observation space. We ended-up with 20 principal components, accounting for 80% of the variance, denoted as 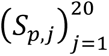 where each *S*_*p*,*j*_ was a linear combination of the original 340 summary statistics.

The distance d(θ) was then defined as a mean squared error (MSE) between the simulated and observed values of these principal summary statistics:

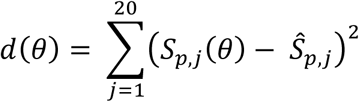

This approach provided a practical balance between simulation accuracy and computational cost, making it well-suited for methods like Approximate Bayesian Computation (ABC).

#### The ABC method

to compare the experimental observations to the output of our model, through the construction of a relevant distance d(θ), we ran the ABC method to determine the posterior probability of each scenario, given our experimental data. First, we defined our prior distributions. We considered *a priori* our four scenarios to be equiprobable. Then, we defined the prior distributions of our model parameters. Except for *ρ*_*C*_ and *ρ*_*H*_, all the parameters did not depend, *a priori*, on the scenario. We considered them uniformly distributed (see **Supplemental Information – B.1 and B.4** for further information on the possible range of values of the different parameters). For our parameter associated with the hypothesis of homogeneity in fate, we had: ℙ[*ρ*_*H*_ = 0 |*H* = 0] = 1 and *ρ*_*H*_| *H* = 1 ∼ 𝒰([0.3, 1]) (𝒰 denotes a uniform distribution) such that we enforced a clear distinction between our scenarios of absence (H=0) or presence (H=1) of homogeneity in fate. We did similarly for *ρ*_*C*_. In a Bayesian framework, the objective was to update our prior, through the integration of the data, to get the posterior distribution. Given the complexity of our model, the likelihood was intractable, hence the need to use an approximate Bayesian Computation method where the posterior distribution might be approximated. Instead of using the expression of a likelihood, we relied on the definition of our distance d(θ) between the model and the observations.

#### The ABC-rejection sampling method

For *i* ∈ {1, ⋯, *N*}, we sampled *a* modelling scenario (H_i_, C_i_) and *a* parameter vector θ_i_ from the prior distributions. We ran as many simulations as observations (1,740), computed the principal summary statistics S_p,i_(θ_i_) and then the distance to the observation d_i_ = d(θ_i_). We kept only the samples *i*^′^ ∈ *I* such that d_i’_ < Ɛ, where Ɛ corresponded to the 0.1 percentile computed over the (d_i_)_1 ≤ i ≤ N_. That is, we kept the 0.1% best samples. These samples were used to approximate our posterior distribution **(Extended Data Fig. 10a)**. Finally, the posterior distribution of each of our four scenarios – when using the ABC-rejection algorithm - was approximated by:

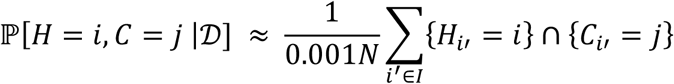

That is, we counted how many times we selected each scenario among the 0.1% best samples. With N=1,733,106, the results of this procedure led us to select the scenario (H=1, C=1) as the most likely **(Supplemental Information - D.1)**. Using the ABC-rejection sampling algorithm to explore how more complex modelling scenarios performed, and testing nearly 11 million of different parametrisations, we found no evidence that more complex scenarios would be more appropriate to describe our experimental observations **(Supplemental Information – E.1-3)**. In addition to the four main scenarios, we considered a minimal scenario with a single time-to-division distribution (i.e., without distinguishing between cell types). Running ∼500,000 simulations under this scenario (this number was chosen to consider a priori all five scenarios equiprobable) has almost no impact on the previous results. Despite involving six fewer parameters, this minimal scenario reaches a very low posterior probability (0.0152) after ABC-rejection sampling and can therefore be discarded **(Supplementary Information - E.4).**

#### The ABC-SMC method

The ABC rejection algorithm requires to sample parameter vectors a high number of times to converge. Then, to refine our results, and ensure their robustness and the convergence of the algorithm, we implemented the ABC-SMC sampler from Toni and colleagues^63^ (details of the implementation in **Supplemental Information - D.2**). With this method, we considered N = 2000 parameter vectors called particles, to approximate the posterior distribution. The ABC-SMC algorithm is sequential, so we iteratively sampled the N particles from intermediate distributions, starting from the prior distribution to the posterior distribution. At each iteration i, we sampled a parameter vector from the previous intermediate distribution (sampling the previously selected particles, according to some weights, and using pre-defined perturbation kernels) and ran the simulation to compute the distance to the observations. As for the ABC-rejection algorithm, particles were accepted if they are below a given threshold, discarded otherwise. In the ABC-SMC method, there was a decreasing sequence of tolerances. The ABC-SMC method ran for many iterations, until convergence, that was, until the distributions of the distances did not evolve for new iterations **(Extended Data Fig. 10b)**. Over the iterations, those distributions evolved until reaching a stable value, which corresponded to the approximated posterior distributions of the modelling scenarios **(Extended Data Fig. 10c).** To ensure the robustness of the ABC-SMC algorithm that we used, also regarding the choice of the perturbation kernels used, we ran different configurations, leading to the same results **(Extended Data Fig. 10e)**. ABC-SMC approach refined the results from the ABC-rejection sampling approach, from iteration 9, allowing to better explore the parameter space, progressively focusing on the region where the parameter vectors were more likely to have generated the experimental data.

We focused on the discrimination between modelling scenarios, rather than the estimation of the parameters of the most likely model. Yet, we also got the intermediate posterior distributions of the model parameters and saw how each iteration allowed to be more informative on the different parameter values, and how the parameters for which the model converged faster than other ones **(Extended Data Fig. 10f** and **Supplemental Information - D.4**). Some parameters stayed undetermined (or weakly determined) at this stage, meaning that their posteriors did not move away from the prior. When comparing the summary statistics computed from the models – sampling from the posterior – to the experimental values, we observed for most of them already a good agreement **(Extended Data Fig. 10d** and **Supplemental Information - D.5**)

### CITE-seq profiling of short-term liquid culture

#### Antibody labelling

magnetic sorted CD34^+^ cells from umbilical cord blood were plated at a cell density of 1×10^5^ cells/mL, using the same culture condition described for the condition Diff. Cells were harvested after 96 hours, counted with a haemocytometer, and resuspended in cold PBS 1X (ThermoFisher Scientific) containing 2% Bovine Serum Albumin (BSA, Euromedex) and 0.01% Tween 20 (Sigma), to a final concentration of 10^6^ cells/mL. 100 µL of this cell suspension was incubated with a mix of previously prepared CITE-seq antibodies. Cells were incubated for 30 minutes at 4 °C, washed three times with the same solution used for the incubation and resuspended in 16.5 µL, at a final concentration of 1000 cells/mL.

#### Library sequencing

approximately 1.65×10^4^ cells were loaded, to obtain a final yield of 1×10^4^ recovered cells. Cells were processed accordingly to the Chromium Single Cell 3’ library v3.1 (10x Genomics). Libraries were sequenced using a Novaseq s1 (Illumina), allocating at least 20,000 reads per cell for the mRNA library, and 5,000 reads for the Antibody Derived Tags (ADT) library **(Extended Data Table 8)**.

#### Computation methods

Raw sequencing reads were processed using Cellranger v3.1.0. To obtain a reads/cell/gene count table, reads were mapped to the human GRCh38 reference genome. Cells with less than 1000 genes per cell and with a high percentage (> 30%) of mitochondrial genes were removed from downstream analyses. Following our filtering procedures, the average UMI count per cell was 4406. The median number of genes detected per cell was 1662, 2.2% mapped to mitochondrial genes. Data normalization and integration were performed using the default Seurat v4 approach FindIntegrationAnchors() followed by IntegrateData(), and differentially expressed genes were determined using a logistic regression in Seurat on the non-integrated data using the FindConservedMarkers() function. Pathway based analyses were performed using the enrichR package. Unsupervised clustering was performed on the significant variable genes using the 10 first PCA followed by the nonlinear dimensionality reduction technique UMAP. Annotation of the data was obtained by mapping published signatures using the AddModuleScore() method of Seurat.

### Single-cell re-distribution

Single cell distribution of individual cell families was performed using the cellenONE X1 platform (Cellenion), in two independent experiments. Briefly, we followed the procedure describe in the section “Time-lapse imaging”, for three tissue culture plates per experiment. Then, we randomly selected individual wells displaying at least four cells from the three 96-wells plates (10 families per plate). We harvested the content of the entire well, centrifugated for 8’ at 300g and resuspended in 6 µL of PBS 1X (Sigma). Then, the whole volume was loaded into the machine, but due to the dead volume, effectively only 4 µL of this cell suspension could be redistributed. Individual cells were each sorted directly into single wells of 96-wells plates (90 cells per plate) at 4 °C, each well containing 3 µL of lysis buffer. The lysis buffer contains 0.03 µL Triton X100 (ThermoFisher Scientific), 0.0375 µL Recombinant RNAse Inhibitor (Takara Bio), 0.2 µL dNTP 10mM (ThermoFisher Scientific), 0.5 µL PEG 40% (Sigma), 0.04 µL SMARTseq3 oligo-dT 100 μM (/5BiosG/ACGAGCATCAGCAGCATACGATTTTTTTTTTTTTTTTTTTTTTTTTTTTTTVN; IDT) and ultrapure RNAse water (Invitrogen).

### SMART-Seq3 library preparation and sequencing

Reverse transcription (RT) and the subsequent steps were performed accordingly to the SMART-seq3 technique^41^. Briefly, lysates in each well underwent RT (42 °C for 90 mins, 10 cycles at 50 °C 2 mins + 42 °C 2 mins, and finally 85 °C for 5 mins) on ice, in presence of template switch oligonucleotide at 100 µM (/5BiosG/AGAGACAGATTGCGCAATGNNNNNNNNrGrGrGIDT), 0.04 µL Maxima H minus reverse transcriptase (ThermoFisher Scientific), 0.1 µl TRIS-HCl 1M PH 8.3 (Invitrogen), 0.12µL NaCl 1M (Invitrogen), 0.04 µL GTP 100mM (ThermoFisher Scientific), 0,32 µL DTT 100mM (ThermoFisher Scientific), 0.2 µL MgCl2 50mM (ThermoFisher Scientific) and 0.05 µL Recombinant RNAse Inhibitor (Takara Bio) and water. The following PCR step was performed immediately after, using 5 µL Kapa Hifi HotStart Ready Mix (Roche), and forward (100 µM TCGTCGGCAGCGTCAGATGTGTATAAGAGACAGATTGCGCAATG) and reverse (25 µM ACGAGCATCAGCAGCATACGA) primers (IDT technologies) for 25 cycles. After DNA purification using AMPure XP microbeads (ratio 0.8:1) (Beckman Coulter, A63881), the cDNA was tagmented in a final volume of 2 µL, using the Nextera XT DNA Library preparation kit (Illumina). The following library amplification was performed adding 1.5 µL of UDI primers and 2,5 µL of PCR mix including the Phusion High Fidelity DNA polymerase (ThermoFisher Scientific). After tagmentation, the different samples were pooled, purified using AMPure XP microbeads (ratio 0.9:1) and sequenced at 100-bp paired end on a high-output flow cell, using a NovaSeq SP, 800M reads (Illumina).

### Single-cell 3’ mRNA sequencing pre-processing

FASTQ files for each single cell were generated using bcl2fastq (version 2.20.0), with default parameters. The Read 1 contain the TSO (constant tag sequence and a UMI) followed by cDNA or only cDNA. The Read 2 contain cDNA only. Index1 and index2 reads correspond to cell-specific barcode. to generate UMI count tables from FASTQ files, the scRNA-SmartSeq3 pipeline v1.2.1 (https://doi.org/10.5281/zenodo.15056215) was used. Briefly, reads were aligned to the human genome (hg38) using STAR (version 2.7.8a) and assigned to genes using FeatureCounts (deeptools version 3.5.0). UMIs were extracted, deduplicated and counted for each gene using umiTools (version 1.1.1). Low-quality cells were filtered out based on a minimum of 500 genes detected and a percentage of mitochondrial RNA lower than 20. Families composed of only one cell were also removed.

### Dimensionality reduction, clustering and pseudotime analysis

Cell fate was determined based on known marker gene expression. For each dataset the Seurat workflow is implemented by applying SCT normalization, PCA dimensionality reduction, clustering cells and running UMAP reduction [1]. The cells cluster in three groups, namely CD34+CD38-/HPC, MyP (Myeloid Progenitors) and EryP/MkP (Erythroid/Megakaryocyte Progenitors). The ELANE-expressing cells are labelled as GraP (Granulocyte Progenitors). The pseudotime analysis was performed using the Slingshot package, specifying the cluster CD34⁺CD38⁻/HPC as the starting cluster. For cells assigned multiple pseudotime values across different lineages, the maximum value was retained for each cell.

### Cosine distance

The variable “distance” was a measure of how cells of the same family tended to be transcriptomically similar, regardless of their cell fate. The cosine distance between two cells was defined as follows^42^:

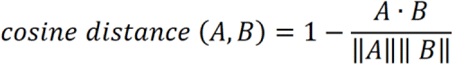

where A represented the gene expression vector of the first cell and B represented the gene expression vector of the second cell. For each family, the cosine distance was computed for each pair of cells and the median was assigned. The cosine distance distributions were computed on the log-normalized UMIs for real families and both types of random families.

### Identification of Variably Expressed Genes Across Families

Single-cell RNA sequencing (scRNA-seq) data from Smart-seq3 was analysed to investigate gene expression variability across families. The pre-processed and normalized data were used for the analysis.

Cells were grouped into families based on well information. Families with fewer than two cells were excluded. An analysis of variance (ANOVA) was performed for each gene to assess differential expression across families:

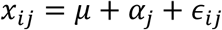

where *x*_*ij*_ represents the expression of gene ii in family *j*, *μ* is the overall mean expression, *α*_*j*_ represents family-specific effects, and *∊*_*ij*_ denotes residual error.

The total variance was decomposed into between-family variance 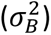 and within-family variance 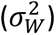, and an F-test was applied to determine whether the between-group variance was significantly greater than the within-group variance:

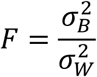

where a high F-value indicates greater variance between families than within families. To control for multiple hypothesis testing, p-values were adjusted using the Benjamini-Hochberg procedure to limit the false discovery rate (FDR). Genes with adjusted p-values (FDR < 0.05) were considered significantly homogeneous within families. To visualize gene expression patterns across families and clusters, a heatmap was generated using pheatmap R package. The expression matrix was prepared by including the significantly family conserved genes. The samples were ordered by family information, while genes were clustered separately for cells and genes using hierarchical clustering with Euclidean distance and the complete linkage method. The genes were clustered into modules by cutting the hierarchical cluster tree.

### MIIC analysis

MIIC was used to investigate which genes were significantly associated with specific families, their recorded division times, colony size, predicted cell types, and transcriptomic dispersion “distance”, using transcriptomics data of 142 cells obtained via SMART-Seq3^41^. In the reconstructed graphical networks, the variables were represented as nodes and the edges encoded for dependency. An edge with an arrow stands for causality, an edge without an arrow stands for association, a bi-directed dotted edge represented an unobserved common cause between two connected variables^45^. The green arrows represented genuine causality while the non-green arrows represented putative causality. In the case of putative causality, it was not possible to exclude the presence of an unobserved common cause. The link-colours between two nodes were assigned based on the Spearman’s correlation coefficient: red stand for positive correlation and blue for negative correlation. Gray was used for categorical variables without natural order, such as the family IDs. The networks were reconstructed from two scRNA-seq datasets, corresponding to two different experiments: 70 cells belonging to 14 families and 28,750 genes for the first, 72 cells belonging to 16 families and 30,682 sequenced genes for the second. The metadata for each dataset was the following: family of origin, times of division, colony size at 90 hours. The network was reconstructed from the 142 cells with their respective raw UMI counts of selected genes and additional features of interest. The gene selection was performed based on their high mutual information with the cellular features of interest: “family”, “T1”, “avg dT2”, “avg dT3”, “predicted.celltype”, “Colony size”, “exp”, “distance”. The genes selected to reconstruct the networks shared the highest mutual information with these properties of interest (i.e. we retained all genes above some tuneable mutual information threshold). Mitochondrial, Ribosomal Protein Large subunit (RPL), Ribosomal Protein Small subunit (RPS), Large Subunit Mitochondrial Ribosomal Proteins (MRPL) and Small Subunit Mitochondrial Ribosomal Proteins (MRPS) genes were filtered out from the analysis. A confidence parameter of 1% was used. This parameter sets the probability upper bound that this edge could be spurious due to sampling noise. Hence, the confidence parameter allowed to filter the least robust edges^44^.

Two networks were obtained either including or excluding the “Family” variable:

- A network with the “family” variable and a total of 1009 variables (including 147 regulatory genes or variables and 862 consequence genes): https://miic.curie.fr/showcase.php?id=Donada_wtfamily
- A network without the “family” variable and a total of 716 variables (including 113 regulatory genes or variables and 603 consequence genes): https://miic.curie.fr/showcase.php?id=Donada_wofamily

### Statistical analysis

Statistical analyses are detailed in figure legends and performed using R (version 4.3) or Python (version 3.10). The number of independent replicates, experiments and donors is stated for each figure in the legend.

## Supporting information

Supplementary material with legends

## Data availability

Raw and pre-processed single-cell RNA-sequencing dataset are deposited here: CITE-Seq: https://zenodo.org/records/15720816

SMART-Seq3: https://zenodo.org/records/15721710 - https://zenodo.org/records/15722093 - https://zenodo.org/records/15722300 - https://zenodo.org/records/15722418 - https://zenodo.org/records/15722505

Additional data for reanalysis and reproducibility effort is available upon request to the lead author.

## Code availability

All the scripts used in this paper are freely accessible on GitHub:

*Live-cell imaging analysis:* https://github.com/TeamPerie/Live-cell_imaging.git

*MultiGen analysis:* https://github.com/TeamPerie/MultiGen.git

*Mathematical model of division and differentiation:* https://gitlab-research.centralesupelec.fr/2012hermangeg/HSPC_proliferation

*CITE-seq analysis:* https://github.com/TeamPerie/CITEseq_analysis_pipeline.git

*Smartseq3 pre-processing:*

https://github.com/bioinfo-pf-curie/scRNA-SmartSeq3/releases/tag/v1.2.1

*Smartseq3 quality control:* https://github.com/TeamPerie/smartseq3_familly_analysis

*Single-cell 3’ mRNA sequencing pre-processing and analysis:*

https://github.com/tizianatocci/Donada2025

https://github.com/TeamPerie/Donada-et-al2025-family-ANOVA.git

## Acknowledgments

We would like to thank all the members of the Perié lab for helpful discussion, particularly Anais Lamoureux, Jason Cosgrove, Lucie Hustin. We would like to thank the members of the Institut Curie Flow Facility for their help with setting up the flow cytometry experiments. We thank the members of the Institut Curie Imaging Platform for their assistance with the live-cell imaging experiments. We thank the Saint Louis hospital cord blood biobank for providing the biological resources necessary for this project. This project has received funding from the European Union’s Horizon 2020 research and innovation programme under the Marie Skłodowska-Curie grant agreement No 847718 and from Fondation pour la Rercherche Médicale (FRM) under the “Fin de thèse ARC FRM” programme (dossier No. ARC202506021018). The study was supported by grants from the Labex CelTisPhyBio (ANR-10-LBX-0038) (to L.P. and A.D.), the Canceropole INCA Emergence (2021-1- EMERG-54b-ICR-1, to L.P.), and the ITMO MIIC grant (21CM044, to L.P. and P.H.C.). As well as funding from the European Research Council (ERC) under the European Union’s Horizon 2020 research and innovation program ERC StG 758170-Microbar (to L.P.), A.D. was supported by a fellowship from Fondation de France (00119150 / WB-2021-36197).

## Authors Contributions

A.D. and A.M. performed, analysed and interpreted the live-cell imaging data. A.D., I.M. and C.P. performed, analysed and interpreted the MultiGen assay. G.P. developed the statistical framework for the MultiGen analysis, supervised directly by K.D. and with feedback from A.D. and L.P. G.H. developed the mathematical framework, under the supervision of P.H.C. and with feedback from A.D. and L.P. A.D., S.T.B. and D.D. performed the transcriptomics experiments. L.H.A., W.S. and T.T. analysed and interpreted the transcriptomics data, supervised directly by H.I. and L.P. with feedback from N.S. and A.D. T.T. performed and interpreted the network analysis, supervised directly by H.I. A.I., C.W., J.V., D.M., C.P. and C.L.C. provided the biological and clinical samples. A.D. prepared the paper final figures, with contributions from T.T., A.M. and W.S. A.D. and L.P. wrote the manuscript, with substantial contribution from L.L. and all the co-authors.

## Declaration of interests

D.M. received research funding from Novartis, Sanofi and CSL Behring, and consulting fee from Incyte, Novartis, Sanofi, CSL Behring, Jazz pharmaceuticals and Mallinckrodt. None of this COI are directly linked to this research.

The other authors have nothing to declare.

## Notes

### Summary of Updates

Revised version with new experiments, revised models and revised text

