## Supplementary material with legends for "Clonal memory of cell division in humans diverges between healthy haematopoiesis and acute myeloid leukaemia"

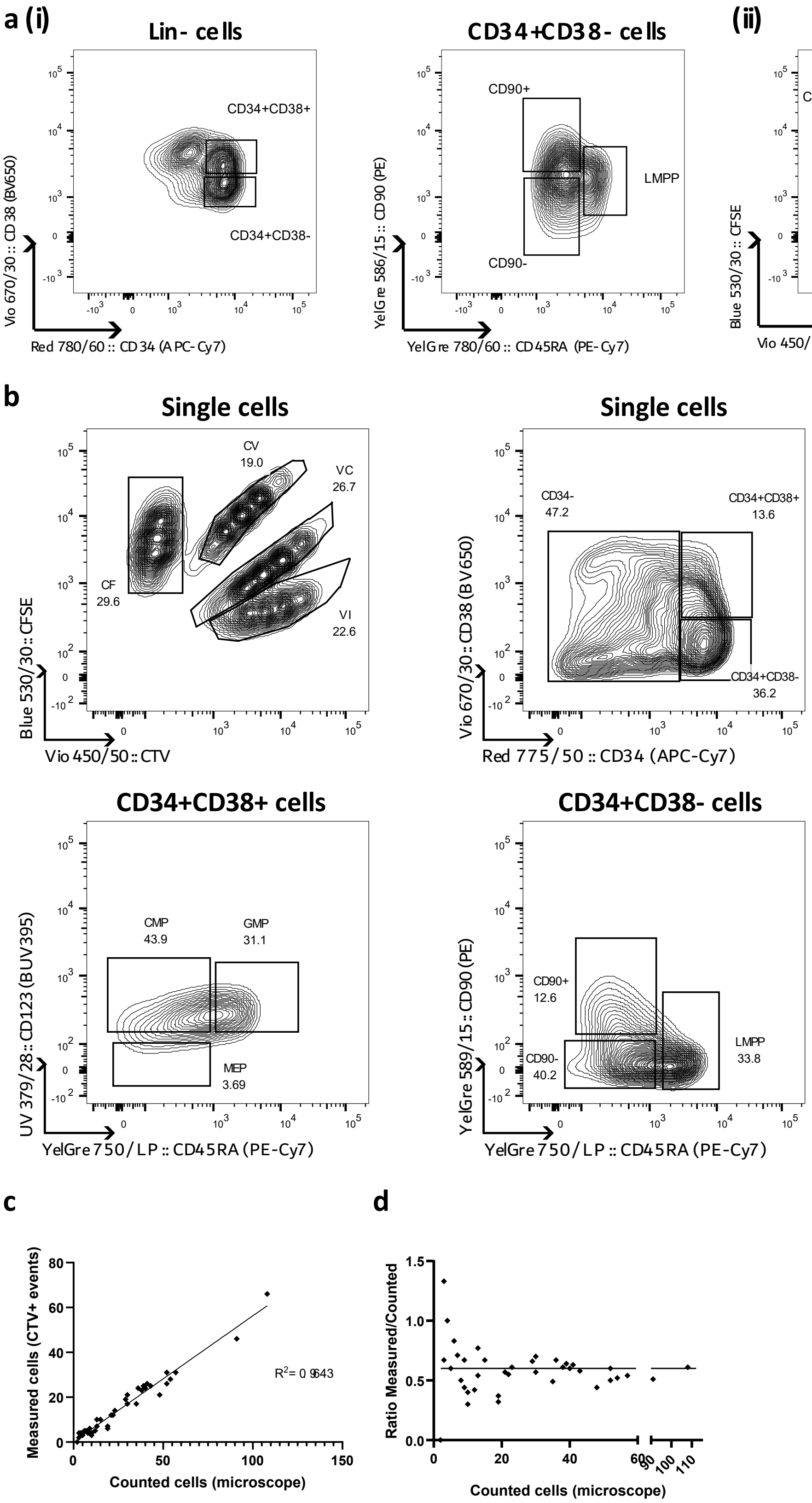

***Extended Data Figure 1: Immunophenotypic separation of isolated and ex vivo expanded HSPCs.*** ***a)*** *Representative plots of FACS sorting of hematopoietic stem cells (CD90⁺s), multipotent progenitors (CD90⁻s) and hematopoietic progenitor cells (HPCs) from isolated mononuclear cells of human cord blood. The same gating strategy* ***(i)*** *was used for bone marrow and peripheral blood analyzed via live-cell imaging. For* ***(ii)*** *MultiGen staining, CTV and CFSE were added accordingly to protocol.* ***b)*** *Gating strategy of ex vivo expanded HSPCs, experimentally prepared accordingly to the MultiGen protocol. CMP, common multipotent progenitor; LMPP, lymphoid-primed multipotent progenitor; MEP, megakaryocyte–erythroid progenitor; GMP, granulocyte-monocyte progenitors.* ***c)*** *Correlation between the number of counted cells through live imaging and the recovered cells, measured as CTV+ events in the single cells gate. Pearson correlation coefficient R, n = 41, two independent experiments.* ***d)*** *Correlation between the recovery rate, calculated as a ratio between measured and counted cells, and counted cells. The horizontal line corresponds to the recovery rate of 0.6 used in the modeling.*

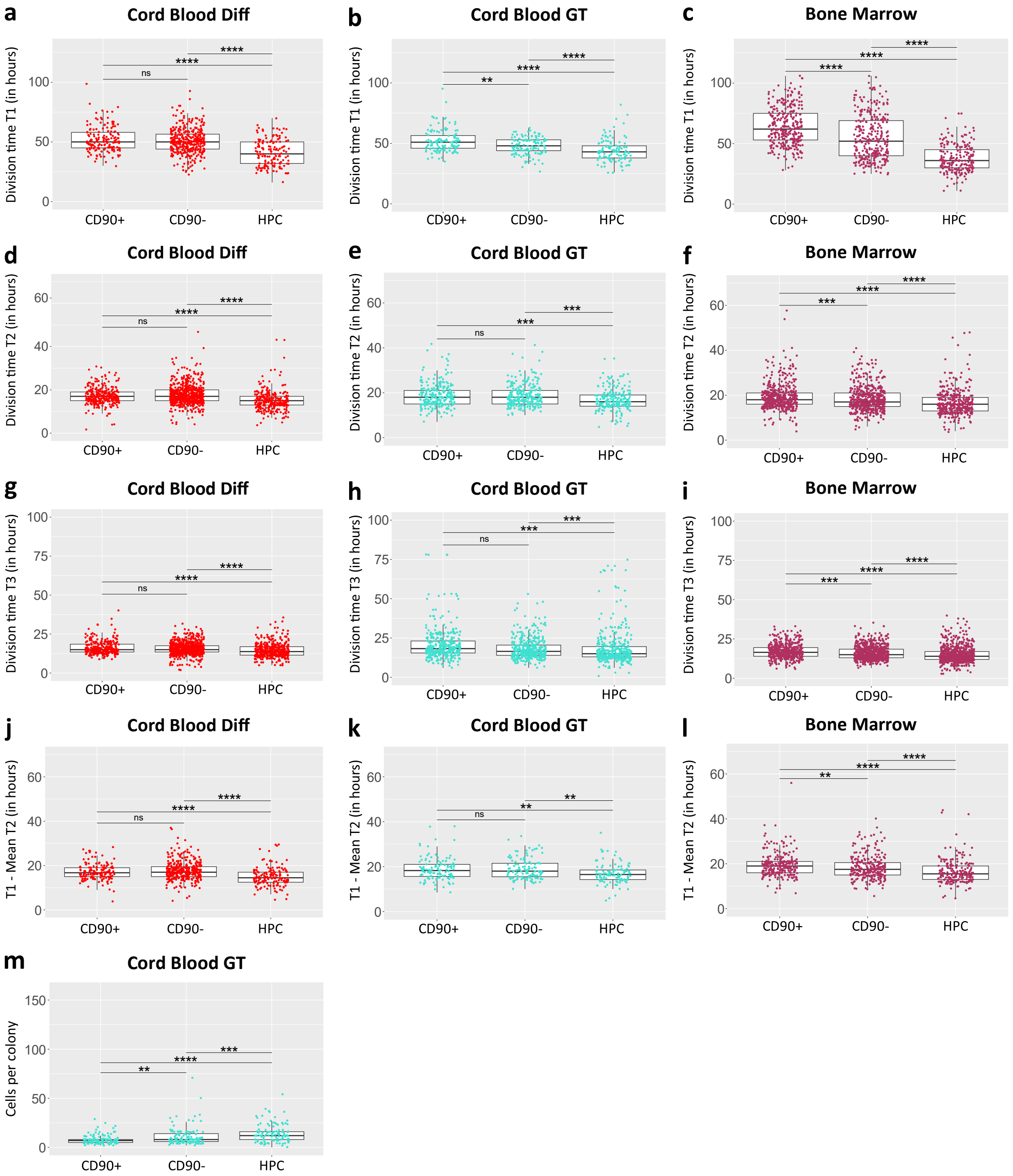


***Extended Data Figure 2: Cord blood and bone marrow HSPCs exhibit consistent divisional kinetics across sources and culture conditions.*** ***a)*** *Time necessary for completing the first division, for foetal cord blood HSPCs expanded in Diff medium. 7 independent experiments, from 7 different donors, n = 669 divisional events (159 CD90*$^+$*s, 363 CD90*$^-$*s and 147 HPCs).* ***b)*** *Same as* ***a),*** *for foetal cord blood HSPCs expanded in GT medium. 2 independent experiments, from 2 different donors, n = 370 divisional events (127 CD90*$^+$*s, 130 CD90*$^-$*s, 113 HPCs).* ***c)*** *Same as* ***a),*** *for bone marrow HSPCs expanded in Diff medium. 8 independent experiments, from 8 different donors, n = 846 divisional events (326 for CD90*$^+$*s, 307 for CD90*$^-$*s, 213 for HPCs).* ***d)*** *Time necessary for completing the second division, for foetal cord blood HSPCs expanded in Diff medium, n = 1268 divisional events (301 for CD90*$^+$*s, 677 for CD90*$^-$*s and 290 for HPCs).* ***e)*** *Same as* ***d),*** *for foetal cord blood HSPCs expanded in GT medium, n = 722 divisional events (243 CD90*$^+$*s, 257 CD90*$^-$*s, 222 HPCs).* ***f)*** *Same as* ***d),*** *for adult bone marrow HSPCs expanded in Diff medium, n = 1474 divisional events (520 for CD90*$^+$*s, 543 for CD90*$^-$*s, 411 for HPCs).* ***g)*** *Time necessary for completing the third division, for foetal cord blood HSPCs expanded in Diff*

*medium; n = 1433 divisional events (271 for CD90$^+$s, 750 for CD90$^-$s and 412 for HPCs).* ***h)*** *Same as* ***g)****, for foetal cord blood HSPCs expanded in GT medium, n = 1179 divisional events (372 CD90$^+$s, 408 CD90$^-$s, 399 HPCs).* ***i)*** *Same as* ***g)****, for adult bone marrow HSPCs expanded in Diff medium; n = 1940 divisional events (520 for CD90$^+$s, 671 for CD90$^-$s, 749 for HPCs).* ***j)*** *Difference between the time necessary to complete the first and the mean of the 2 second divisions, for foetal cord blood HSPCs expanded in Diff medium. This divisional delay encompasses the time necessary for the most immature HSPCs to exit $G_0$. n = 640 (152 for CD90$^+$s, 342 for CD90$^-$s, 146 for HPCs).* ***k)*** *Same as* ***j)****, for foetal cord blood HSPCs expanded in GT medium. n = 362 (122 for CD90$^+$s, 129 for CD90$^-$s, 111 for HPCs).* ***l)*** *Same as* ***j)****, for adult bone marrow HSPCs expanded in Diff medium. n = 745 (265 for CD90$^+$s, 274 for CD90$^-$s, 206 for HPCs).* ***m)*** *Colony size, measured as number of light-refringent cells in the well at T = 96 hours, for foetal cord blood HSPCs expanded in GT medium. 2 independent experiments, n = 375 (132 CD90$^+$s, 127 CD90$^-$s, 116 HPCs). **p < 0.01, ***p < 0.001, **** p < 0.0001, Dunn post-hoc test with Benjamini-Hochberg for multiple corrections.*

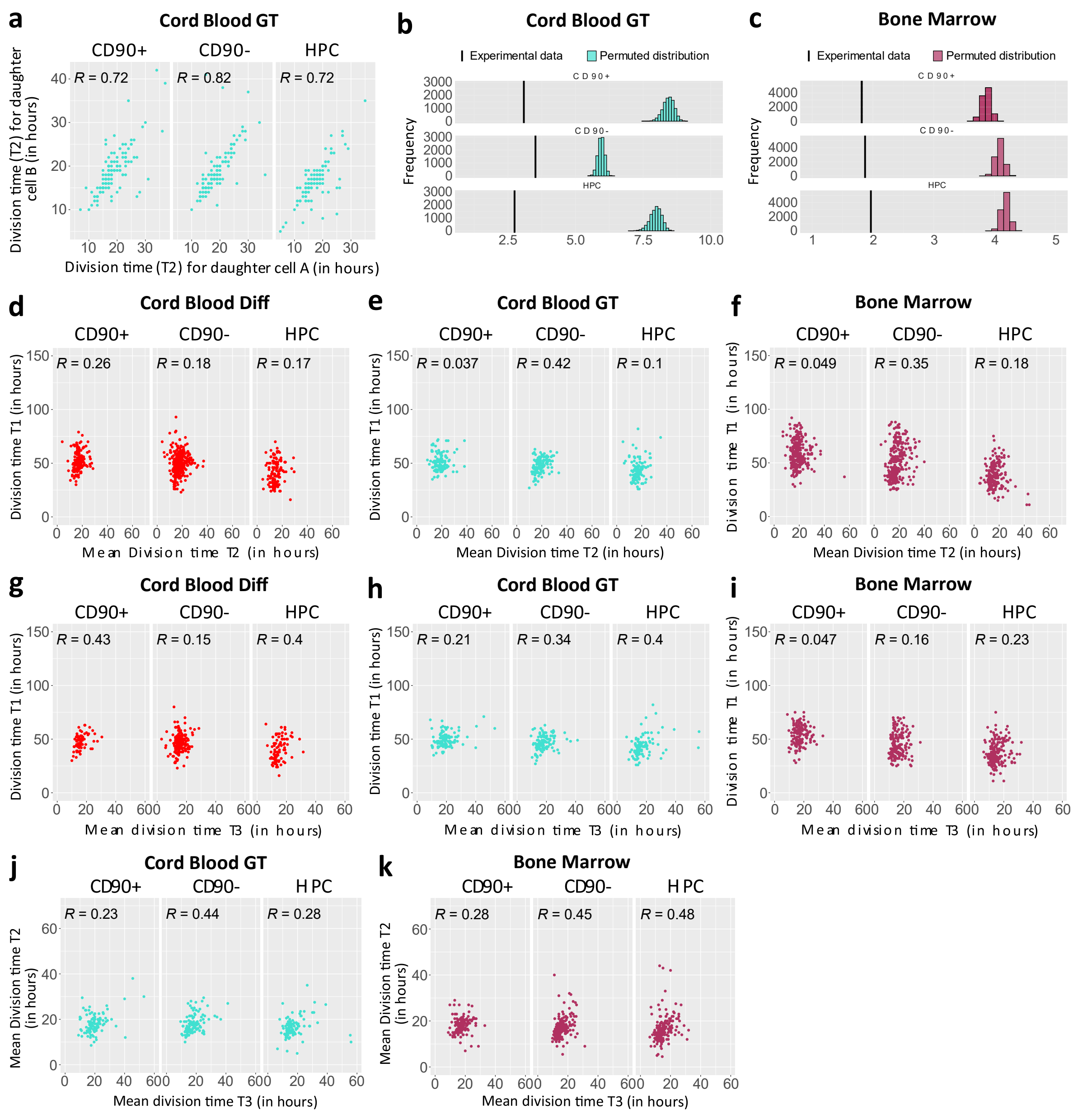


***Extended Data Figure 3: Correlation in cell division lengths, inside and between generations.***

***a)*** *Correlation in division time (expressed in hours) between two sister cells in generation 1, for HSPCs isolated from foetal cord blood and expanded in GT medium. Spearman's rank correlation coefficient R, 2 independent experiments, from 2 different donors, 722 divisional events (243 for CD90*$^+$*s, 257 for CD90*$^-$*s, 222 for HPCs).* ***b)*** *Mean SD of third division time (T3) of cells, derived from the same ancestor cells (black line), and permuted families (cyan histograms), for foetal cord blood HSPCs expanded in GT medium. Only families with the four T3 divisions recorded were used. 2 independent experiments (n = 78 for CD90*$^+$*, n = 92 for CD90*$^-$ *and n = 92 for HPC), p values < 1 x 10*$^{-4}$*.* ***c)*** *Mean SD of third division time (T3) of cells, derived from the same ancestor cells (black line), and permuted families (maroon histograms), for adult bone marrow HSPCs. Only families with the four T3 divisions recorded were used. 8 independent experiments (n = 113 for CD90*$^+$*, n = 159 for CD90*$^-$ *and n = 175 for HPC), p values < 1 x 10*$^{-4}$*.* ***d-f)*** *Correlation in division time (expressed in hours) between the division time T1 and the mean division time T2, for* ***d)*** *foetal cord blood expanded in Diff medium,* ***e)*** *foetal cord blood expanded in GT medium and* ***f)*** *adult bone marrow.* ***g-i)*** *Correlation in division time (expressed in hours) between the division time T1 and the mean division time T3, for* ***g)*** *foetal cord blood expanded in Diff medium,* ***h)*** *foetal cord blood expanded in GT medium and* ***i)*** *adult bone marrow.* ***j-k)*** *Correlation in division time (expressed in hours) between* *the mean division time T2 and the mean division time T3, for* ***j)*** *foetal cord blood expanded in GT medium, and* ***k)*** *adult bone marrow. For panels* ***d-k****, Spearman's rank correlation coefficient R.*

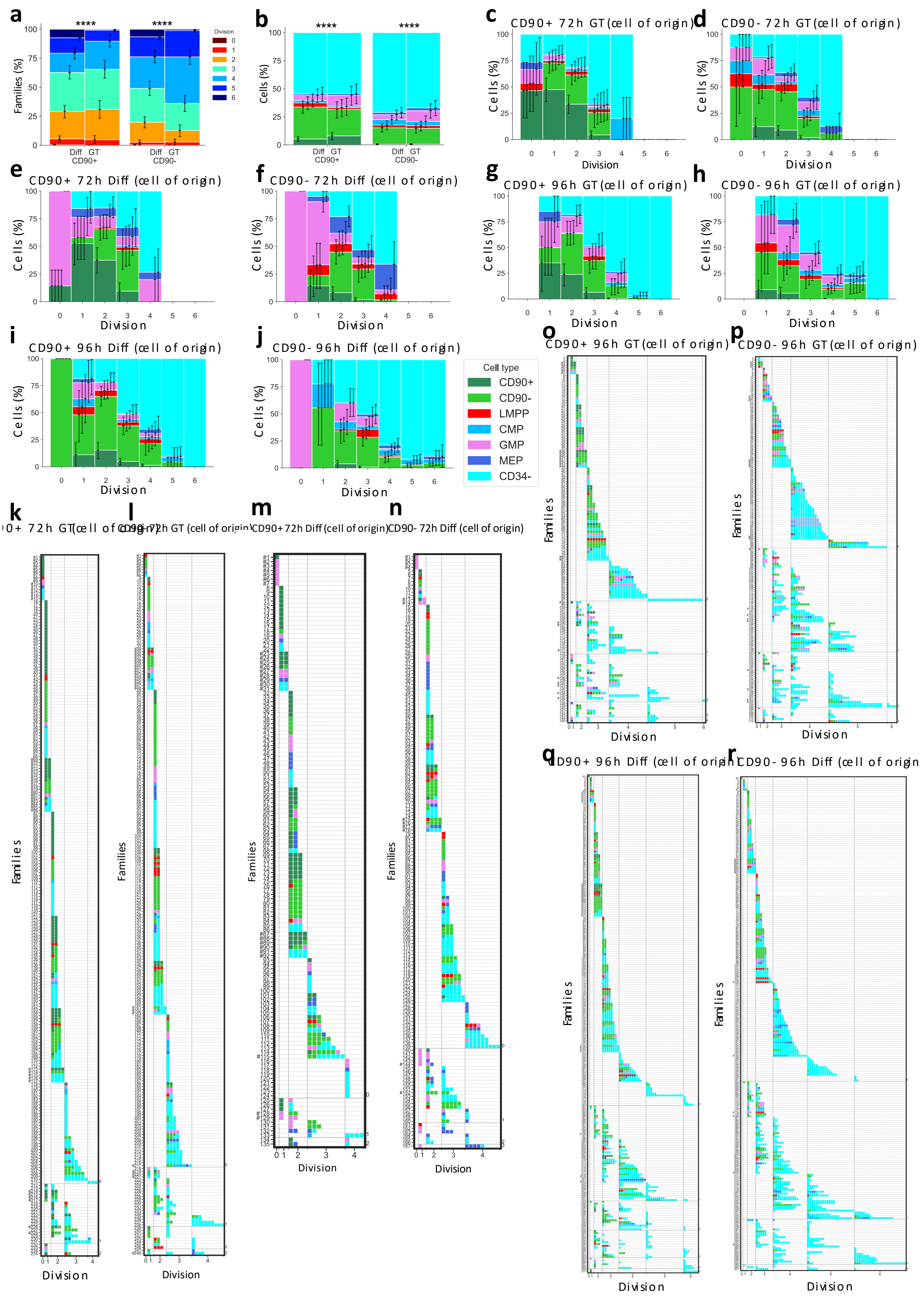


***Extended Data Figure 4: HSPC-derived families display interfamilial heterogeneity, both in cell division and fate commitment. a)*** *Distribution of the maximum division reached by individual families, derived from CD90⁺ and CD90⁻ and expanded in two different media (Diff and GT) for 96 hours of cell culture. Six*

*independent experiments ($n = 145$ for CD90$^+$ GT, $n = 172$ for CD90$^-$ GT, $n = 271$ for CD90$^+$ Diff, $n = 261$ for CD90$^-$ Diff). **b)** Proportions of cell types per families derived from CD90$^+$ and CD90$^-$ and cultured in the GT and Diff media for 96 hours. **c-j)** Cell type percentages per each cell division, for families derived from **c)** CD90$^+$, cultured in GT media for 72h, **d)** CD90$^-$/GT/72h, **e)** CD90$^+$/Diff/72h, **f)** CD90$^-$/Diff/72h, **g)** CD90$^+$/GT/96h, **h)** CD90$^-$/GT/96h, **i)** CD90$^+$/Diff/96h, **j)** CD90$^-$/Diff/96h. **k-r)** Heatmaps representing simultaneously cell division, differentiation, and lineage for single cells. Each row represents an individual family, while columns are ranked according to the number of divisions performed by each cell. Finally, the color code indicates the cell surface phenotype. Families are derived from individual **k)** CD90$^+$, cultured in GT medium for 72h, **l)** CD90$^-$/GT/72h, **m)** CD90$^+$/Diff/72h, **n)** CD90$^-$/Diff/72h, **o)** CD90$^+$/GT/96h, **p)** CD90$^-$/GT/96h, **q)** CD90$^+$/Diff/96h, **r)** CD90$^-$/Diff/96h. For the panels **a** to **j**, the error bars indicate 95% confidence intervals calculated via basic bootstrap. The p values are calculated using the permutation test (Material and Methods section), ****$p < 0.001$*

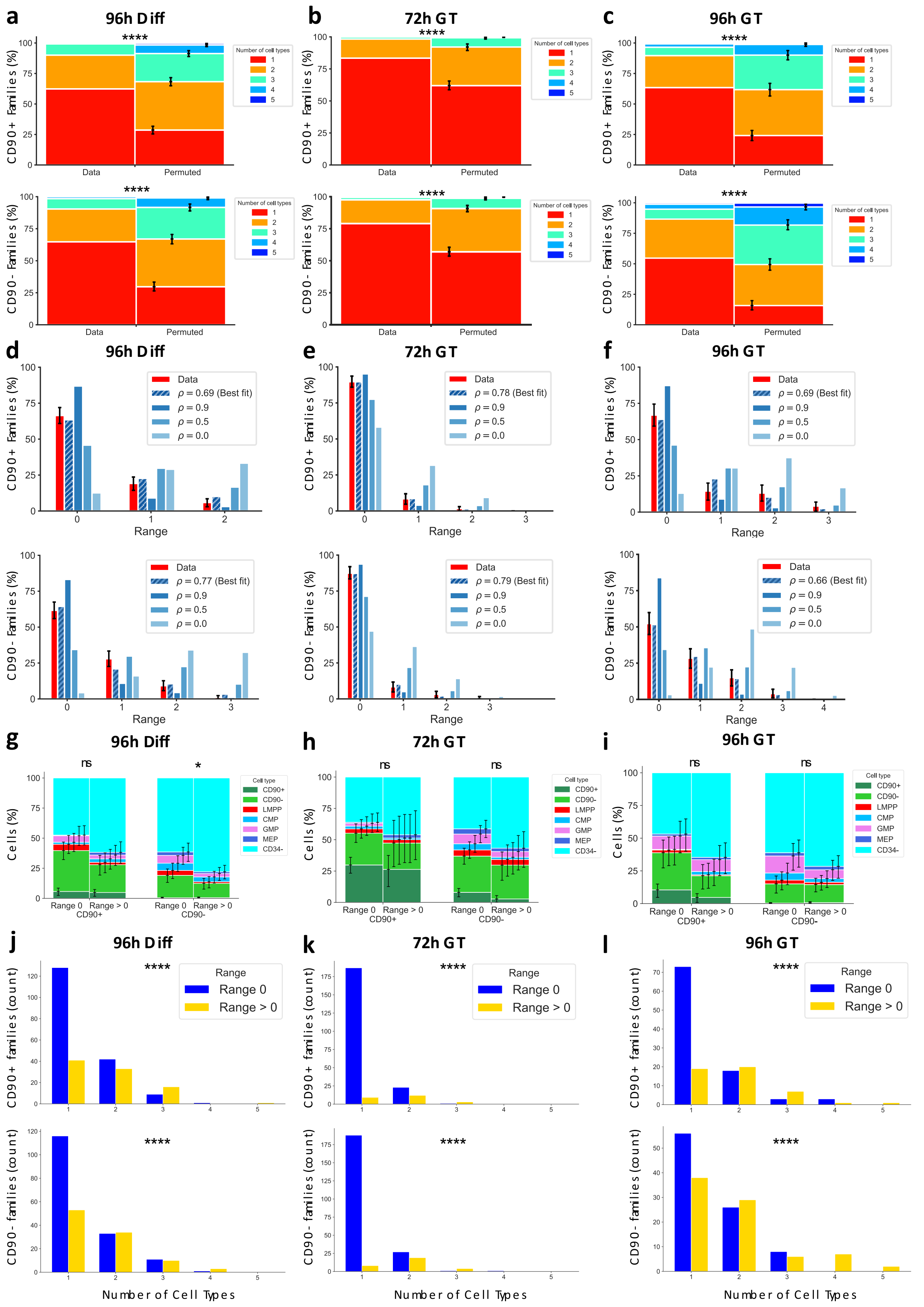


***Extended Data Figure 5: HSPC-derived families display homogeneity in fate and concordance in division at different timepoints and in different experimental conditions. a-c)*** *Distribution of the number of cell types per family, compared with the distribution derived from 250,000*

*permutations of the experimental data:* ***a)*** *CD90$^+$ or CD90$^-$ in Diff medium for 96 hours (Diff/96h),* ***b)*** *CD90$^+$ or CD90$^-$ in GT/72h,* ***c)*** *CD90$^+$ or CD90$^-$ in GT/96h.* ***d-f)*** *Percentage of families per generational range, for cells expanded in* ***d)*** *GT/72h,* ***e)*** *Diff/96h, or* ***f)*** *GT/96h. The red bar corresponds to the experimental data (235/238, 271/261, 145/172 families, respectively, for CD90$^+$/CD90$^-$), while the blue bars describe different simulations, parametrized by a single correlation coefficient $\rho$ indicating the likelihood for a given cell to divide independently from the other cells from the same family. r = 0 means that each member of the family divides independently from the others, r = 1 means that all cells for a given family divide the same number of times, for all families.* ***g-i)*** *Proportions of cell types per families that have a division range = 0 or > 0, derived from* ***g)*** *CD90$^+$ and CD90$^-$ cultured in Diff media for 96 hours,* ***h)*** *CD90$^+$ or CD90$^-$ in GT/72h and* ***i)*** *CD90$^+$ or CD90$^-$ in GT/96h.* ***j-l)*** *Number of progenitor subsets per family, for families with range = 0 or > 0 cultured in* ***j)*** *Diff/96h,* ***k)*** *GT/72h, or* ***l)*** *GT/96h. For panels* ***a-i,*** *the error bars indicate 95% confidence intervals calculated via basic bootstrap. p values are calculated using the permutation test; *p < 0.05, ****p < 0.001*

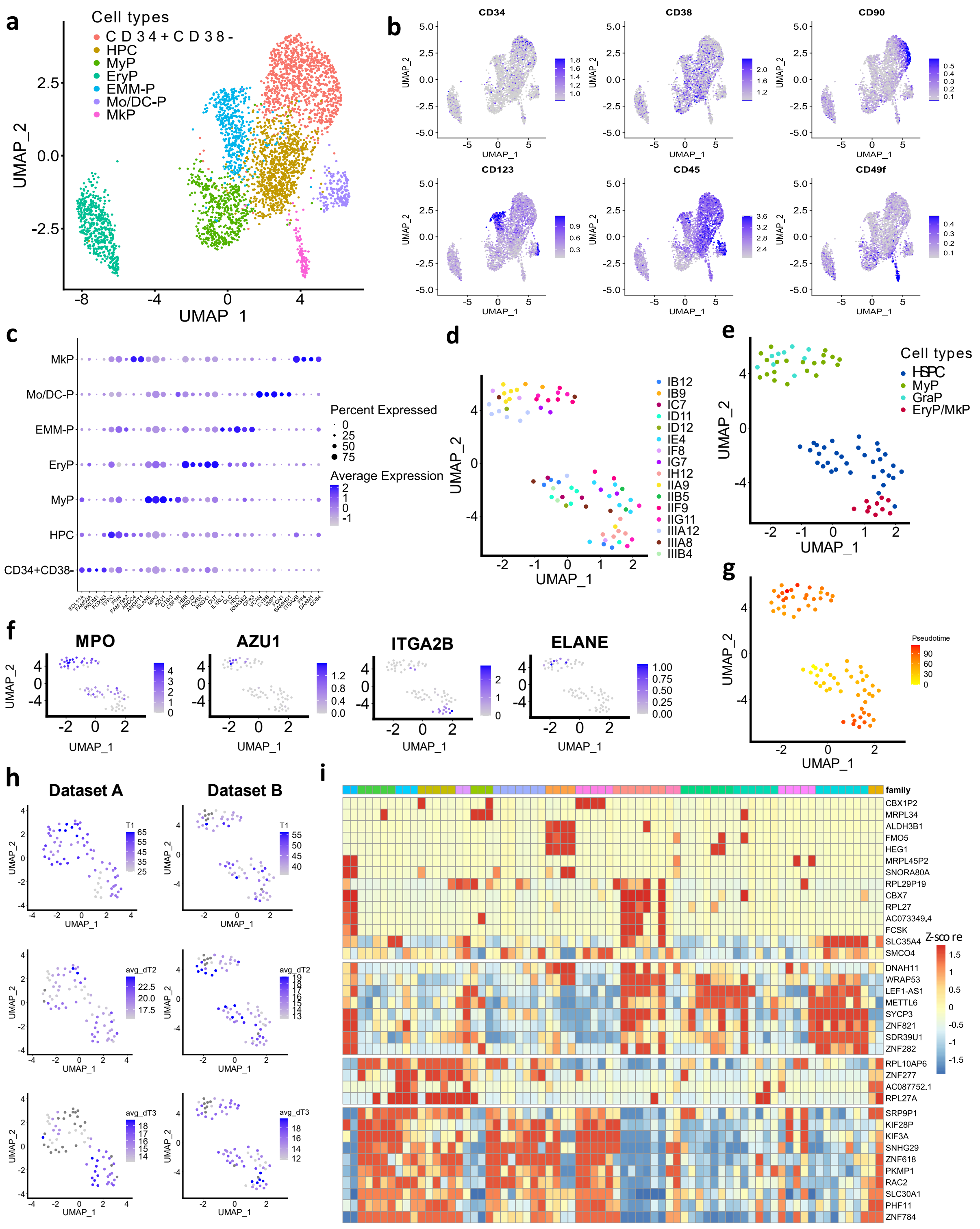


***Extended Data Figure 6: Human HSPCs transcription profile after cell culture*** ***a-b)*** *UMAP representation of the CITE-seq experiment, colour-coded accordingly to the* ***a)*** *annotated cell type and* ***b)*** *ADT expression at the surface.* ***c)*** *Lineage-specific gene expression, for genes used for the CITE-seq annotation.* ***d-e-f-g)*** *UMAP representations of a single cell transcriptomic dataset (72 cells), color-coded accordingly to* ***d)*** *family code,* ***e)*** *annotated cell type,* ***f)*** *lineage-specific genes and* ***g)*** *the pseudotime.* ***h)*** *Display of the time to complete the first three cell divisions values, for the UMAP representations of datasets A and B.* ***i)*** *Heatmap displaying the z-score of the 36 most variant genes (rows) between families from the Dataset B. Cells (columns) are color-coded according to their original family.*

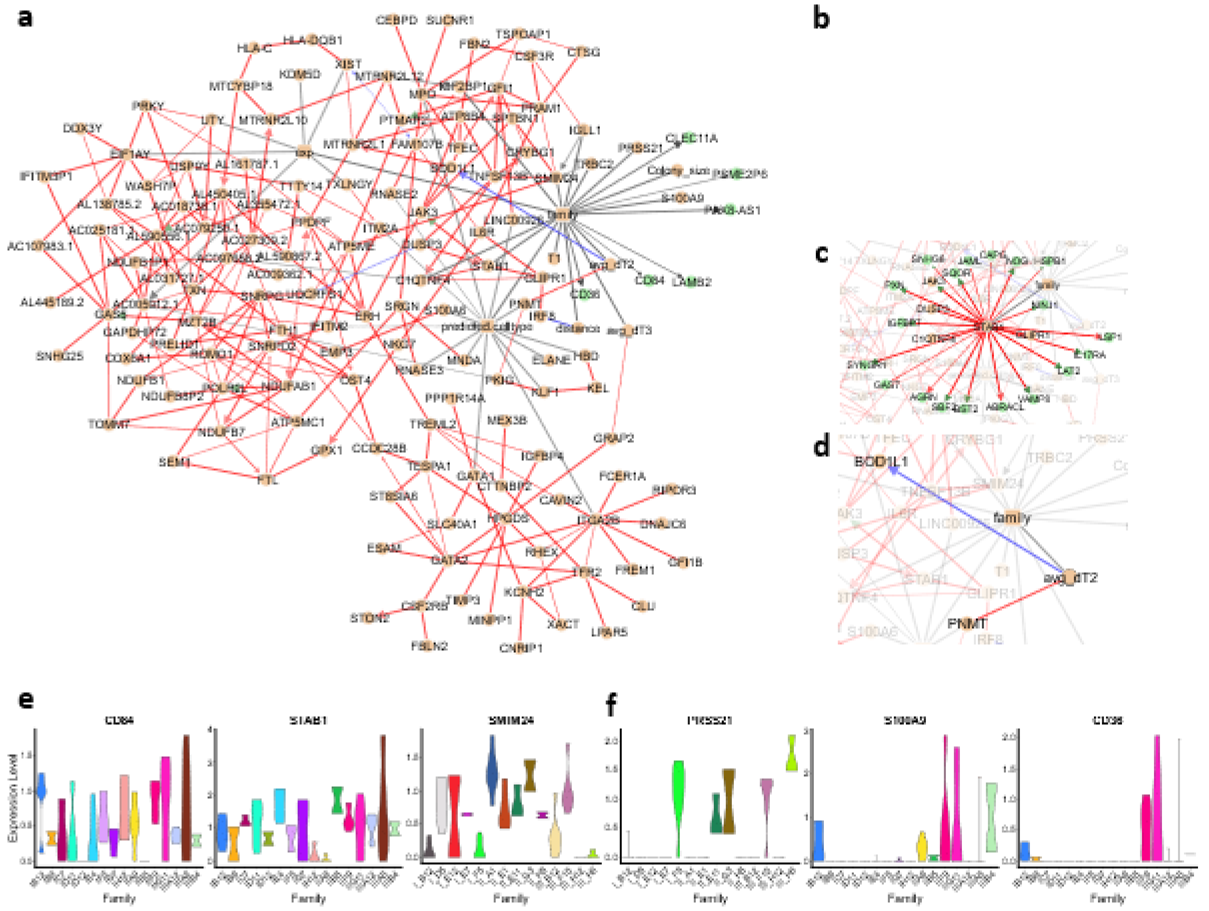


***Extended Data Figure 7: MIIC network analysis reveals genes associated to clonal memory.***
***a)*** *Full MIIC network, including the nodes "family", "predicted.celltype" and "distance" (Cosine distance, defined as the median cosine distance for all the pairs of cells inside a family).* ***b-d)*** *Snapshots of the MIIC network, centred on the variables* ***b)*** *"distance",* ***c)*** *STAB1 and* ***d)*** *"avg_DT2" (mean division time 2).* ***e-f)*** *Gene expression in the dataset B of a selection of "family"-associated genes accordingly to the MIIC analysis, displayed per individual family and organised accordingly to* ***e)*** *genes expressed across families and* ***f)*** *genes whose expression is restricted to a subset of families.*

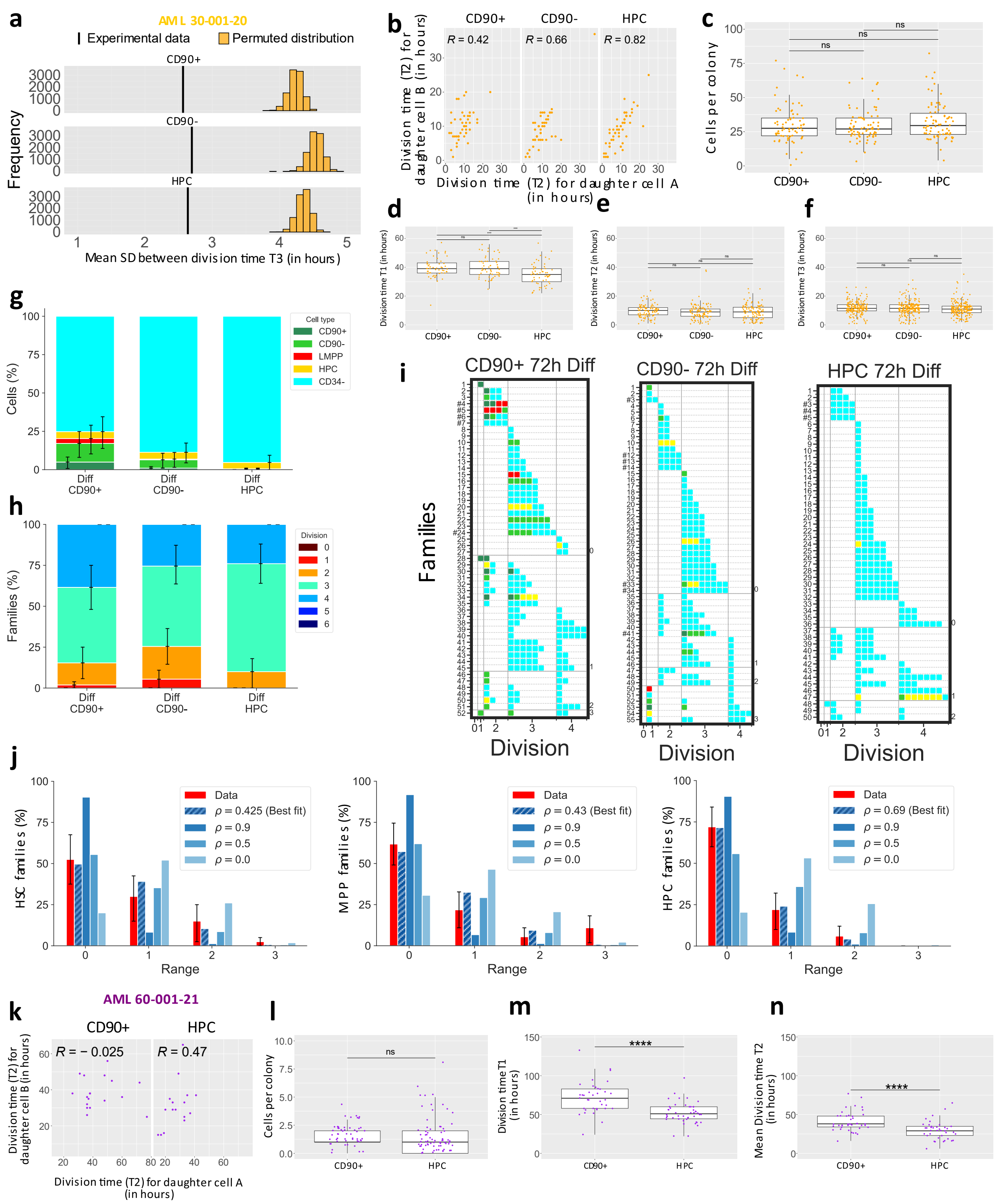


***Extended Data Figure 8: AML blasts display reduced synchronicity in division and other anomalies in divisional properties.*** ***a)*** *Mean SD of third division time (T3) of cells derived from the same ancestor cells (black line), and permuted families (orange histograms), for CD34⁺ from AML patient #30-001-20 peripheral blood. Only families with four T3 divisions recorded were used, n = 44 for CD90⁺, n = 51 for CD90⁻ and n = 51 for HPCs; p < 1 x 10⁻⁴.* ***b)*** *Correlation in division time (expressed in hours) between two sister cells in generation 1, for HSPCs isolated from the same patient, n = 50 for CD90⁺, n = 54 for CD90⁻ and n = 56 for HPCs.* ***c)*** *Colony size, measured as number of light-refringent cells in the well at T = 96 hours, for HSPCs isolated from the same patient; n = 70 for CD90⁺, n = 75 for CD90⁻, and n = 78 for HPCs.* ***d)*** *Time necessary for completing the first division; n = 52 for CD90+s, n = 57 for CD90- and HPC.* ***e)*** *Time necessary for completing the second division, n = 100 for CD90+, n = 108 for CD90- and n = 112 for HPC.* ***f)*** *Time necessary for completing the third division, n*

*= 191 for CD90+, n = 217 for CD90- and n = 215 for HPC.* ***g)*** *Proportions of cell types per families derived from $CD90^+$, $CD90^-$, and HPCs isolated from AML patient #30-001-20 and cultured for 72 hours.* ***h)*** *Distribution of the maximum division reached by individual families, derived from $CD90^+$, $CD90^-$ and HPC isolated from AML patient #30-001-20.* ***i)*** *Heatmaps representing simultaneously cell division, differentiation, and lineage for single cells, for $CD90^+$, $CD90^-$, and HPCs isolated from AML patient #30-001-20.* ***j)*** *Percentage of families per division range, defined as the difference between divisions within each family. The red bar corresponds to the experimental data for $CD90^+$, $CD90^-$ and HPC isolated from AML patient #30-001-20. The dashed blue bar corresponds to the correlation coefficient ρ best fit value. The filled blue bars are simulations, parametrized by different values of ρ.* ***k)*** *Correlation in division time (expressed in hours) between two sister cells in generation 1, for HSPCs isolated from AML patient #60-001-21 peripheral blood, n = 16 for $CD90^+$ and HPC.* ***l)*** *Colony size, measured as number of light-refringent cells in the well at T = 96 hours, n = 59 for $CD90^+$ and n = 80 for HPC.* ***m)*** *Time necessary for completing the first division, n = 37 for $CD90^+$ and n = 46 for HPC.* ***n)*** *Time necessary for completing the second division, n = 41 for $CD90^+$, and n = 38 for HPC. For the panels* ***g***, ***h*** *and* ***j***, *the error bars indicate 95% confidence intervals calculated via basic bootstrap.*

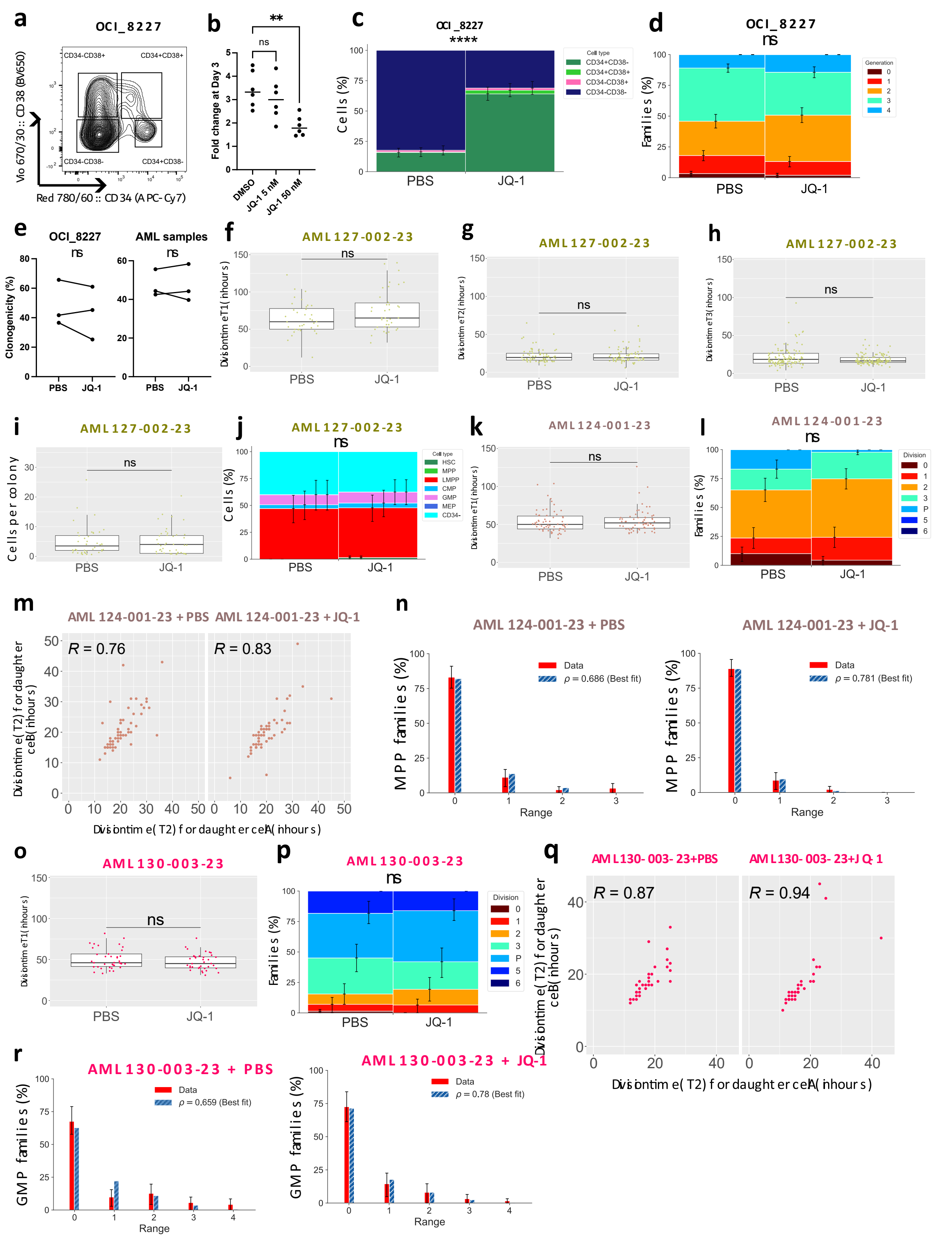


**Extended Data Figure 9: Exposure to epigenetic inhibitor JQ-1 increase the cell division concordance for AML-derived cell families.** ***a)*** *Flow cytometry gating for OCI_8227 cells, based on the expression of CD34 and CD38 and used in the MultiGen experiments.* ***b)*** *Cell numbers fold change for OCI_8227 cells after 72 hours exposure to JQ-1 at 5 and 50 nM. Each condition was assessed in triplicate, n = 2.* ***c)*** *Proportions of cell types per families derived from CD34⁺ OCI_8227 cells, treated with JQ-1 5 nM or PBS for 72 hours.* ***d)*** *Distribution of the maximum division reached by individual families, for OCI_8227 cells exposed to JQ-*

*1 or PBS.* ***e)*** *Clonogenicity for OCI_8227 cells and AML primary cells, expressed as percentage and calculated as the ratio between single progenitors seeded at D0 and number of families analyzed via MultiGen after 72 hours.* ***f)*** *Time necessary for completing the first division, for LMPP isolated from the peripheral blood of patient #127-001-23 and exposed to sublethal doses of JQ-1 or PBS; n = 36 and 38, respectively.* ***g)*** *Time necessary for completing the second division, for the same patient and exposed to sublethal doses of JQ-1 or PBS; n = 72 and 68, respectively.* ***h)*** *Time necessary for completing the third division, for the same patient and exposed to sublethal doses of JQ-1 or PBS; n = 122 and 120, respectively.* ***i)*** *Colony size, measured as number of light-refringent cells in the well at T = 96 hours, for the same patient and exposed to sublethal doses of JQ-1 or PBS; n = 42 each.* ***j)*** *Proportions of cell types per families derived from the same patient and exposed to sublethal doses of JQ-1 or PBS; n = 69 and 68 respectively.* ***k)*** *Time necessary for completing the first division for LMPPs isolated from AML patient #124-001-23, treated for 72 hours with sublethal doses of JQ-1 or PBS; n = 63 each.* ***l)*** *Distribution of the maximum division reached by individual families, for the same patient and exposed to sublethal doses of JQ-1 or PBS.* ***m)*** *Correlation in division time (expressed in hours) between two sister cells from generation 1, for the same patient and exposed to sublethal doses of JQ-1 or PBS; n = 59 each.* ***n)*** *Percentage of families per division range for the same patient and exposed to sublethal doses of JQ-1 or PBS; n = 91 and 89, respectively. The red bars correspond to the data and the dashed blue bar represents the ρ best fit.* ***o)*** *Time necessary for completing the first division, for GMPs isolated from the peripheral blood of patient #130-003-23 and exposed to sublethal doses of JQ-1 or PBS; n = 36 and 38, respectively.* ***p)*** *Distribution of the maximum division reached by individual families, for the same patient and exposed to sublethal doses of JQ-1 or PBS.* ***q)*** *Correlation in division time (expressed in hours) between two sister cells from generation 1, for the same patient and exposed to sublethal doses of JQ-1 or PBS; n = 34 and 31 respectively.* ***r)*** *Percentage of families per division range for the same patient and exposed to sublethal doses of JQ-1 or PBS; n = 62 and 71, respectively). The red bars correspond to the experimental data, the dashed blue bar represents the ρ best fit. For the panel* ***b,*** *1-way ANOVA with Dunnett correction for multiple comparisons, ns for p > 0.05, ** for p < 0.01. For the panels* ***c, d, j, l, n, p*** *and* ***r****, the error bars indicate 95% confidence intervals calculated via basic bootstrap. The p values are calculated using the permutation test; ns, not significant; ****p < 0.001. For panels* ***f, g, h, i, k*** *and* ***o****, ns for p > 0.05, Dunn post-hoc test with Benjamini-Hochberg for multiple corrections. For panels* ***m*** *and* ***q****, Spearman's rank correlation coefficient R. For panel* ***f****, paired T test, ns for p > 0.05.*

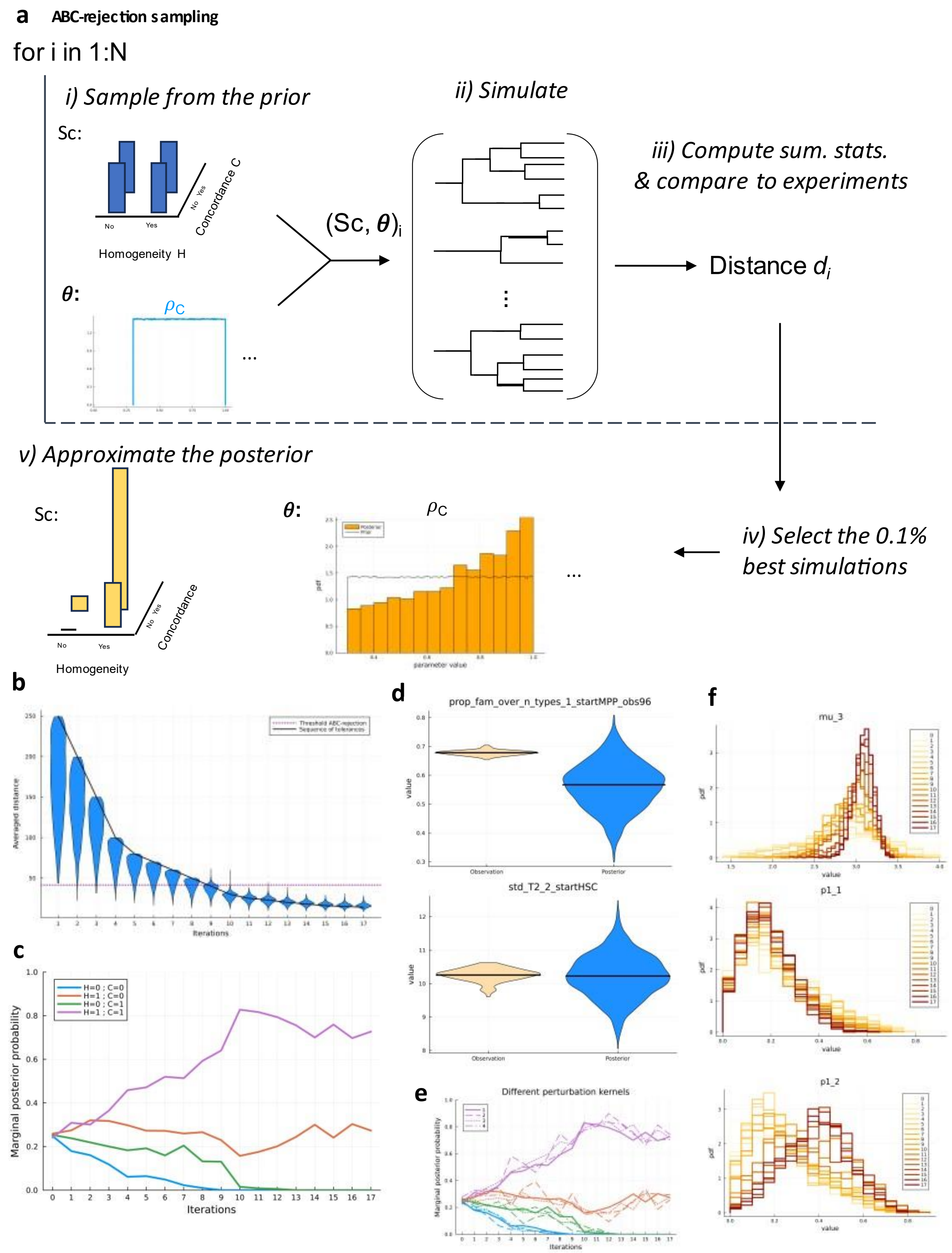


***Extended Data Figure 10: Calibrating the mathematical model.*** ***a)*** *Overview of the ABC rejection sampling algorithm used to estimate the probability of each modelling scenario, repeated N times (N=2,216,106). i) Sampling of the prior for the modelling scenario (denoted by* $Sc_i$ *- H=0 or 1; C=0 or 1), and a parameter vector* $\vartheta_i$*. ii) Simulation of as many dynamics as the number of experimental families (1740). iii) Computation of the summary statistics and calculation of the distance to the observation* $d_i$*. Repeating the comparison N times yields a set of N parameter vectors (and scenarios) associated with distances. iv) Selection of the 0.1% best distances,*

*v) and use to approximate the posterior distributions.* ***b)*** *ABC-SMC method for refining the estimation of the posterior distribution of each modelling scenario. Each iteration i produce the distribution of the (averaged over $B_i$ simulations) distances of all particles (i.e., of all accepted parameter vectors). The horizontal purple dotted line corresponds to the threshold obtained with the previous ABC-rejection sampling algorithm. The black line corresponds to the decreasing sequence of tolerances. For iterations such that $B_i = 1$, then the distances of the accepted particles are necessarily below the threshold (this is the criterion to be accepted). Otherwise, the particles are selected when at least one simulation over the $B_i$ is below the tolerance. After 9 iterations, results improved compared to the ABC-rejection sampling algorithm (panel a). From iteration 15, the distributions of the distances do not evolve anymore (convergence).* ***c)*** *Evolution of the scenarios approximated probabilities over the iterations (with the ABC-SMC method).* ***d)*** *Evaluation of the fitting quality for 2 summary statistics over the 340 generated. The experimental value (left), obtained from random sampling 90% of the experimental observations used for computing the summary statistics, is compared to the model obtained when sampling the posterior distribution (right).* ***e)*** *Robustness of the ABC-SMC method. Evolution of the approximated probabilities of our four scenarios (purple: H=1, C=1; orange: H=1, C=0; green: H=0, C=1; blue: H=0, C=0) over iterations when using different perturbation kernels. The numbers in the legend indicate the configuration, with configuration 1 being the same as in panel c).* ***f)*** *Intermediate marginal posterior distributions of three model parameters (for the most likely modelling scenario), from iteration 0 (the prior) to iteration 17 (our approximated posterior). Lighter colours correspond to the initial iterations, darker colours correspond to the last ones. Mu_3 (median value of the log-normal distribution modelling HPC division) corresponds to a rapid convergence to the posterior distribution). p1_1 (probability that a sister cell from an CD90+ will also be an CD90+) do not move from the prior, indicating that the observation model does not convey enough information to properly identify the value of this parameter. p1_2 (probability that a sister cell from an CD90+ will be an CD90-) do move away from the posterior, but lower to those in parameter mu_3.*
