## Supplementary information for "Clonal memory of cell division in humans diverges between healthy haematopoiesis and acute myeloid leukaemia"

### Supplemental Information -
### Mathematical modelling and statistical inference

### Material and Methods

In this Supplemental Information, we present the elements of the mathematical methodology. In section A, we precise which experimental observations are included in our methodology, how we pretreat the data, and we introduce the notations.
In section B, we detail our mathematical model and the different scenarios we consider. In section C, we describe the observation and statistical model, i.e., how we confront the output of our dynamic model to the experimental observations. In section D, we present the methodology to select among the four different scenarios considered, and in section E, we extend our study to more scenarios.
Most part of the work presented here is associated with a computational code, implemented in Julia, and available at:
`https://gitlab-research.centralesupelec.fr/2012hermangeg/HSPC_proliferation`

### A Experimental observations - pretreatment and notations

The content related to the experimental observations and the assumptions made in this section concern only this supplemental information, that is, the mathematical methodology.

#### A.1 Considered experiments

Experimental observations used for this work come from two experiments: Live Cell Imaging (Incucyte©) and MultiGen. They are derived from cord blood samples, purified to retain stem and progenitor cells (i.e., $CD34^+$ cells). The samples come from different individuals and are pooled together. We, therefore, consider that we can neglect the heterogeneity between individuals. The mathematical methodology is also only applied for the experiments made with the culture condition "Diff".

#### A.2 Cell types

We will consider four possible cellular types: HSC, MPP, HPC, and $CD34^-$ (respectively associated with indices 1, 2, 3, and 4, in ascending order of maturity), without taking into account the differences in surface marker expression within each of these types.

#### A.3 Family, well, colony

We can refer interchangeably to the term family, well, or colony. The experiments are based on the observation of cells, during their first divisions (Live Cell Imaging, see Fig. A.1) or at a given observation time (MultiGen, see Fig. A.5), starting from a cell that has been cultured in a well. We call family the colony of cells created from this initial cell. When, in the MultiGen experiment, multiple starting cells are cultured in the same well, we assume that this is equivalent to having four wells, each with its starting cell (thus, we assume that there are no interactions between families within a well). This is represented in Figure A.5. Hence the fact that we can talk interchangeably about wells or families.

#### A.4 Live Cell Imaging Experiment

In this experiment, we can observe the times of the first, second, and third divisions of an initial cell of a given type $p \in \{\text{HSC, MPP, HPC}\}$, cultured in a well in the presence of a cytokine cocktail (Diff) for $T_{max} = 96$ hours. From this experiment, it is possible to know the time interval ($\Delta t = 1$h) during which the divisions of interest took place. However, it is not possible to know the cellular type of the cells occurring during the dynamics (apart from the starting cell).

The observations are censored by interval, where we observe the upper bound of the one-hour time interval during which the division took place. In practice, we will consider left interval censoring. This is so that divisions not observed are assigned the maximum observation time, namely, 96 hours (instead of $+\infty$ otherwise).
For $T_1 \in \mathbb{R}^+$ corresponding to the exact (unobserved) first division time, we note $D_1 \in \mathbb{N}$ its integer part (then, $D_1 \leq T_1$). Idem for the two second division times $D_{2,1}$ and $D_{2,2}$, where $D_{2,1} \leq D_{2,2}$, and

also for the third division times: $D_{3,1} \leq D_{3,2} \leq D_{3,3} \leq D_{3,4}$.

We exclude from the dataset any family that does not have at least two cells at 96 hours.
Our dataset thus consists of the observation of 152, 361, and 146 families starting respectively from an HSC, MPP, or HPC. We denote $\mathcal{I} = \bigcup_{p \in \{1,2,3\}} \mathcal{I}_p$ the set of families observed by the Live Cell Imaging experiment, distinguishing according to the initial condition, that is, if the starting cell type is HSC (index 1), MPP (index 2), or HPC (index 3).

To sum up, in the Live Cell Imaging experiment, we observe, for a given initial cell type (HSC, MPP, or HPC), the time of the first division, the times of the second division, as well as the colony size (i.e., the number of cells) at 96 hours, and in some cases the third division times.
It is possible that, for a given well, we do not observe the data of both $D_1$, $D_{2,1}$, $D_{2,2}$, and colony size. The dataset will then be divided into three:

- The list of first division times $D_1$, for a given starting cell type. This dataset will be used to determine the empirical distribution of first division times (see § B.3.1),
- The set of triplets $(D_1, D_{2,1}, D_{2,2})$ - according to the initial cell type - which will be used for calculating summary statistics (see § C.2). Note that the summary statistics associated with the Live Cell Imaging experiment will be calculated for each of the 3 datasets separately:
  - 152 triplets observed starting from an HSC
  - 361 triplets starting from an MPP
  - 146 starting from an HPC
- The list of colony sizes at 96 hours (according to the starting type), which will be used for calculating some summary statistics.

Those data are available in the folder `data/` at:
`https://gitlab-research.centralesupelec.fr/2012hermangeg/HSPC_proliferation`

When the times of the third divisions are reported, the information will be also used for computing some summary statistics related to the third division time (see § C.2 for more details about the summary statistics).

Figure A.2 shows the distribution of first division times, which will also be used in the model.
Figure A.3 shows the distribution of second division times (aggregate of $D_{2,1}$ and $D_{2,2}$), as well as their joint distribution, varying according to the starting cell type.
Figure A.4 illustrates the distribution of colony sizes at 96 hours, based on the initial cell type.

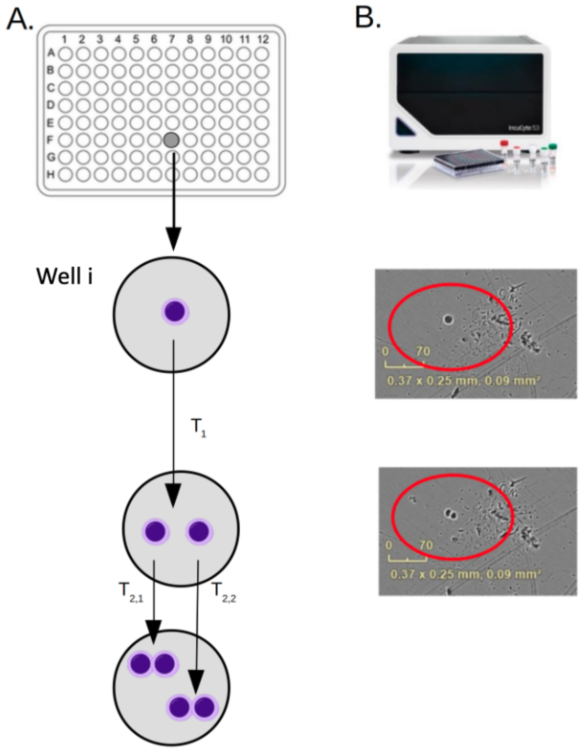

Figure A.1: Schema of the Live Cell Imaging (Incucyte©) experiment. Cells of a known type are cultured in the presence of a cytokine cocktail, one cell per well (A). The cell will divide after a time $T_1$, and then the divisions continue (associated with a differentiation process). An image is taken every hour (B), which then allows for determining the time interval during which the divisions occurred. $T_{2,1}$ and $T_{2,2}$ are measured from the start of the experiment; to get an idea of the time required for each of the two sister cells to divide, $T_1$ should be subtracted from them.

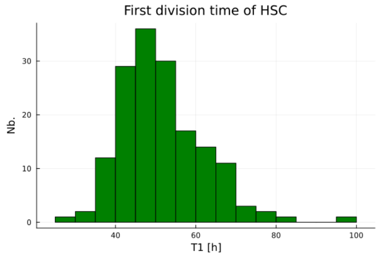

(a) HSC

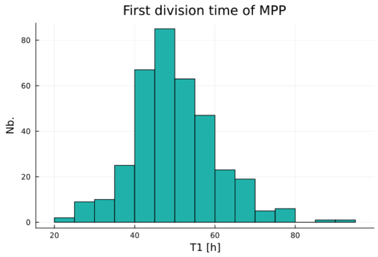

(b) MPP

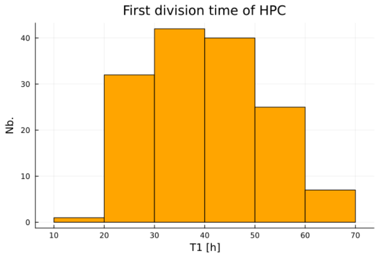

(c) HPC

Figure A.2: Distribution of the first division times ($D_1$) observed starting from an HSC (left), MPP (center), or HPC (right). Divisions not observed before 96 hours are indicated at 96 hours (due to left interval censoring, with the last interval $[96, +\infty[$).

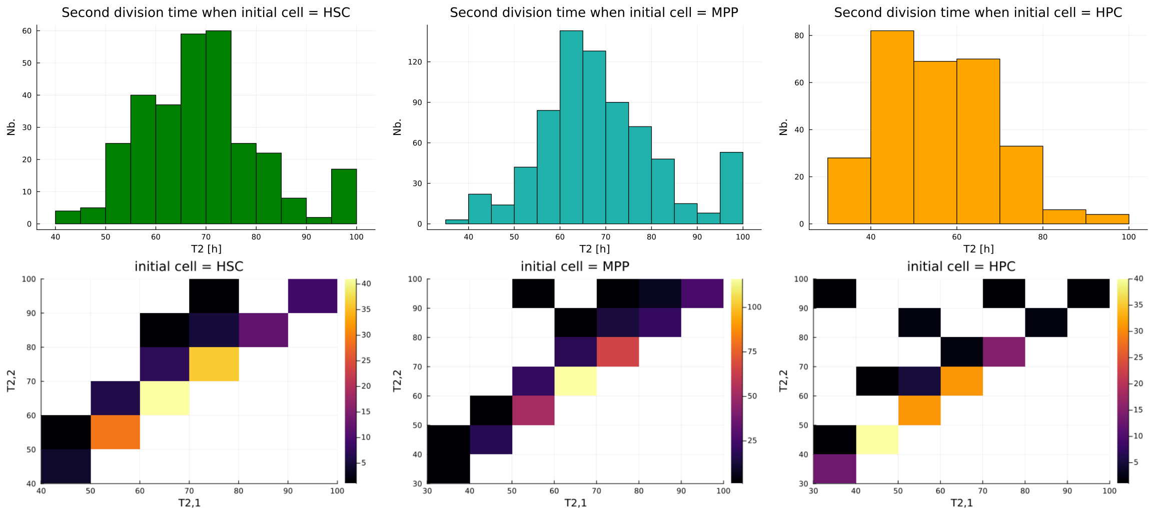

Figure A.3: Top: Distribution of second division times according to the cellular type of their mother cell (aggregate of $D_{2,1}$ and $D_{2,2}$). Bottom: Joint distributions of $D_{2,1}$ and $D_{2,2}$ with $D_{2,2} \geq D_{2,1}$. Note: these are the times of divisions counted from the start of the experiment, including information on the time of the first division.

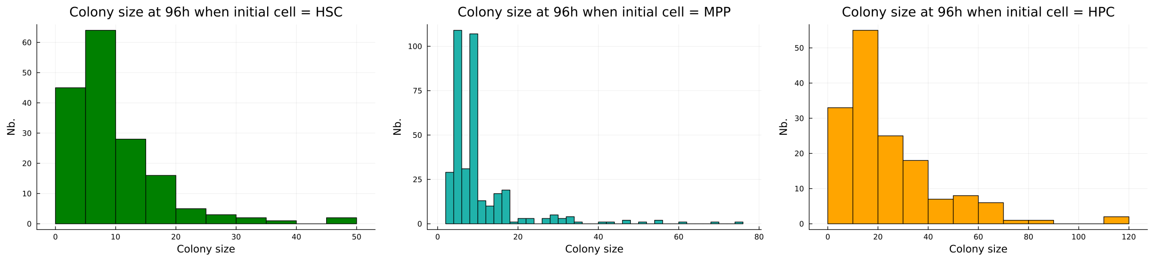

(a) HSC (b) MPP (c) HPC

Figure A.4: Distribution of colony sizes at 96 hours according to the initial cell type.

#### A.5 MultiGen Experiment

The MultiGen experiment, initially described by Horton et al. [S6] and used by Tak et al. [S12] for the study of HSPCs (Hematopoietic Stem and Progenitor Cells, including HSC, MPP, and HPC) in mice, is schematized in Figure A.5. The experimental protocol for this assay in described in detail in [S3]. In this experiment, cells are cultured in wells in the presence of a cytokine cocktail (Diff cocktail), with four cells per well. We know the initial type $p \in \{\text{HSC, MPP, HPC}\}$ of each cell, that is, the initial condition (Fig. A.5-A). For a given well, the four cells are each marked with fluorescent dyes that allow distinguishing, after $T_{max} = 72$ or 96 hours, the cells derived from each of them (Fig.A.5-D), but also to know the number of divisions undergone, counted from the initial cell (Fig.A.5-F). We also use the term generation, with the first generation being that of the initial cell. A cell in the third generation will thus have undergone two divisions.
The principle of tracking divisions is based on the dilution of the fluorescence marker during mitosis, with an equitable distribution to the two daughter cells. Two fluorescence markers are used here, CTV and CFSE. Among the four starting cells, one is marked only with CFSE, another only with CTV, the other two with both markers, but in different proportions which will allow distinguishing the families of each of them (the proportions being assumed to remain the same during the divisions).
The dilution of fluorescence markers during divisions means that, after a certain number of divisions, it becomes difficult to discern the exact number of divisions undergone. In our dataset, we do not observe cells beyond generation 7. We consider that beyond 7 generations, we would no longer be able to distinguish the real generation (due to the dilution of fluorescence markers), **meaning any cell with an actual generation higher than 7 will be assigned the value 7**.

The set of cells deriving from a common ancestor is generally called a colony. However, here, we prefer the definition of Tak et al. [S12] and will call this set of cells a "family." The choice of term is not neutral, it emphasizes the idea of a transmission of certain elements from one generation to another. In this case, Tak et al. showed in mice that the fate decisions of cells belonging to the same family could be inherited from their ancestor.
After $T_{max}$, the cells produced during the dynamics of proliferation and differentiation are recovered. They undergo a washing step and are then marked with surface markers, which then allows deducing their cellular type (Fig. A.5-E). Various surface markers are used, namely CD10, CD123, CD34, CD38, CD45, and CD90, which would theoretically make it possible to distinguish among the less differentiated cell types ($CD34^+CD38^-$) the HSC from MPP and LMPP (lymphoid-myeloid common progenitors), and among the slightly more differentiated cells ($CD34^+CD38^+$) the MEP (megakaryocyte-erythroid common progenitors), CMP (myeloid common progenitors) and GMP (granulocyte-monocyte progenitors). To simplify - especially because certain cellular types such as LMPP, GMP, or CMP were underrepresented - we will only consider here the markers CD34, CD38, and CD90, and we will thus study four cell types: HSC, MPP, HPC, and $CD34^-$. Note that the $CD34^-$ cell type, more mature, is heterogeneous and might include cells committed to different differentiation pathways.
To define the cell type based on the expression of surface markers, expression thresholds are set based on the expression of cells in control wells.

During the retrieval of the cells and the subsequent experimental steps, a part of the cells in the well is lost. Thus, we cannot observe the entire contents of the well, but only a fraction (Fig. A.5-B and C). **We call the recovery rate the variable $\eta$ equal to the ratio of the number of cells recovered to the number of those present in the well**. We assume this variable is independent of the number of cells in the well. We mentioned above that four distinct cells were deposited per well. It allows multiplying by four the number of families observed for a single plate. We assume that the colonies are independent of each other, that is, we assume there is no interaction between the cells. Under this assumption as well as the one that the recovery rate is independent of the number of cells actually present, then we can consider it equivalent to consider an experiment with four times more wells, in which a single cell would be cultured per well. We will consider this latter case and propose in the following a model at the family level.

We denote $\mathcal{J} = \bigcup_{p \in \{1,2,3\}} \bigcup_{T_{max} \in \{72,96\}} \mathcal{J}_{p,T_{max}}$ the set of families observed by the MultiGen experiment, distinguishing according to the observation time $T_{max}$ and to the initial condition, that is, if the starting cell type is HSC (index 1), MPP (index 2), or HPC (index 3) .

The set of observed families is as follows:

- 135 families for $T_{max} = 72$h starting from an HSC
- 167 families for $T_{max} = 72$h starting from an MPP
- 271 families for $T_{max} = 96$h starting from an HSC
- 261 families for $T_{max} = 96$h starting from an MPP
- 247 families for $T_{max} = 96$h starting from an HPC

Summary statistics associated with the MultiGen experiment will be calculated for each of these 5 datasets $\mathcal{J}_{p,T_{max}}$, separately.

Figure A.6 gives a representation of 100 wells observed at 96h, when starting with an HSC, through the MultiGen experiment.

To note also, although we will assume that there is no de-differentiation when constructing our model, we sometimes observed cells labeled as less mature than the initial cell types. These observations do not necessarily indicate that there has been de-differentiation, but would rather be related to uncertainty in determining a cell type from a gating strategy.

Figure A.6 shows heatmap for observations at 96 hours for different wells/families, starting from an HSC.

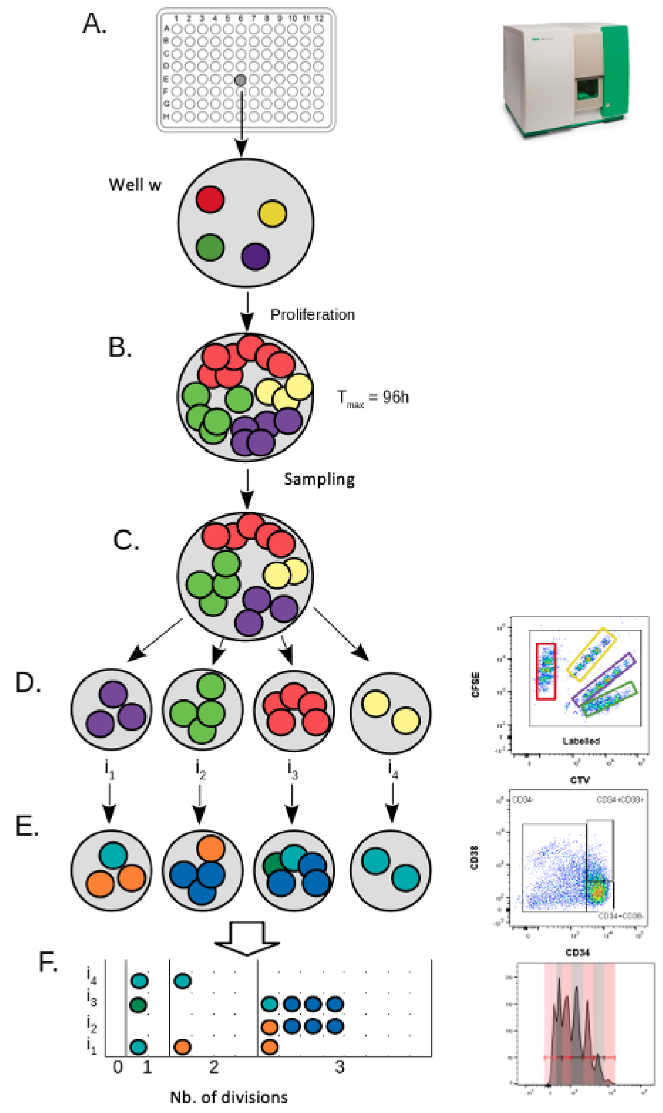

Figure A.5: Schema of the MultiGen experiment. In 96-well culture plates, four cells of the same type are placed in each well (except for control wells used to define expression thresholds for the markers). These four cells correspond to four distinct families (schematized by four different colors: red, yellow, green, and violet). The cells are cultured for 96 hours and then analyzed by flow cytometry. At 96 hours, the initial cells have led to the creation of colonies (families) (B). The contents of the well are then collected. The cells undergo a wash and are then stained; a certain number of cells will be lost during this step (sampling) (C). For the sampled cells, the family to which they belong is identified based on the concentration of two fluorescence markers, CTV and CFSE (D). The identified families consist of cells of different types among HSC (green), MPP (cyan), HPC (orange), or $CD34^{-}$ (blue), which can be identified by flow cytometry using surface markers (E). By also studying the dilution of the fluorescence markers during mitoses, it is also possible to determine the number of divisions each cell has undergone (F).

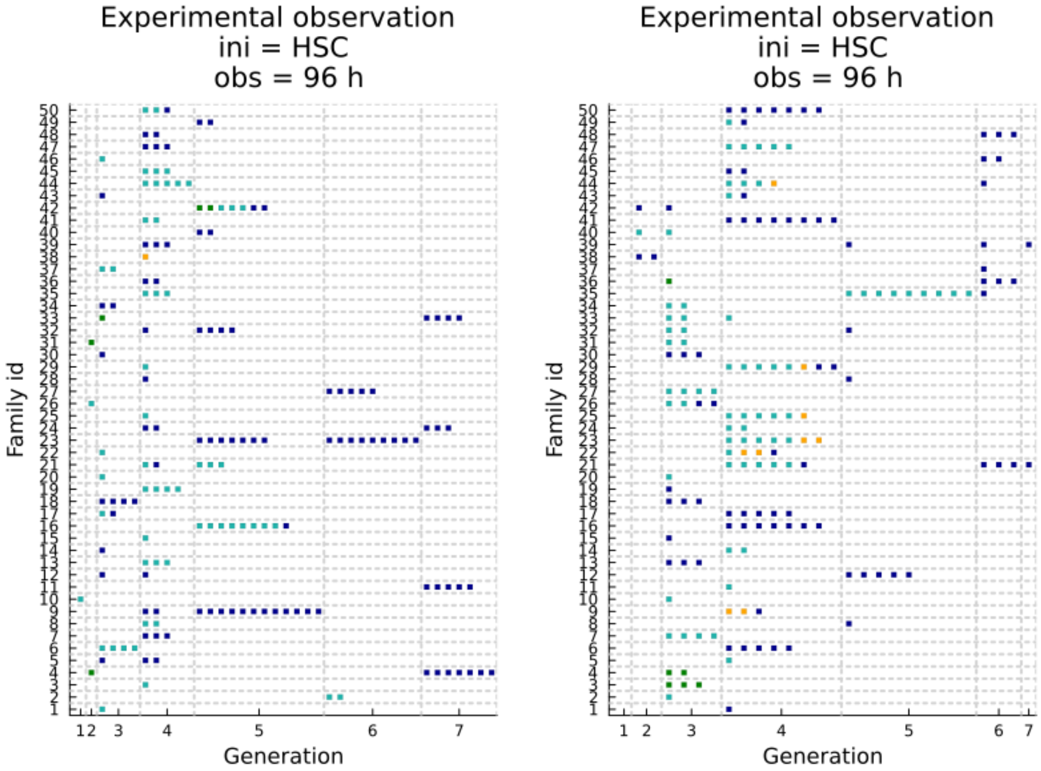

Figure A.6: Heatmap presenting observations at 96h for different wells/families, starting from an HSC. Each line indicates an observed family, with each colored square within that line representing an observed (sampled) cell. The color indicates its type (green: HSC; cyan: MPP; orange: HPC; dark blue: $CD34^{-}$), and the column indicates the generation (with the initial cell being in generation 1). The family IDs here are only for identifying each line in the graphic representation and do not uniquely identify the observed family.

### B Dynamical model(s)

#### B.1 Formalism

In the following, we present a generic model to describe the dynamics of proliferation and differentiation of stem and progenitor cells (HSPC). Variations of this model (i.e., different scenarios) will be considered to account for different modelling hypotheses. This (generic) model has been constructed so that its parameters can be estimated from the experimental observations described in section A: it is notably a stochastic model of short-term dynamics (96 hours), continuous in time, at the scale of a family, in which each cell can only be characterized by a discrete type $p \in \mathcal{P} = \{\text{HSC, MPP, HPC, CD34}^-\}$.

The system we model will be entirely characterized by the state variables $N$, $\boldsymbol{p} = (p_1, \cdots, p_N)$, $\boldsymbol{m} = (m_1, \cdots, m_N)$, and $\boldsymbol{T_A} = (T_{A,1}, \cdots, T_{A,N})$ (see Fig. B.1).
$N \geq 1$ corresponds to the number of cells that existed between the beginning ($t = 0$) and the end ($t = T_{max}$) of the experiment. The $N$ cells are then indexed by $j \in \{1, \cdots, N\}$ corresponding to their order of appearance (with index $j = 1$ corresponding to the initial cell). For a given cell $j$, it is characterized by its type $p_j \in \mathcal{P} = \{\text{HSC, MPP, HPC, CD34}^-\}$, the index of its mother cell $m_j \in \{0, \cdots, j-1\}$ (with $m_1 = 0$), and the time at which it appeared $T_{A,j} \in [0, T_{max}]$ (with $T_{A,1} = 0$). As the indices correspond to the order of appearance of the cells, we have:

$$\forall\ 1 \leq j \leq j' \leq N,\ T_{A,j} \leq T_{A,j'} \tag{B.1}$$

A cell can only divide into two, and the two daughter cells appear at the same moment, so we have:

$$\forall\ 1 < j' < N,\ \text{such that } j' \text{ is even}, m_{j'} = m_{j'+1} \text{ and } T_{A,j'} = T_{A,j'+1} \tag{B.2}$$

Note that **we assume that there is no cell death** (in consistence with the experimental observations); $N$ is necessarily odd.

For any value of $N$, $\boldsymbol{p}$, $\boldsymbol{m}$, and $\boldsymbol{T_A}$ that respect the constraints of relations (B.1) and (B.2), there corresponds a state of the system (Fig B.1.A) from which we can deduce the process of proliferation and differentiation over time $t \in [0, T_{max}]$ (Fig. B.1.C). Indeed, at each time $t$, we can deduce the set $\mathcal{C}_t \subset \{1, \cdots, N\}$ of cells present at that moment (Fig. B.1.B). For $j \in \{1, \cdots, N\}$, $j \in \mathcal{C}_t$ if and only if the following two conditions are met:

1. $T_{A,j} \leq t$
2. $\forall\ j' \in \{j+1, \cdots, N\} : (m_{j'} = j) \Rightarrow (T_{A,j'} > t)$

With the above definitions, it is thus equivalent to consider the system or the associated dynamic process. We prefer to use the term process. In the following, we will describe a stochastic model; $N$ is then a random variable and $\boldsymbol{p}$, $\boldsymbol{m}$, and $\boldsymbol{T_A}$ random vectors (note that they are of size $N$). Modeling this process will involve specifying the underlying probability laws.

If, for example, we had chosen to model division times with exponential distributions, to describe, conditional on the cell type of the mother cell, the possible types of daughter cells and their associated probabilities with a transition matrix, and finally to assume cells are independent of each other, we would then be in the case of a continuous-time multitype branching process, which is a Markovian process (see for example [S1]).

In the following sections, we will describe in more detail our modeling assumptions that lead to a specification of the model more complex than that of branching processes, which will then require studying the model by Monte-Carlo type simulations, particularly for discriminating between variants of the model (scenarios) or for parameter estimation.

The dynamic model is implemented in the Julia programming language. The implementation is available in the folder `"dynamic_model/"` at the following link:
`https://gitlab-research.centralesupelec.fr/2012hermangeg/HSPC_proliferation`

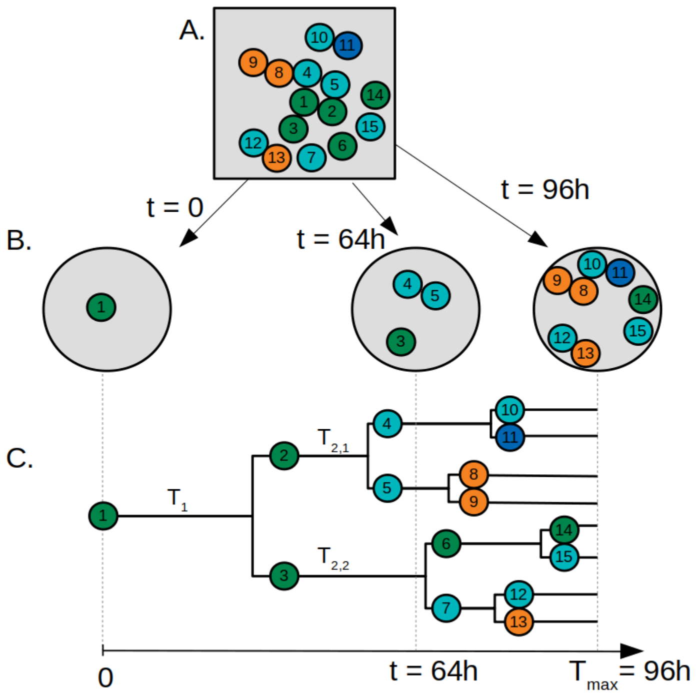

Figure B.1: Illustration of the system studied, with the following values for the state variables: $N = 15$, $\boldsymbol{p} = (1,1,1,2,2,1,2,3,3,2,4,2,3,1,2)^t$ associating the types HSC (green color), MPP (cyan), HPC (orange), and CD34$^-$ (dark blue) with values 1, 2, 3, and 4 respectively, $\boldsymbol{m} = (0,1,1,2,2,3,3,5,5,4,4,7,7,6,6)^t$ and $\boldsymbol{T_A} = (0,34,34,60,60,65,65,68,68,72,72,76,76,90,90)^t$. From the state of the system (A), it is possible to deduce the state of the system at any time $t$ (B) and reproduce the process of proliferation and differentiation over time (C). $T_1$ will correspond to the time of the first division, and $T_{2,1}$ and $T_{2,2}$ to the times of the second divisions such that $T_{2,1} \leq T_{2,2}$. Here, $T_1 = T_{A,2} = T_{A,3}$, $T_{2,1} = T_{A,4} = T_{A,5}$ and $T_{2,2} = T_{A,6} = T_{A,7}$.

#### B.2 Differentiation

##### B.2.1 Transition between cell types

We define differentiation as the transition from one cell type $p \in \mathcal{P}$ to another less immature type, with the set $\mathcal{P}$ thus equipped with an order relation. From the most to the least immature among the types considered, we have:

1. HSC
2. MPP
3. HPC
4. CD34$^-$

Henceforth, we will equate the set $\mathcal{P}$ with the set $\{1,2,3,4\}$.
**We neglect the heterogeneity between cells of the same cell type**, which is a simplification, particularly in the case of CD34$^-$ cells, these latter being known to be heterogeneous.
**We assume that differentiation can only occur following a division**, that is, a cell will not change its cell type during its lifespan. This hypothesis is generally justified [38] even though observations have reported cell differentiation in the absence of divisions [S5, S9]. Finally, **we assume that there is no phenomenon of de-differentiation**, meaning a cell cannot, through division, produce a cell of a more immature type.

Let $k$ be one of the two daughter cells (arbitrary) resulting from the division of cell $m_k$. Its type $p_k$ is a discrete random variable, taking values in $\mathcal{P}$ with the probability $\mathbb{P}[p_k = j | p_{m_k} = i] = M_{i,j}$ where

$\boldsymbol{M} = (M_{i,j})_{1 \le i,j \le 4}$ is the following transition matrix:

$$\boldsymbol{M} = \begin{pmatrix} p_{1\to 1} & p_{1\to 2} & p_{1\to 3} & 1 - p_{1\to 1} - p_{1\to 2} - p_{1\to 3} \\ 0 & p_{2\to 2} & p_{2\to 3} & 1 - p_{2\to 2} - p_{2\to 3} \\ 0 & 0 & p_{3\to 3} & 1 - p_{3\to 3} \\ 0 & 0 & 0 & 1 \end{pmatrix} \tag{B.3}$$

The zeros in this matrix indicate that we prevent any de-differentiation. In the following, we will also use the notation $p_{i,j}$ instead of $p_{i\to j}$.

In the absence of homogeneity in fate, i.e., if the two sister cells are independent in terms of their cell type choice, the probability of having a symmetric division (i.e., both daughter cells $k$ and $k'$ having the same type) for a mother cell of type $a \in \mathcal{P}$ is given by:

$$\mathbb{P}[p_k = p'_k | p_{m_k} = a] = \sum_{i=1}^{4} (p_{a\to i})^2$$

**We denote $H = 0$ for the hypothesis of an absence of homogeneity in fate**.
We want to contrast this hypothesis with the alternative hypotheses of the existence of homogeneity in fate, be it the same for all cells ($H = 1$) or different according to the mother cell type ($H = 2$) (see the two next paragraphs).

##### B.2.2 How to model Homogeneity in fate

Tak et al. [S12] have shown, from MultiGen experiments conducted on mouse stem and progenitor cells, a similarity in the cell types of cells from the same family, i.e., descending from a common ancestor, justifying the study of the hypothesis of homogeneity in fate in our model.

To model this homogeneity in fate, we introduce a dependency between the two sister cells in terms of their cell type "choice".
Let $k, k+1$ be two sister cells resulting from the division of a cell $m_k$ of type $p_{m_k} = a$.
We then define the joint law for their cell types, conditioning on the type of their mother cell, as follows:

$$\mathbb{P}[p_k = i, p_{k+1} = j | p_{m_k} = a] = \rho_{H,a} p_{a\to i} \delta_{i,j} + (1 - \rho_{H,a}) p_{a\to i} p_{a\to j} \tag{B.4}$$

With $\rho_{H,a} \in [0, 1]$ quantifying the importance of homogeneity in fate (according to the mother cell type $a$) and $\delta_{i,j}$ being 1 if $i = j$, 0 otherwise.

In terms of implementation, each sister cell has its type drawn according to the transition matrix (B.3). A random number is then drawn on $[0, 1]$, and if it is less than $\rho_{H,a}$, then the second sister's cell type (arbitrary choice) will be made identical to that of the first sister. Thus, the higher $\rho_{H,a}$, the greater this additional probability that both sisters are identical.

We verify that the marginal law of $p_k$ is the same as in the case without homogeneity. Indeed, we obtain it by integrating relation (B.4) with respect to $p_{k+1}$:

$$\begin{aligned} \mathbb{P}[p_k = i | p_{m_k} = a] &= \sum_{j=1}^{4} \mathbb{P}[p_k = i, p_{k+1} = j | p_{m_k} = a] \\ &= \rho_{H,a} p_{a\to i} + (1 - \rho_{H,a}) p_{a\to i} \sum_{j=1}^{4} p_{a\to j} \\ &= \rho_{H,a} p_{a\to i} + (1 - \rho_{H,a}) p_{a\to i} \\ &= p_{a\to i} \end{aligned}$$

In this model where a hypothesis of homogeneity in fate is made ($H \neq 0$), we now have a higher probability of having a symmetric division than in the case $H = 0$. Indeed, for $p_{m_k} = a \in \mathcal{P}$, the probability of symmetric division (both sister cell having the same type is equal to:

$$
\begin{aligned}
\mathbb{P}[p_k = p_{k+1} | p_{m_k} = a] &= \sum_{i=1}^{4} \mathbb{P}[p_k = i, p_{k+1} = i | p_{m_k} = a] \\
&= \sum_{i=1}^{4} \left(\rho_{H,a} p_{a\to i} + (1-\rho_{H,a}) p_{a\to i}^2\right) \\
&= \rho_{H,a} + (1-\rho_{H,a}) \sum_{i=1}^{4} p_{a\to i}^2 \\
&\geq \sum_{i=1}^{4} (p_{a\to i})^2
\end{aligned}
\tag{B.5}
$$

We verify that the limiting case $\rho_{H,a} = 0$ corresponds to the case without homogeneity in fate ($H = 0$). In the limiting case where $\rho_{H,a} = 1$, we see in relation (B.5) that sister cells will necessarily have the same cell type (but not necessarily the same as their mother).

In our model, the cell types of the sister cell pair depend only on that of their mother. We have notably not introduced a dependency on the type of a more distant ancestor, nor on the number of divisions undergone, nor on the time taken by the mother cell to divide. Thus, if we "forget" the dynamics of proliferation, i.e., the time taken between each division, and only consider the discrete extracted process (which corresponds to our differentiation process), then we obtain a multi-type Bienayme Galton-Watson process that could be analyzed analytically.
However, we also wish to study the proliferation process. We present in § B.3 our modeling proposal for this proliferation process.

##### B.2.3 Different scenarios of Homogeneity in fate

We denote $H \in \{0, 1, 2\}$ as the parameter indicating the homogeneity (in fate) scenario we are in. We denote $\rho_{H,a} \in [0, 1]$ as the parameter quantifying to which extent two sister cells would have the same type, when their mother is of type $a$. Remember, homogeneity in fate here is associated with an increased probability that two sister cells, after their mother's division (of a given type $a$), are of the same type (not necessarily the same as their mother).
The considered scenarios are:

- $H = 0$ corresponds to the scenario without homogeneity in fate ($\rho_{H,a} = 0\ \forall a$).
- $H = 1$ corresponds to the scenario with homogeneity in fate, independent of the mother's cell type $i$ ($\rho_{H,a} = \rho_{H,a'} > 0\ \forall a, a'$).
- $H = 2$ corresponds to the scenario where we made a distinguish between the mother cell types (HSC, MPP, and HPC) ($0 < \rho_{H,a} \neq \rho_{H_{a'}} \forall a, a', a \neq a'$). Note that sister cells derived from a CD34$^-$ will necessarily be homogeneous, as they will necessarily be CD34$^-$.

See also § B.4 which indicates the range of values which can be taken by the parameters for each scenario.

#### B.3 Proliferation

##### B.3.1 First Division

We introduce the random variable $T_1$ corresponding to the time of the first division, that is, the time it takes for the first (initial) cell to divide. If $N > 1$, then at least one division will occur and $T_1$ will correspond to the appearance time of cell 2: $T_{A,2}$ (as well as cell 3), as illustrated in Figure B.1.C. Otherwise, $T_{A,2}$ is not defined.

We consider that the times of first division in the model will be sampled directly from the empirical distribution (constructed from the observed times of first division, see § A.4). Since the empirical distribution is discrete, with observed times censored within intervals, and we want to simulate continuous times, we then add a value sampled uniformly from [0,1] to the sampled times.

##### B.3.2 Subsequent Divisions

All subsequent divisions will be modeled using lognormal distributions. Regarding the log-normal distribution, recall that a random variable $X \sim \mathcal{LN}(\mu, \sigma)$ if the variable $Y = \log(X)$ follows a normal distribution with mean $\mu$ and variance $\sigma^2$. The expectation and variance of $X$ then involve the parameters $\mu$ and $\sigma$ in their expression:

$$\mathbb{E}[X] = \exp(\mu + \sigma^2/2)$$
$$\mathbb{V}[X] = \left(e^{\sigma^2} - 1\right) e^{2\mu+\sigma^2}$$

The median will only depend on $\mu$ and will be $\exp(\mu)$.

For any cell of type $p_j$, its division time will follow a lognormal distribution:

$$T_j | p_j \sim \mathcal{LN}(\mu_{p_j}, \sigma_{p_j})$$

Where the median time is given by $\exp(\mu_{p_j})$.

Apart from the first division where the type of the mother cell is known, when studying subsequent divisions, this information is hidden. Indeed, as we modeled in section B.2.1, a mother cell by dividing will give two daughter cells each of which can have a type among several possibilities, according to different probabilities, which depend on the mother cell's type.
As seen in Fig. A.2, cells of different types have different distributions for their first division times. Not having any reason to consider it would be different for subsequent divisions, we then assume that the dynamics of proliferation, that is, the time required for a cell to divide, will depend on its type.
Let $j > 1$ be a cell of type $p_j$, we denote $T_j$ as the random variable corresponding to its division time (recall that we assume no cell death). $T_j$ will thus depend on $p_j$.
In a model without the hypothesis of concordance, denoted $C = 0$, we make no assumptions regarding other possible dependencies. In particular, we do not consider a dependency relation with the time taken by the mother cell $T_{m_j}$ to divide, with the time taken by its sister cell $T_{j'}$ (sister cell $j'$ such that $j' \neq j$ and $m_{j'} = m_j$) to divide or with the number of generations separating it from cell 1 (the initial cell).

We then model $T_j$ using a log-normal distribution whose parameters are specific to the cell type $p_j \in \mathcal{P}$: $T_j | p_j \sim \mathcal{LN}(\mu_{p_j}, \sigma_{p_j})$. We choose to use only the family of log-normal distributions, regardless of the type of the dividing cell. We could also have chosen a gamma distribution, which would have been a relevant model for the number of phases a cell encounters during division. However, we prefer here the family of log-normal distributions, also used by other authors for modeling the division times of hematopoietic cells [S4, S7, S8, S10], and which presents a practical-to-use multivariate version (for modeling concordance, see following paragraph) implemented in the Julia language.

##### B.3.3 How to model Concordance

We have presented above a description of the proliferation process under the hypothesis $C = 0$ of absence of concordance. However, many results tend to show a strong correlation in the division times of sister cells (but not with that of the mother cell). We denote $C \neq 0$ the hypothesis that there is concordance between sister cells. In this paragraph, we introduce how we model concordance between sister cells. We will also potentially consider concordance between cousins cells, as we will present it in the next paragraph, and also more complex hypotheses of concordance ($C \geq 2$).

Let $k, k'$ be two sister cells, with their division times $T_k$ and $T_{k'}$ modeled as random variables. Under the hypothesis $C = 0$, the variables $T_k | p_k$ and $T_{k'} | p_{k'}$ were independent, distributed according to log-normal laws with parameters $(\mu_{p_k}, \sigma_{p_k})$ and $(\mu_{p_{k'}}, \sigma_{p_{k'}})$ respectively.

We now introduce a dependency relation between these two variables while ensuring that their marginal laws remain the same as before:

$$(T_k, T_{k'}) | p_k, p_{k'} \sim MV\text{-}\mathcal{LN}\left((\mu_{p_k}, \mu_{p_{k'}}), \mathbf{\Sigma}\right) \quad \text{(B.6)}$$

where $MV\text{-}\mathcal{LN}$ corresponds here to a bivariate log-normal distribution with:

$$\mathbf{\Sigma} = \begin{pmatrix} \sigma^2_{p_k} & \rho_C \sigma_{p_k} \sigma_{p_{k'}} \\ \rho_C \sigma_{p_k} \sigma_{p_{k'}} & \sigma^2_{p_{k'}} \end{pmatrix}$$

and $\rho_C \in [0, 1]$ a parameter quantifying the importance of concordance.
With the hypothesis denoted as $C = 1$, we model that there is concordance between all sister cells, even of different cell types, and that concordance (i.e., the parameter $\rho_C$) does not depend on the cell type. In particular, with this model, two sister CD34$^-$ cells would remain concordant. However, this population of cells is very heterogeneous, potentially containing cells already committed to certain cellular lineages, making the hypothesis of concordance between CD34$^-$ cells potentially unjustified. In the next paragraph, we present different scenarios of concordance we have considered.

##### B.3.4 Concordance between cousin cells and different scenarios of concordance

We denote $C \in \{0, 1, \cdots, 5\}$ as the parameter indicating the concordance scenario (between sisters) we are in. Concordance between cousins may also be considered. During simulations, we will always consider the division of four cousins together, extending what we presented in the previous paragraph, as follows:

$$(T_{i_1}, T_{i_2}, T_{i_3}, T_{i_4})|i_1, i_2, i_3, i_4 \sim MV\text{-}\mathcal{LN}\left((\mu_{i_1}, \mu_{i_2}, \mu_{i_3}, \mu_{i_3}), \mathbf{\Sigma}\right)$$

with:

- indices 1 and 2 referring to the first pair of sisters and 3 and 4 to the second pair,
- $i_1$ corresponding to the type of sister cell 1 that divides at $T_{i_1}$,
- $MV\text{-}\mathcal{LN}$ representing a multivariate log-normal distribution with:

$$\mathbf{\Sigma} = \begin{pmatrix} \sigma^2_{i_1} & \rho_{s,1,2}\ \sigma_{i_1}\sigma_{i_2} & \rho_{C,c}\ \rho_{C,m}\ \sigma_{i_1}\sigma_{i_3} & \rho_{C,c}\ \rho_{C,m}\ \sigma_{i_1}\sigma_{i_4} \\ \rho_{C,1,2}\ \sigma_{i_2}\sigma_{i_1} & \sigma^2_{i_2} & \rho_{C,c}\ \rho_{C,m}\ \sigma_{i_2}\sigma_{i_3} & \rho_{C,c}\ \rho_{C,m}\ \sigma_{i_2}\sigma_{i_4} \\ \rho_{C,c}\ \rho_{C,m}\ \sigma_{i_3}\sigma_{i_1} & \rho_{C,c}\ \rho_{C,m}\ \sigma_{i_3}\sigma_{i_2} & \sigma^2_{i_3} & \rho_{C,3,4}\sigma_{i_3}\sigma_{i_4} \\ \rho_{C,c}\ \rho_{C,m}\ \sigma_{i_4}\sigma_{i_1} & \rho_{C,c}\ \rho_{C,m}\ \sigma_{i_4}\sigma_{i_2} & \rho_{C,3,4}\ \sigma_{i_4}\sigma_{i_3} & \sigma^2_{i_4} \end{pmatrix}$$

In the covariance matrix above:

- $\rho_{C,1,2}$ and $\rho_{C,3,4}$ correspond to concordance parameters associated with sister pairs (1,2) and (3,4), respectively.
- $\rho_{C,m} = \min(\rho_{C,1,2}, \rho_{C,3,4})$.
- $\rho_{C,c}$ corresponds to concordance between cousins. By multiplying $\rho_{C,c}$ by the minimum of $\rho_{C,1,2}$ and $\rho_{C,3,4}$, we ensure that concordance between cousins is always weaker than between sisters (otherwise, there might be a risk of the matrix not being positive definite).

The value of $\rho_{C,1,2}$ (similarly for $\rho_{C,3,4}$) depends on the scenario:

- $C = 0$: No Concordance: $\rho_{C,1,2} = 0$.
- $C = 1$: Identical concordance regardless of the cell pair: $\rho_{C,1,2} = \rho_{C,A}$.
- $C = 2+$, i.e, $C \in \{2, 3, 4, 5\}$: More complex models with concordance under certain conditions:
  - $C = 2$: Concordance only between sister cells of the same type: $\rho_{C,1,2} = \rho_{C,A}$ if $i_1 = i_2$, 0 otherwise.
  - $C = 3$: Different concordance if sister cells are of different types: $\rho_{C,1,2} = \rho_{C,A}$ if $i_1 = i_2$, $\rho_{C,1,2} = \rho_{C,B}$ otherwise.
  - $C = 4$: Concordance among all sister cells, except when at least one is CD34$^-$: $\rho_{C,1,2} = \rho_{C,A}$ if $i_1 \neq 4$ and $i_2 \neq 4$, 0 otherwise.
  - $C = 5$: Different concordance if one of the sisters is CD34$^-$: $\rho_{C,1,2} = \rho_{C,A}$ if $i_1 \neq 4$ and $i_2 \neq 4$, $\rho_{C,1,2} = \rho_{C,B}$ otherwise.

With $\rho_{C,A}$ and $\rho_{C,B}$ being the concordance parameters to be estimated.
See also § B.4 which indicates the range of values which can be taken by the parameters for each scenario.

Function `construct_cov_mat` in `/dynamic_model/support_functions.jl` details the construction of the covariance matrix $\mathbf{\Sigma}$.

#### B.4 Priors

The table B.1 summarizes the parameters used in the model along with their *a priori* distributions. Some parameters have joint priors sometimes defined implicitly. The particular cases mentioned in the table are listed below:

($*^1$) $p_{1,1}$, $p_{1,2}$, and $p_{1,3}$ are each drawn uniformly from $[0, 1]$, until the condition $p_{1,1} + p_{1,2} + p_{1,3} \leq 1$ is met.

($*^2$) $p_{2,2}$ and $p_{2,3}$ are each drawn uniformly from $[0, 1]$, until the condition $p_{2,2} + p_{2,3} \leq 1$ is met.

($*^3$) Different scenarios of homogeneity in fate are considered:

- If $H = 0$: no homogeneity, $\rho_{H,1} = \rho_{H,2} = \rho_{H,3} = 0$.
- If $H = 1$: Homogeneity in fate independent of type, $\rho_{H,1} \sim \mathcal{U}([0.3, 1])$ and $\rho_{H,2} = \rho_{H,3} = \rho_{H,1}$.
- If $H = 2$: different values of the parameter depending on the mother cell type . $\rho_{H,1}$, $\rho_{H,2}$, and $\rho_{H,3}$ are each drawn uniformly from $[0, 1]$, until one of both following conditions is met:
  - $\mid \rho_{H,1} - \rho_{H,2} \mid + \mid \rho_{H,1} - \rho_{H,3} \mid + \mid \rho_{H,2} - \rho_{H,3} \mid \geq 0.3$.
  - $\rho_{H,1} + \rho_{H,2} + \rho_{H,3} \geq 0.3$.

($*^4$) $\rho_{C,B}$ is drawn uniformly from $[0, 1]$ until $\mid \rho_{C,B} - \rho_{C,A} \mid \geq 0.2$ is achieved.

($*^5$) $C$ takes its values in $\{0, 1, \cdots, 5\}$ with the associated probabilities, respectively, 1/3, 1/3, 1/12, 1/12, 1/12, 1/12. This choice is made considering three main concordance scenarios, *a priori* equally probable: no concordance ($C = 0$, in which case parameters $\rho_{C,A}$ and $\rho_{C,B}$ will not be used), identical concordance for all types ($C = 1$, in which case $\rho_{C,B}$ is not used), or concordance associated with certain conditions ($C \geq 2$). Hence, for this last case, values 2, 3, 4, and 5 correspond to sub-scenarios, hence the choice of an *a priori* probability of 1/12.

($*^6$) $\rho_{C,c}$ follows a prior distribution such that favoring both extrem cases, either concordance between cousins ($\rho_{C,c} = 1$) or not ($\rho_{C,c} = 0$).

The prior distribution of the vector parameter is implemented in `/stat_model/prior.jl`.

| Parameter | Label | Meaning | Prior |
|---|---|---|---|
| $p_{1,1}$ | p1_1 | Transition probability from type 1 (HSC) to 1 | $\in [0,1]$ (*1) |
| $p_{1,2}$ | p1_2 | Transition probability from type 1 (HSC) to 2 (MPP) | $\in [0,1]$ (*1) |
| $p_{1,3}$ | p1_3 | Transition probability from type 1 (HSC) to 3 (HPC) | $\in [0,1]$ (*1) |
| $p_{1,4}$ | p1_4 | Transition probability from type 1 (HSC) to 4 (CD34-) | $= 1 - p_{1,1} - p_{1,2} - p_{1,3}$ |
| $p_{2,2}$ | p2_2 | Transition probability from type 2 (MPP) to 2 | $\in [0,1]$ (*2) |
| $p_{2,3}$ | p2_3 | Transition probability from type 2 (MPP) to 3 (HPC) | $\in [0,1]$ (*2) |
| $p_{2,4}$ | p2_4 | Transition probability from type 2 (MPP) to 4 (CD34-) | $= 1 - p_{2,2} - p_{2,3}$ |
| $p_{3,3}$ | p3_3 | Transition probability from type 3 (HPC) to 3 | $\sim \mathcal{U}([0,1])$ |
| $p_{3,4}$ | p3_4 | Transition probability from type 3 (HPC) to 4 (CD34-) | $= 1 - p_{3,3}$ |
| $\mu_1$ | mu_1 | $\exp(\mu_1)$ is the median of the HSC division time | $\sim \mathcal{U}([1.5, 4.0])$ |
| $\sigma_1$ | sig_1 | associated to the variability of the HSC division time | $\sim \mathcal{U}([0, 0.5])$ |
| $\mu_2$ | mu_2 | $\exp(\mu_2)$ is the median of the MPP division time | $\sim \mathcal{U}([1.5, 4.0])$ |
| $\sigma_2$ | sig_2 | associated to the variability of the MPP division time | $\sim \mathcal{U}([0, 0.5])$ |
| $\mu_3$ | mu_3 | $\exp(\mu_3)$ is the median of the HPC division time | $\sim \mathcal{U}([1.5, 4.0])$ |
| $\sigma_3$ | sig_3 | associated to the variability of the HPC division time | $\sim \mathcal{U}([0, 0.5])$ |
| $\mu_4$ | mu_4 | $\exp(\mu_4)$ is the median of the CD34- division time | $\sim \mathcal{U}([1.5, 4.0])$ |
| $\sigma_4$ | sig_4 | associated to the variability of the CD34- division time | $\sim \mathcal{U}([0, 0.5])$ |
| $\rho_{H,1}$ | rho_H_1 | Homogeneity in fate associated with mother cells of type 1 (HSC) | $\in [0,1]$ (*3) |
| $\rho_{H,2}$ | rho_H_2 | Homogeneity in fate associated with mother cells of type 2 (MPP) | $\in [0,1]$ (*3) |
| $\rho_{H,3}$ | rho_H_3 | Homogeneity in fate associated with mother cells of 3 (HPC) | $\in [0,1]$ (*3) |
| $\rho_{H,4}$ | rho_H_4 | CD34- are necessarily homogeneous | = 0 (or any other value) |
| $H$ | H | Scenario of homogeneity in fate | $\sim \mathcal{U}(\{0,1,2\})$ (*3) |
| $\eta$ | recovery_rate | Recovery (sampling) rate | $\sim \mathcal{U}([0.4, 0.9])$ |
| $\rho_{C,A}$ | rho_C_sisters_A | associated with the concordance between sister cells, if applicable | $\sim \mathcal{U}([0.3, 1.0])$ |
| $\rho_{C,B}$ | rho_C_sisters_B | associated with the concordance between sister cells, if applicable | $\in [0,1]$ (*4) |
| $C$ | C | Scenario of concordance | $\in \{0,1,2,3,4,5\}$ (*5) |
| $\rho_{C,c}$ | rho_C_cousins | Concordance between cousins, if applicable | $\sim$ Beta(0.5, 0.5) (*6) |

Table B.1: List of parameters used in the model and its implementation, for which the *prior* is indicated (when the parameter is to be estimated). Specific conditions (*) are specified in the text. $\sim \mathcal{U}$ indicates a uniform draw on the set specified as an argument. Label corresponds to the notation used in the implementation.

### C Observation model

For a fixed set of parameters, it is possible to simulate the proliferation and differentiation process as many times as desired, starting from a cell of a given type, and stopping at an observation time of $T_{max} = 72$ or 96 hours. The simulation provides access to the entire system's information at any given time. However, what can be experimentally observed is only a part of the system, either the times of the first divisions or the system at $T_{max}$, with additional noise (interval censoring for the times of the first divisions, sampling for the observation at $T_{max}$).
The experimental observations available are numerous and highly variable. As such, they cannot be used as-is for comparison with the outputs of the simulations. Therefore, the comparison of model outputs to experimental data requires the use of descriptive statistics (summary statistics), which aggregate information from different wells (different families) to create summaries (calculation of mean, variance, proportions, etc.). The idea is that while two families may exhibit a lot of variability, making their comparison challenging, when comparing quantities obtained by aggregating a large number of wells, these quantities are relatively stable, making their pairwise comparison easier and more relevant.

#### C.1 Observation variables

The state of our system (or process) is entirely characterized by $N$, $\boldsymbol{p}$, $\boldsymbol{m}$, and $\boldsymbol{T_A}$, random variables (and vectors) for which we have specified the dependency relations previously, in section B. However, these state variables are not directly observable: we will only have access to the state of the system through observation variables.

The objective, through observations, is to understand the behavior of the system, that is - since we have proposed a parametric model - to estimate the values of the parameters. We will estimate the model parameters considering that we observe multiple random realizations (considered independent and identically distributed) of our stochastic process. Recall that the system is described at the scale of a family originating from a single cell whose type at $t = 0$ (initial condition) is known. We will thus observe several families. A family may either be observed by the Live Cell Imaging experiment or by the MultiGen experiment. We index our observations by $i \in \mathcal{I}$ and $j \in \mathcal{J}$ with $\mathcal{I}$ and $\mathcal{J}$ corresponding to the set of $N_{\mathcal{I}}$ and $N_{\mathcal{J}}$ families observed by the Live Cell Imaging and MultiGen experiments, respec-

tively. The initial condition is assumed to be known without uncertainty, that is, we have information about the type of cell 1 ($p_1$) and assume $T_A = 0$, which amounts to neglecting anything that may have influenced the fate of the cell before the start of the experiment.
We will then denote $\mathcal{I} = \bigcup_{p\in\{1,2,3\}} \mathcal{I}_p$ (and similarly for $\mathcal{J}$), differentiating the observed families depending on whether they originate from an HSC, MPP, or HPC. Note that we do not have observations starting with a CD34$^-$ cell.

If the system is observed via the Live Cell Imaging experiment, our observation variables will be $D_1$, $D_{2,1}$, and $D_{2,2}$, and $D_{3,1}$, $D_{3,2}$, $D_{3,3}$, and $D_{3,4}$, which correspond to the lower bounds of the intervals during which the first division ($T_1$), the second divisions $T_{2,1}$ and $T_{2,2}$, and the third divisions $T_{3,1}$, $T_{3,2}$, $T_{3,3}$, and $T_{3,4}$ took place. As the observations are also right-censored due to the observation time limit $T_{max}$, if the division occurred after this time, these variables would be assigned the value $T_{max}$ (which is the lower bound of the interval $]T_{max}, +\infty]$). Apart from censorship, we do not consider uncertainties related to the measurement of these division times. We also observe $N(T_{max}) = \text{card}(\mathcal{C}_{T_{max}})$, that is, the number of cells present at $T_{max}$, which we consider estimated without uncertainty.

If the system is observed via the MultiGen experiment, it will then be observed at $t = T_{max}$. There is sampling noise: we only have information for a set of cells $\hat{\mathcal{C}}_{T_{max}} \subset \mathcal{C}_{T_{max}}$. We hypothesize that:

$$\mathbb{P}[i \in \hat{\mathcal{C}}_{T_{max}} | i \in \mathcal{C}_{T_{max}}] = \eta \tag{C.1}$$

with $\eta \in [0, 1]$ called the sampling rate (recovery rate), assumed to be constant and independent of $\text{card}(\mathcal{C}_{T_{max}}) = N(T_{max})$.
For each sampled cell $k \in \hat{\mathcal{C}}_{T_{max}}$, we observe the random variables corresponding to its type $p_k$ and its number of divisions $n_k$ counted from the initial cell, with $n_1 = 0$ (corresponding to generation 1) (see Fig. A.5). With the dilution of fluorescence markers, beyond a certain number of divisions, we consider that we can no longer discern them. Any cell beyond generation 7 (which corresponds to the maximum generation observed in our dataset) will then be assigned to this generation. We do not consider other uncertainties in these observations.

Finally, the uncertainties related to observations are associated either with interval censoring in the case of Live Cell Imaging data or with sampling in the case of MultiGen.
From the above observation variables and their $N_{\mathcal{I}}$ and $N_{\mathcal{J}}$ realizations, we will next seek to estimate which scenario is the most likely.
Given the high number of different realization of the experiments, the observations will actually be synthetised by summary statistics, as detailled in the next paragraph.

#### C.2 Summary Statistics

At the level of a family (derived from a single cell of a known cell type), we can for example observe:

- The number of different generations
- The difference between the third and second division times
- The number of cells observed in generation 3 at $T_{max}$
- The colony size at $T_{max}$
- Whether the cells are distributed over exactly 2 different generations
- Whether the cells are distributed over 2 different cell types
- Whether the cells are only in the third generations
- etc.

Then, aggregating these results over all families (for a given initial cell type and observation time), we get what we call summary statistics:

- The mean number of generations observed
- The mean difference between the third and second division times
- The mean number of cells observed at generation 3 for a given $T_{max}$ (either 72 or 96h)
- The mean colony size for a given $T_{max}$
- The proportion of families whose cells are exactly distributed over 2 different generations
- The proportion of families whose cells are exactly distributed over 2 different cell types
- The proportion of families whose cells are only in generation 3
- etc.

Summary statistics will then be defined separately for:

- Live Cell Imaging experiment-related observations (e.g., average times of second division, average colony size at 96 hours, etc.). There will be 25 such statistics.
- MultiGen experiment-related observations (e.g., average number of cells observed in generation 2, proportion of families covering exactly 2 cell types at the observation time, etc.). There are 53 of these statistics.

The Live Cell Imaging-related summary statistics will then be calculated in three different configurations, depending on the starting cell type (start), resulting in $3 \times 25 = 75$ statistics.
The MultiGen-related summary statistics will be calculated in 5 different configurations, depending on $T_{max} \in \{72, 96\}$ and the starting type (noting that there are no observations at 72 hours starting from an HPC), resulting in $5 \times 53 = 265$ statistics, for a total of 340.
These 340 summary statistics can be calculated based on real observations. They can also be calculated from a simulation of the system for a given set of parameters. In what follows, the term "simulation" will be reserved (as much as possible) for simulating the proliferation and differentiation process, with the same set of parameters, for a set of families (avoiding the use of the term simulation for just one well).
Thus, the summary statistics can be calculated in the same way, whether from a model simulation or from experimental observations (in particular, using the same number of wells/families for their calculation). This approach allows for a consistent and comparable way to evaluate and compare the model's output against experimental data.

The summary statistics are listed and implemented in the function:
`/stat_model/summary_statistics.jl`.

Summary statistics are used to reduce the number of observations. Yet, 340 summary statistics remains a high number of observations, and it is, therefore, not feasible to try looking for a parameter vector which would lead to 340 summary statistics when, computed from the simulation, would be all close to the ones computed from the experimental observations.
Therefore, we still need to reduce the number of our observations variables. This is done by using a standard dimension reduction technique, namely a principal component analysis, as presented in the next paragraph.

#### C.3 Principal Component Analysis (PCA)

Given the large number of summary statistics, it is impractical to use them directly for model calibration. Ideally, we would seek parameter values that, when used to simulate the system, produce summary statistics from the simulation that closely match those calculated from experimental data. However, this is not feasible in practice, as the model is necessarily a simplification of reality. A compromise is needed, which involves defining a single distance combining the different summary statistics. The question arises as to whether certain summary statistics are more informative or relevant than

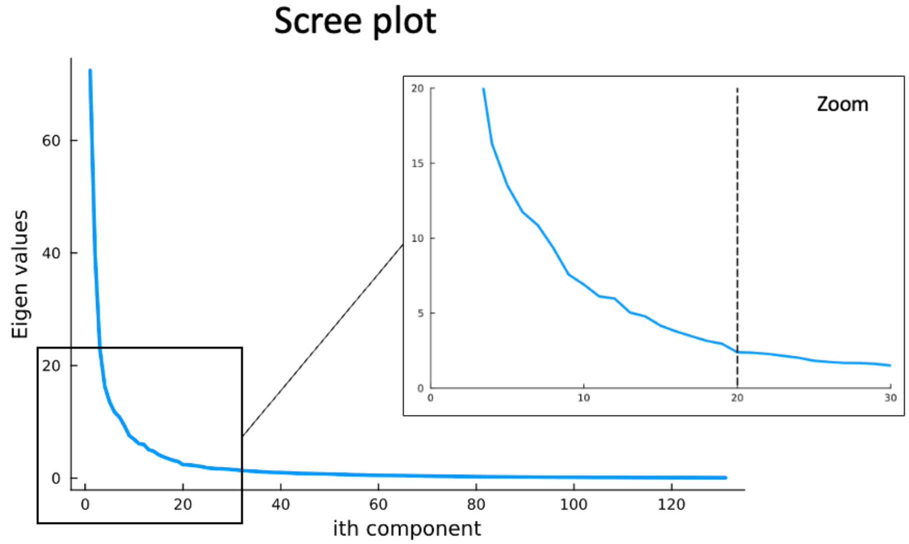

Figure C.1: Scree plot of eigenvalues (indicating the variance of the dataset explained by each component). Components are ranked by decreasing importance. We choose to consider up to the 20th principal component.

others. To avoid a selection bias based on biological *a priori*, we choose to construct a reduced set of new summary statistics as linear combinations of the initial 340 summary statistics. This reduced set construction is based on Principal Component Analysis (PCA), a standard method for dimension reduction.
To perform PCA, we generate a synthetic dataset of size 122,495 × 340, simulating our system 122,495 times, each time with a set of parameters drawn according to the prior, and calculating our 340 summary statistics for each simulation.

The summary statistics are then normalized (mean and standard deviation calculated for each, which will be retained for future normalization). The resulting distributions for the summary statistics are thus centered and reduced.

Executing PCA, we find that 99% of the variance of the synthetic dataset is explained by 131 components. However, we wish to retain fewer components. The scree plot (Figure C.1) helps determine the number of components to keep. An heuristic is to stop at the "elbow" on this plot, i.e., where adding more components does not significantly increase the explained variance. This elbow seems to be at the 20th component.
Another heuristic is to consider components until 80% of the dataset's variance is explained. With 20 components, we achieve a principal ratio of 0.7995. This choice satisfies both criteria.
Thus, we obtain a dimension reduction matrix, allowing us to convert a vector of size 340 (corresponding to our 340 summary statistics) to 20 components, which will then be used for model selection.

#### C.4 Contribution of the Summary Statistics

To understand how the principal components are constructed and how they combine the initial summary statistics, we can examine their contributions. For each of the initial 340 summary statistics, we have its weight (in absolute value) within each component. We can then deduce the percentage contribution of each summary statistic to a given component. By considering the 20 components we retained (and weighting them), we can deduce the overall contribution (in percentage) of each summary statistic. Figure C.2 represents the contributions in ascending order. Some statistics have zero contribution; for instance, those that will always have values equal to 0, such as the statistic indicating the number of HSC cells when starting from an MPP (since de-differentiation is not allowed). Such summary statistics could have been discarded a priori, there are now automatically discarded as a result of the PCA analysis.

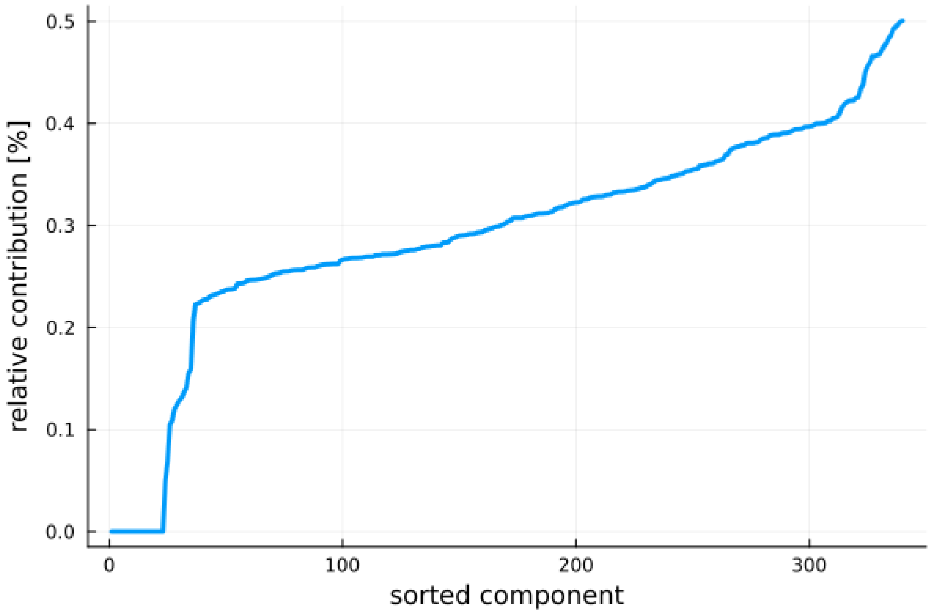

Figure C.2: For each initial descriptive statistic, its contribution (in %) to the 20 selected components is calculated and plotted in ascending order.

Table C.1 lists the 25 summary statistics with the highest contributions. The whole list is available in `/simulate/pca/contribution_sum_stats.csv`.
These statistics are the most likely to allow discrimination between parameters/scenarios.

#### C.5 Distance

Ultimately, a distance needs to be defined to compare experimental observations with simulations. This distance is chosen as the L2 norm on the 20 principal components obtained after PCA. In practical terms, this means calculating the sum of squared errors for each of the components, comparing the value obtained from simulations with the value obtained from experiments.
The L2 norm is a common choice in statistical modeling for measuring the difference between two sets of values. In this context, it serves to quantify the discrepancy between the simulated data (as per the proposed model) and the real experimental data. A smaller L2 norm indicates a closer match between the model and the observed data, suggesting a better model fit.
By focusing on the principal components, this approach effectively weighs the contributions of different aspects of the data according to their variance and importance as determined by the PCA. This weighting ensures that the most significant features of the data, as captured by the principal components, play a larger role in determining the overall distance.

| Label | Contribution [%] |
| --- | --- |
| mean_nb_cells_type_i_2_startMPP_obs96 | 0.5 |
| std_T2_2_startHSC | 0.49 |
| std_nb_cells_type_i_2_startHSC_obs96 | 0.49 |
| std_T2_startHSC | 0.49 |
| std_nb_cells_type_i_2_startMPP_obs96 | 0.49 |
| std_T2_2_startMPP | 0.48 |
| std_T2_startHPC | 0.48 |
| mean_nb_cells_type_i_1_startHSC_obs72 | 0.47 |
| prop_families_only_in_type_i_4_startHPC_obs96 | 0.47 |
| std_nb_cells_type_i_1_startHSC_obs72 | 0.47 |
| std_nb_cells_type_i_2_startMPP_obs72 | 0.46 |
| mean_nb_cells_type_i_2_startHSC_obs96 | 0.46 |
| std_T2_2_startHPC | 0.46 |
| std_T2_1_startHSC | 0.46 |
| std_diff_T2_2_T1_startHSC | 0.45 |
| std_T2_startMPP | 0.45 |
| std_T2_1_startHPC | 0.45 |
| std_diff_T2_2_T1_startHPC | 0.43 |
| std_T2_1_startMPP | 0.43 |
| mean_nb_cells_type_i_2_startMPP_obs72 | 0.42 |
| std_diff_T2_1_T1_startHPC | 0.42 |
| std_coverage_type_startMPP_obs96 | 0.42 |
| prop_fam_over_n_gen_4_startHPC_obs96 | 0.42 |
| std_diff_T2_2_T1_startMPP | 0.42 |
| prop_fam_over_n_gen_1_startHPC_obs96 | 0.42 |

Table C.1: List of the 25 descriptive statistics contributing the most to the 20 components considered. Labels correspond to those returned by function get_labels_sum_stats in /stat_model/summary_stats.jl.

### D Estimating the posterior probability of each of our main scenarios

The four main scenarios we want to compare are the following:

- $(H = 0; C = 0)$ (No homogeneity in fate; no concordance in division)
- $(H = 0; C = 1)$
- $(H = 1; C = 0)$
- $(H = 1; C = 1)$

The four scenarios are considered (a priori) equiprobable.
Here, we do not consider any concordance between cousin cells, that is, $\rho_{C,c} = 0$.
Otherwise, the prior distributions are the same as presented in § B.4 (and in Tab. B.1).

Scenarios where $H = 1$ differ from those where $H = 0$ by an additional parameter $\rho_H := \rho_{H,1} = \rho_{H,2} = \rho_{H,3} \in [0.3, 1.0]$ (the three parameters can take different values only in the more complex scenario $H = 2$ which is studied in Appendix E).
Scenarios where $C = 1$ differ from those where $C = 0$ by an additional parameter $\rho_C := \rho_{C,A} \in [0.3, 1.0]$ to estimate (here, $\rho_{C,B} = 0$, this parameter being used only for more complex scenarios of concordance, as studied in Appendix E).

#### D.1 Using an ABC rejection method

First, we use an ABC rejection sampling method to estimate the posterior probability of each of our four main scenarios [33, 34].
We sample 1,773,106 times a parameter vector from the prior distribution (we consider the indexes of the hypotheses $H$ and $C$ as discrete parameters), and for each parameter vector, simulate the model (that is, simulate as many families as we have in our experimental dataset) to compute our summary statistics and our distance to the observations (see Fig. D.1).

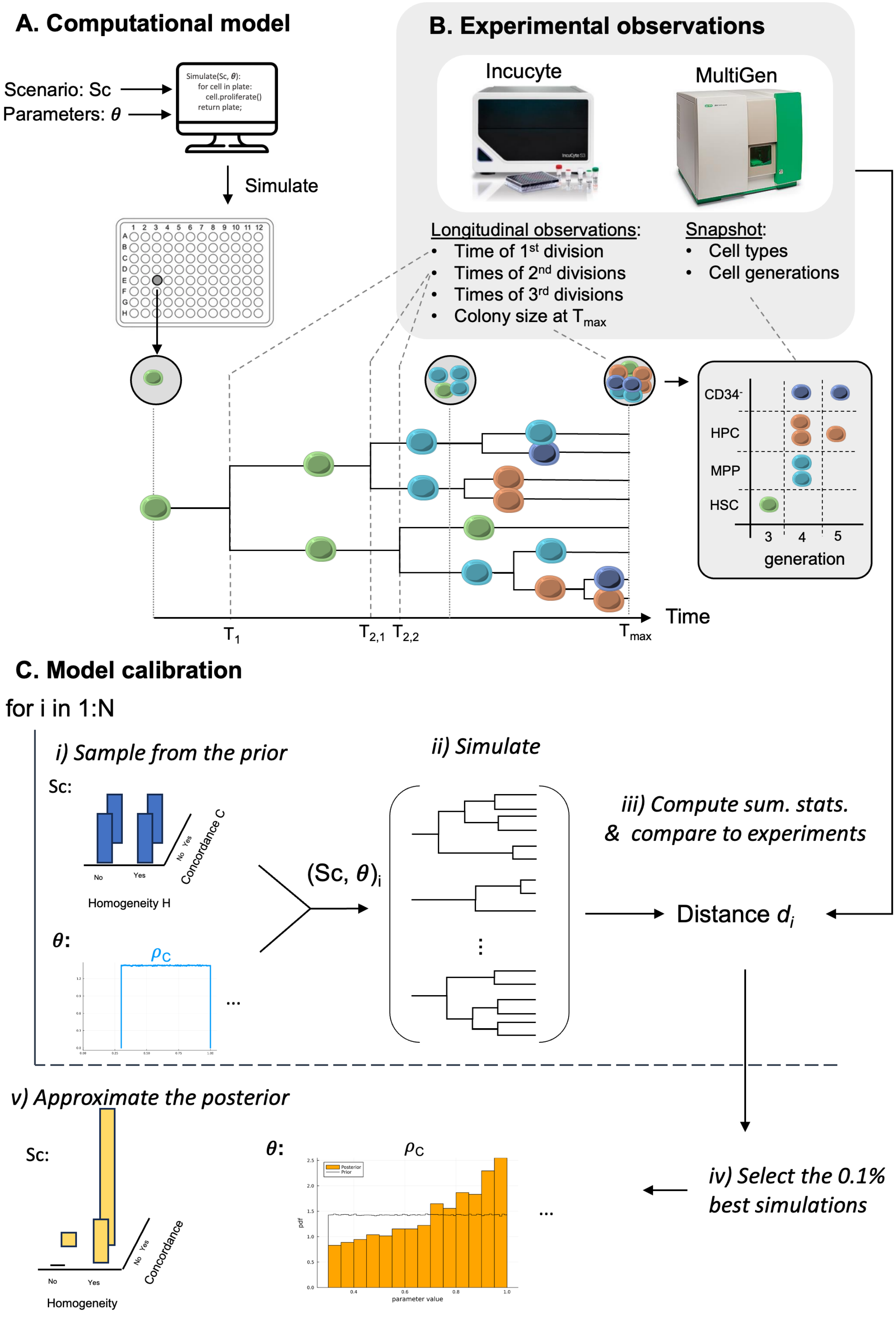
A. Computational model
Scenario: Sc
Parameters: θ
Simulate(Sc, θ):
for cell in plate:
cell.proliferate()
return plate;
Simulate
B. Experimental observations
Incucyte
MultiGen
Longitudinal observations:
Time of 1st division
Times of 2nd divisions
Times of 3rd divisions
Colony size at Tmax
Snapshot:
Cell types
Cell generations
CD34-
HPC
MPP
HSC
3
4
5
generation
Time
T1
T2,1
T2,2
Tmax
C. Model calibration
for i in 1:N
i) Sample from the prior
Sc:
No
Yes
Homogeneity H
Concordance C
θ:
ρC
ii) Simulate
(Sc, θ)i
iii) Compute sum. stats. & compare to experiments
Distance di
iv) Select the 0.1% best simulations
v) Approximate the posterior
Sc:
Homogeneity
Concordance
θ:
ρC
Posterior
Prior
pdf
parameter value

Figure D.1: Illustration of the methodology for estimating the posterior probability of each of our four main scenarios, using an ABC-Rejection sampler.
A) We have an implementation of our computational model, that we can instantiate using as input the scenario to consider and the parameter values $\boldsymbol{\theta}$. Once instantiate, we can simulate the model, that is, compute the proliferation and differentiation process for many wells (many families or colonies). For each family, we can retrieve - from the simulations - the observations variables which can be associated to what we observe from the Live Cell Imaging (Incucyte©) and MultiGen assays (B).
C) To calibrate the model and estimate the posterior probabilities of each of our model, we repeat $N$ times (here, $N = 1,773,106$) the following procedure. i) We sample from the prior the scenario ($H = 0$ or 1; $C = 0$ or 1), and, given the scenario, parameter values. ii) We simulate, using these values, as many dynamics as we have experimental families. iii) From this large set of dynamics, we compute our summary statistics, and by comparing the ones based on the computations to those based on the experiment, we compute a distance to the observation. After having repeated this procedure $N$ times, we end up with a set of $N$ parameter vectors (and scenarios) associated with distances. We select the 0.1% best distances. The parameter vectors (and scenarios) for which the distance was among the 0.1% best ones are used to approximate the posterior distributions.

We keep only the parameter vectors for which the computed distance to the observation was low enough, below a threshold $\epsilon$. This threshold is chosen as the 0.1 percentile based on the 1,773,106 distances that were computed (see Fig. D.2).

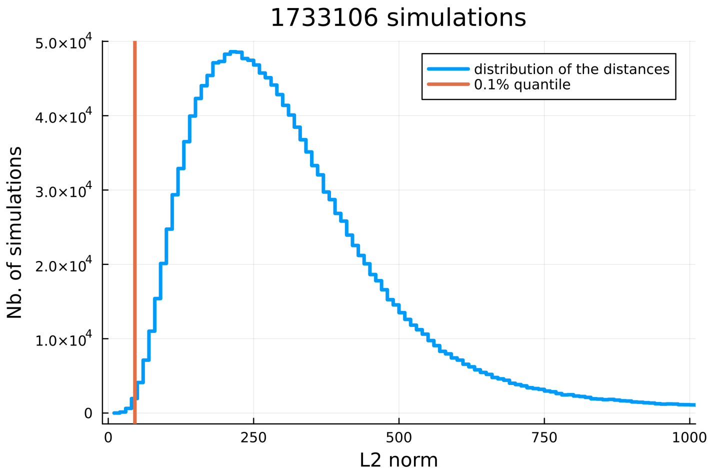

Figure D.2: Distribution of distances to experimental observations across the $\sim$ 2 millions simulations. The orange vertical line indicates the tolerance associated with the 0.1% percentile. Scenarios (and parameters) leading to simulations with a distance less than this tolerance (here, equal to 45.83) will be retained to approximate the posterior probabilities of each scenario. Only one in a thousand simulations is thus retained.

Then, among the 1,734 parameter vectors selected, we look at the frequency of each of our 4 models, these frequencies approximating the posterior distribution of each of scenario. We find:

- $\mathbb{P}[H = 1, C = 1|\mathcal{D}] \approx 70.70\%$
- $\mathbb{P}[H = 1, C = 0|\mathcal{D}] \approx 22.32\%$
- $\mathbb{P}[H = 0, C = 1|\mathcal{D}] \approx 6.92$
- $\mathbb{P}[H = 0, C = 0|\mathcal{D}] \approx 0.06\%$

The scenario where both the hypotheses of homogeneity in fate and concordance in division appears to be the most likely.
The scenario where both hypotheses would be absent is clearly rejected.
There is a slight probability, around 0.22, that the hypothesis of homogeneity in fate alone would be sufficient to explain our observations.

ABC rejection methods require a huge number of simulations for the algorithms to converge.
In the next paragraphs, we explore an other approach to refine our results, and use an ABC-SMC method.

#### D.2 Using an ABC-SMC method

We consider $N = 2,000$ parameter vectors, that we call particles.
In the previous paragraph, over the 1,773,106 simulations, the best $N$ parameter vectors led to a distance to the simulation below 41.64.
Using a sequential approach, we will progressively decrease the tolerance, ensuring to always have $N$ particles whose distance to the observations are below the threshold of the given step.

We implemented the ABC-SMC sampler as presented in [70] (see Algorithm 1).
The ABC-SMC algorithm is implemented in `/ABC_SMC/ABC_SMC.jl`.

**Algorithm 1** ABC-SMC Algorithm (adapted from Toni et al. [70])

Set $N$ the number of particles (parameter vectors)
Set $N_i$ the number of iterations
Initialize a tolerance sequence $\{\epsilon_i\}_{i=1}^{N_i}$ with $\epsilon_0 > \epsilon_1 > \cdots > \epsilon_{N_i} > 0$.
Initialize a sequence $\{B_i\}_{i=1}^{N_i}$ where $B_i$ corresponds to the number of time we run the model (and compute the distance), for a given particle, at iteration $i$.
Initialize a sequence $\{K_i\}_{i=1}^{N_i}$ of perturbation kernels
For iteration 0, sample $\{\boldsymbol{\theta}_0^{(j)}\}_{j=1}^{N}$ parameter vectors (particles) from the prior and set the corresponding particle weights $\{w_0^{(j)}\}_{j=1}^{N}$ to $1/N$.
**for** $i = 1$ to $N_i$ **do**
  **for** $j = 1$ to $N$ **do**
    $b_i^{(j)} \leftarrow 0$
    **while** $b_i^{(j)} = 0$ **do**
      Sample $\boldsymbol{\theta}^*$ from the previous population $\{\boldsymbol{\theta}_{i-1}^{(k)}\}_{k=1}^{N}$ with weights $\{w_{i-1}^{(k)}\}_{k=1}^{N}$.
      Perturb $\boldsymbol{\theta}^*$ to obtain $\boldsymbol{\theta'}$ using a perturbation kernel $K_i(\boldsymbol{\theta'}|\boldsymbol{\theta}^*)$.
      **if** $p(\boldsymbol{\theta'}) = 0$ **then**
        **continue**
      **end if**
      **for** $m = 1$ to $B_i$ **do**
        Simulate the model, compute the summary statistics and the distance $d_{m,j}$ to the observations
      **end for**
      Set $b_i^{(j)}$ the number of times $d_{m,j} < \epsilon_i$
    **end while**
    Set $\boldsymbol{\theta}_i^{(j)} = \boldsymbol{\theta'}$.
    Calculate the weight $w_i^{(j)}$ using eq. (D.1).
  **end for**
  Normalize the weights $\{w_i^{(j)}\}_{j=1}^{N}$.
**end for**

The weights are calculated using the following equation:

$$w_i^{(j)} = \frac{p[\boldsymbol{\theta}_i^{(j)}]\ b_i^{(j)}}{\sum_{k=1}^{N} w_{i-1}^{(k)} K_i\left(\boldsymbol{\theta}_{i-1}^{(k)}, \boldsymbol{\theta}_i^{(j)}\right)} \tag{D.1}$$

The method relies on a proper choice of:

- sequence of tolerances $\{\epsilon_i\}_{i=1}^{N_i}$ with $N_i$ the number of iterations,
- perturbation kernels $\{K_i\}_{i=1}^{N_i}$,
- number of times we have to compute the summary statistics for a given particle $\{B_i\}_{i=1}^{N_i}$.

We choose $N_i = 17$ and:

$$\{\epsilon_i\}_{i=1}^{N_i} = [250, 200, 150, 100, 80, 70, 60, 50, 40, 30, 25, 23, 20, 18, 16, 15, 14]$$

$$\{B_i\}_{i=1}^{N_i} = [1, 1, 1, 1, 1, 1, 1, 1, 2, 2, 3, 3, 3, 4, 4, 4, 5]$$

We choose component-wise uniform perturbation kernels. Those kernel are defined as follows:

$$K_{i,k}(\theta'_k \mid \theta^*_k) \sim \mathcal{U}([\theta^*_k - \varepsilon_{k,i}, \theta^*_k + \varepsilon_{k,i}])$$

where $k$ indexes the component (the parameter) and $i$ the iteration. $\mathcal{U}$ denotes a uniform distribution. We progressively decrease the value of $\varepsilon_{k,i}$ over the iterations of the ABC-SMC algorithm.

For parameters $\mu_1$, $\mu_2$, $\mu_3$, $\mu_4$, we use the uniform perturbation kernel with:

$\{\varepsilon_{\mu_k,i}\}_{i=1}^{N_i} = [0.9, 0.65, 0.525, 0.463, 0.431, 0.416, 0.408, 0.404, 0.402, 0.401, 0.4, 0.24, 0.208, 0.202, 0.2, 0.2, 0.2]$

For parameters $\sigma_1$, $\sigma_2$, $\sigma_3$, $\sigma_4$, we use a uniform perturbation kernel with:

$\{\varepsilon_{\sigma_k,i}\}_{i=1}^{N_i} = [0.7, 0.456, 0.316, 0.230, 0.178, 0.147, 0.128, 0.117, 0.110, 0.106, 0.1, 0.1, 0.1, 0.1, 0.1, 0.1, 0.1]$

For parameter $\eta$, we use:

$\{\varepsilon_{\eta,i}\}_{i=1}^{N_i} = [0.7, 0.456, 0.316, 0.230, 0.178, 0.147, 0.128, 0.117, 0.110, 0.106, 0.1, 0.1, 0.1, 0.1, 0.1, 0.1, 0.1]$

For parameters $p_{l,m}$ $(1 \leq l \leq m \leq 3)$, we use a uniform perturbation kernel with:

$\{\varepsilon_{p_{l,m},i}\}_{i=1}^{N_i} = [0.4, 0.19, 0.127, 0.108, 0.102, 0.101, 0.1, 0.1, 0.1, 0.1, 0.1, 0.08, 0.08, 0.08, 0.08, 0.08, 0.08]$

We also have the possibility to move from a scenario to another one. We have $K(H' = 0 \mid H^* = 1) = 0.3$ and $K(C' = 0 \mid C^* = 1) = 0.3$.

The implementation of the kernel can be found in `/ABC_SMC/kernel.jl`

We display the results for each iteration on Fig. D.3. At each intermediate iteration $i$, as for the last iteration, the marginal posterior probability of each scenario is given by:

$$\mathbb{P}[H = m, C = n] \approx \sum_{j=1}^{N} w_i^{(j)} \mathbb{1}_{H_i^{(j)}=m} \mathbb{1}_{C_i^{(j)}=n}$$

Here, $H_i^{(j)}$ and $C_i^{(j)}$ are binary parameters - indicating in which scenario we are in - which can be considered as components of the parameter vector $\boldsymbol{\theta}_i^{(j)}$. To note that we have the equivalence $\{H_i^{(j)} = 0\} \Leftrightarrow \{\rho_{H_i}^{(j)} = 0\}$ and $\{H_i^{(j)} = 1\} \Leftrightarrow \{\rho_{H,i}^{(j)} > 0\}$ such that the problematic of model (scenario) selection can be seen as a problematic of parameter estimation where the prior for parameter $\rho_H$ would be zero inflated. The same goes for $C$ and $\rho_C$.

We display in Fig. D.4 how the approximated marginal probabilities of our scenarios change over the iterations.
In Fig. D.5, we display the evolution of the distribution of the averaged distance (for all $N$ particles) over the iterations. We can see how the ABC-SMC algorithm improves the results from iteration number 10, compared to the ABC-rejection sampling method. From iteration number 15, there are only slight differences between the distributions of the distances, suggesting that the convergence of the algorithm would already have been obtained at iteration 15.

Compared to the ABC rejection method, we can explore more efficiently the parameter space with the ABC-SMC method. Finally, after the final iteration (iteration $i = 17$), we get:

- $\mathbb{P}[H = 1, C = 1|\mathcal{D}] = 72.69\%$
- $\mathbb{P}[H = 1, C = 0|\mathcal{D}] = 27.31\%$
- $\mathbb{P}[H = 0, C = 1|\mathcal{D}] = 0.0\%$
- $\mathbb{P}[H = 0, C = 0|\mathcal{D}] = 0.0\%$

We retrieve the same main conclusions as previously: 1) the hypothesis of homogeneity in fate ($H = 1$) being crucial to explain our observations (the two scenarios where $H = 0$ being now rejected), and 2) that the scenario without concordance ($H = 1, C = 0$) could still explain the experimental observations, with a marginal posterior distribution here equal to 0.27.

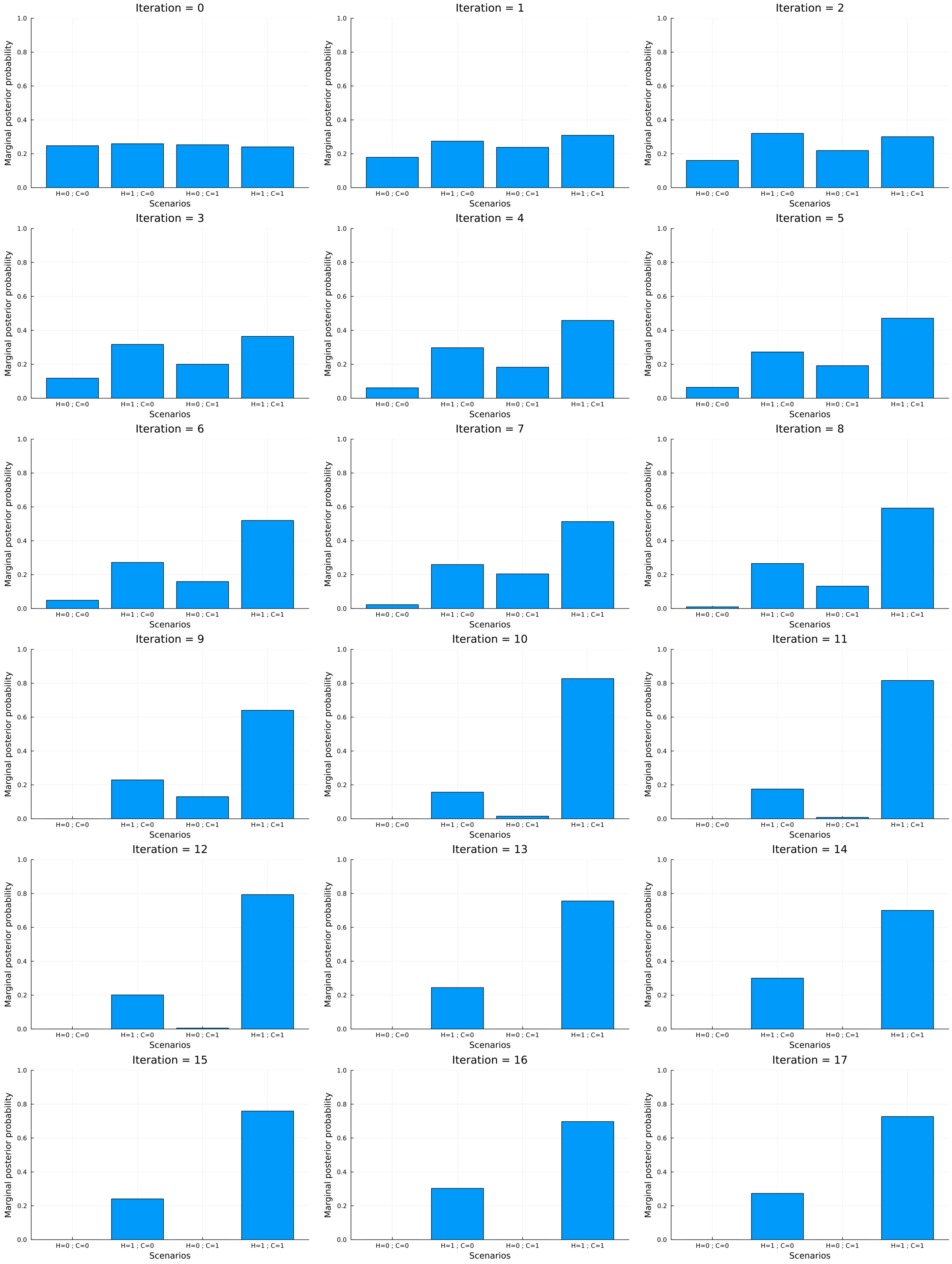

Figure D.3: Intermediate marginal Posterior Distributions of each of our scenarios. From left to right: $(H = 0, C = 0)$, $(H = 1, C = 0)$, $(H = 0, C = 1)$, $(H = 1, C = 1)$.

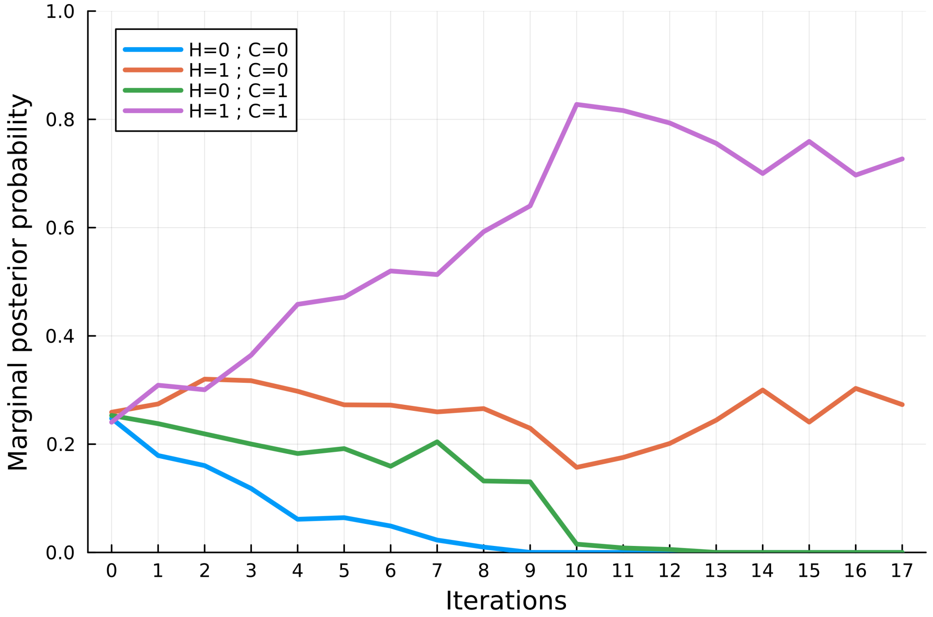

Figure D.4: Evolution of the approximated marginal probabilities of our scenarios over iterations.

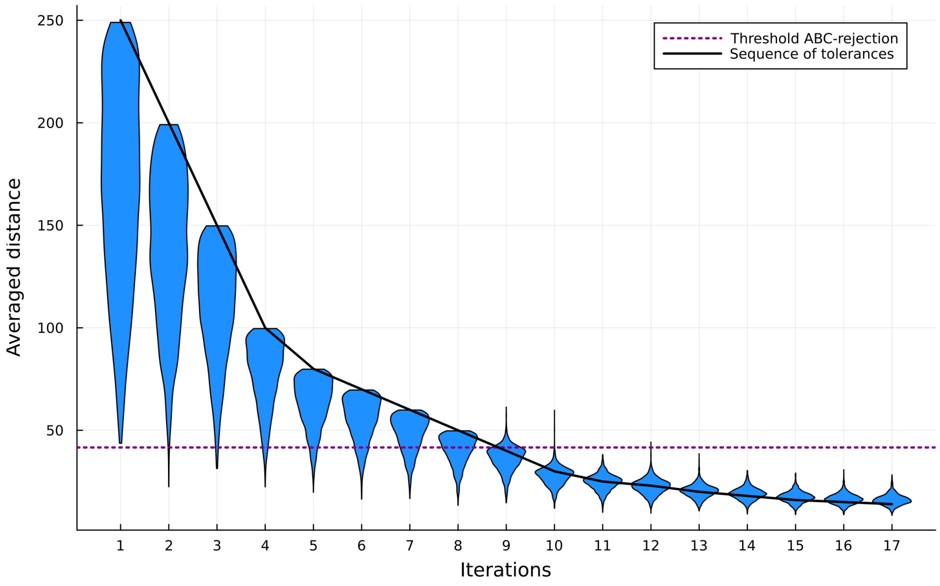

Figure D.5: For each iteration $i$, we display the distribution of the (averaged over $B_i$ simulations) distances of all particles. The horizontal purple line corresponds to the threshold obtained previsouly with the ABC-rejection sampling. The black line corresponds to the decreasing sequence of tolerances. For iterations such that $B_i = 1$, then the distances are below the threshold. Otherwise, the particles are selected when at least one simulation over the $B_i$ is below the tolerance.

##### D.3 Robustness of the ABC-SMC sampling methods: exploring different kernels

To verify to which extent our previous estimations are robust, we explore how changing the sequence of kernels impact the results of the estimations.

###### D.3.1 Considering different configurations for the kernels

**Configuration 2:** Same perturbations kernel as for configuration 1 (that is, the one presented in § D.2), except that we now consider a lower probability to move from a scenario to another one: $K(H' = 0 \mid H^* = 1) = 0.2$ and $K(C' = 0 \mid C^* = 1) = 0.2$.

**Configuration 3:** Compared to configuration 1, the differences are the following:
For parameters $\sigma_1$, $\sigma_2$, $\sigma_3$, $\sigma_4$, we use a uniform perturbation kernel with:

$$\{\varepsilon_{\sigma_k,i}\}_{i=1}^{N_i} = [0.85, 0.69, 0.56, 0.46, 0.38, 0.31, 0.26, 0.22, 0.18, 0.16, 0.05, 0.05, 0.05, 0.05, 0.05, 0.05, 0.05, ]$$

For parameters $p_{l,m}$, we use a slightly different uniform perturbations kernel compared to configuration 1, with differences for the last iterations:

$$\{\varepsilon_{p_{l,m},i}\}_{i=1}^{N_i} = [0.4, 0.19, 0.127, 0.108, 0.102, 0.101, 0.1, 0.1, 0.1, 0.1, 0.1, 0.062, 0.06, 0.06, 0.06, 0.06, 0.06]$$

**Configuration 4:** This configuration differs completely from configuration 1, but we still use uniform perturbation kernels.
For parameters $\mu_1$, $\mu_2$, $\mu_3$, $\mu_4$, we use the uniform perturbation kernel with:

$$\{\varepsilon_{\mu_k,i}\}_{i=1}^{N_i} = [1.0, 0.9, 0.8, 0.7, 0.75, 0.6, 0.55, 0.5, 0.45, 0.4, 0.35, 0.3, 0.25, 0.2, 0.18, 0.16, 0.12]$$

For parameters $\sigma_1$, $\sigma_2$, $\sigma_3$, $\sigma_4$, we use a uniform perturbation kernel with:

$$\{\varepsilon_{\sigma_k,i}\}_{i=1}^{N_i} = [0.7, 0.6, 0.65, 0.6, 0.4, 0.3, 0.2, 0.15, 0.1, 0.1, 0.09, 0.09, 0.08, 0.08, 0.07, 0.07, 0.07]$$

For parameter $\eta$, we use:

$$\{\varepsilon_{\eta,i}\}_{i=1}^{N_i} = [0.8, 0.7, 0.6, 0.5, 0.4, 0.3, 0.3, 0.25, 0.2, 0.15, 0.15, 0.13, 0.12, 0.11, 0.1, 0.09, 0.08]$$

For parameters $p_{l,m}$, we use a uniform perturbations kernel with:

$$\{\varepsilon_{p_{l,m},i}\}_{i=1}^{N_i} = [0.4, 0.3, 0.2, 0.15, 0.12, 0.11, 0.1, 0.09, 0.08, 0.07, 0.06, 0.05, 0.045, 0.04, 0.04, 0.035, 0.03]$$

We also have the possibility to move from a scenario to another one. We have $K(H' = 0 \mid H^* = 1) = 0.4$ and $K(C' = 0 \mid C^* = 1) = 0.4$.

###### D.3.2 Results

We run the three additional configurations described previously, with $N = 1,000$ particles. We show in Fig. D.6 how the estimations differ compared to the main configuration (configuration 1) which was described in § D.2. We see that our results are consistent over different choices of perturbation kernels.

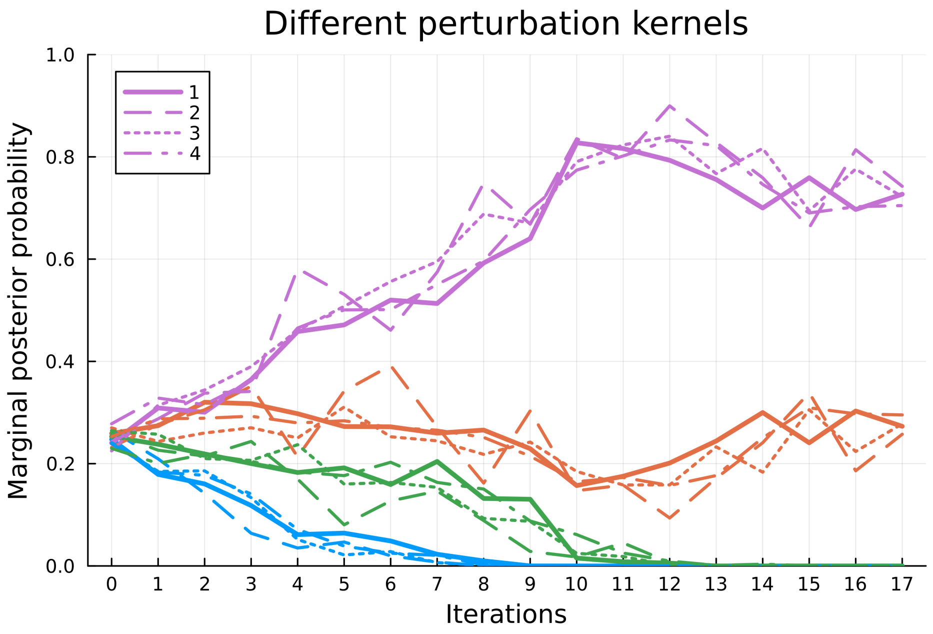

Figure D.6: Evolution of the approximated marginal probabilities of our four scenarios (purple: $H = 1, C = 1$; orange: $H = 1, C = 0$; green: $H = 0, C = 1$; blue: $H = 0, C = 0$) over iterations when using different configurations (different perturbation kernels). The numbers in the legend indicate the configuration, with configuration 1 being the main one, described in § D.2.

#### D.4 Sensitivity analysis

As mentioned by Toni et al. [70]: *"ABC SMC provides us with a global parameter sensitivity analysis (Sanchez & Blower 1997 [S11]) on the fly as the intermediate distributions are being constructed"*. Therefore, it is interesting to look at the intermediate approximate posterior distributions since they give information about the parameters for which the model is the most sensitive.

We display on Figure D.7, for each parameter, the intermediate marginal posterior distributions (conditioned to $H = 1$ and $C = 1$), from the prior to the last iteration.
Not all marginal posterior distributions will strongly deviate from the prior. It is interesting to see that the parameters for which there are few differences between the prior and posterior are those associated to HSC (parameters $p_{1,1}, p_{1,3}, \sigma_1$). We observe that some intermediate distributions deviate rapidly from the prior and quickly converge to the posterior (for example parameters $\mu_k$ for $k \in \{1, 2, 3, 4\}$ or $\eta$). These parameters are those for which the model is the most sensitive. For other parameters, as $\sigma_2$ or $p_{1,2}$, the intermediate distributions are close to the prior for the first iterations, and mostly deviate from it for latter iterations (approximately from iteration 10). These parameters have therefore a lower influence on the model.

To further study that point, we evaluate the Wasserstein distance between each successive intermediate marginal distribution, as well as between the prior and approximate marginal posterior (iteration 17). See the notebook: `/post_treatment/ABC_SMC/sensitivity_analysis.ipynb`
We compare on Fig. D.8 the results, by grouping together similar parameters. Concerning $\mu_k$ and $\sigma_k$, for $k \in \{1, 2, 3, 4\}$, it is interesting to note that, the more mature the cell, the higher the Wasserstein distance between the marginal prior and posterior distribution. Already after the first iteration, there is a significant deviation from the prior for parameters $\mu_4$ and $\mu_3$.
Over the first iterations (approximately until the 8th iteration), the differences between the marginal distributions are more important for $\mu_4$, $\mu_3$, and $\sigma_4$, when the other parameters ($\mu_2$, $\sigma_1$, $\sigma_2$) take over after that iteration.
Concerning the parameters $\sigma_k$, for $k \in \{1, 2, 3, 4\}$, we observe a Wasserstein distance between the prior and posterior distributions two-fold higher for parameter $\sigma_4$ compared to $\sigma_2$ and $\sigma_3$, and 4-fold higher compared to $\sigma_1$. That is, the timing of division of the HSPC is - according to our data, our model,

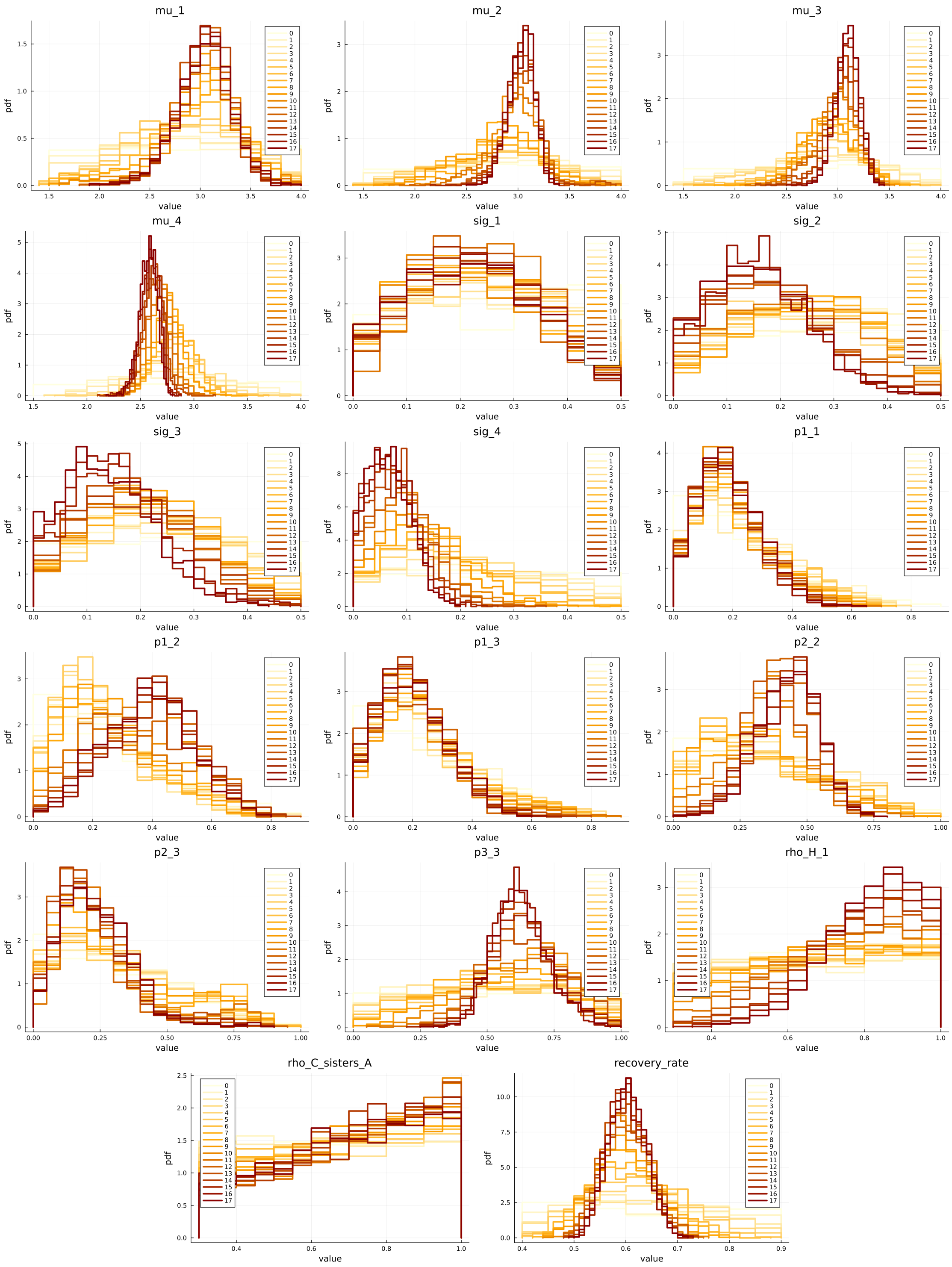

Figure D.7: Intermediate marginal posterior distributions of each of the model parameters (for scenario $H = 1$ and $C = 1$), from iteration 0 (the prior) to iteration 17 (our approximated posterior). Lighter colors correspond to the first iterations when darker colors correspond to the last ones.

and the way we constructed our distance - more identifiable for less immature type (4: $CD34^-$) than for HSC (type 1).

We get similar observations concerning the parameters $p_{i,j}$ (for a transition from type $i$ to type $j$, with $1 \leq i \leq j \leq 3$), where the marginal intermediate posterior distributions of parameters $p_{3,3}$ and $p_{2,3}$ (related to the transition to type 3:HPC) deviate faster from the prior compared to the other parameters.

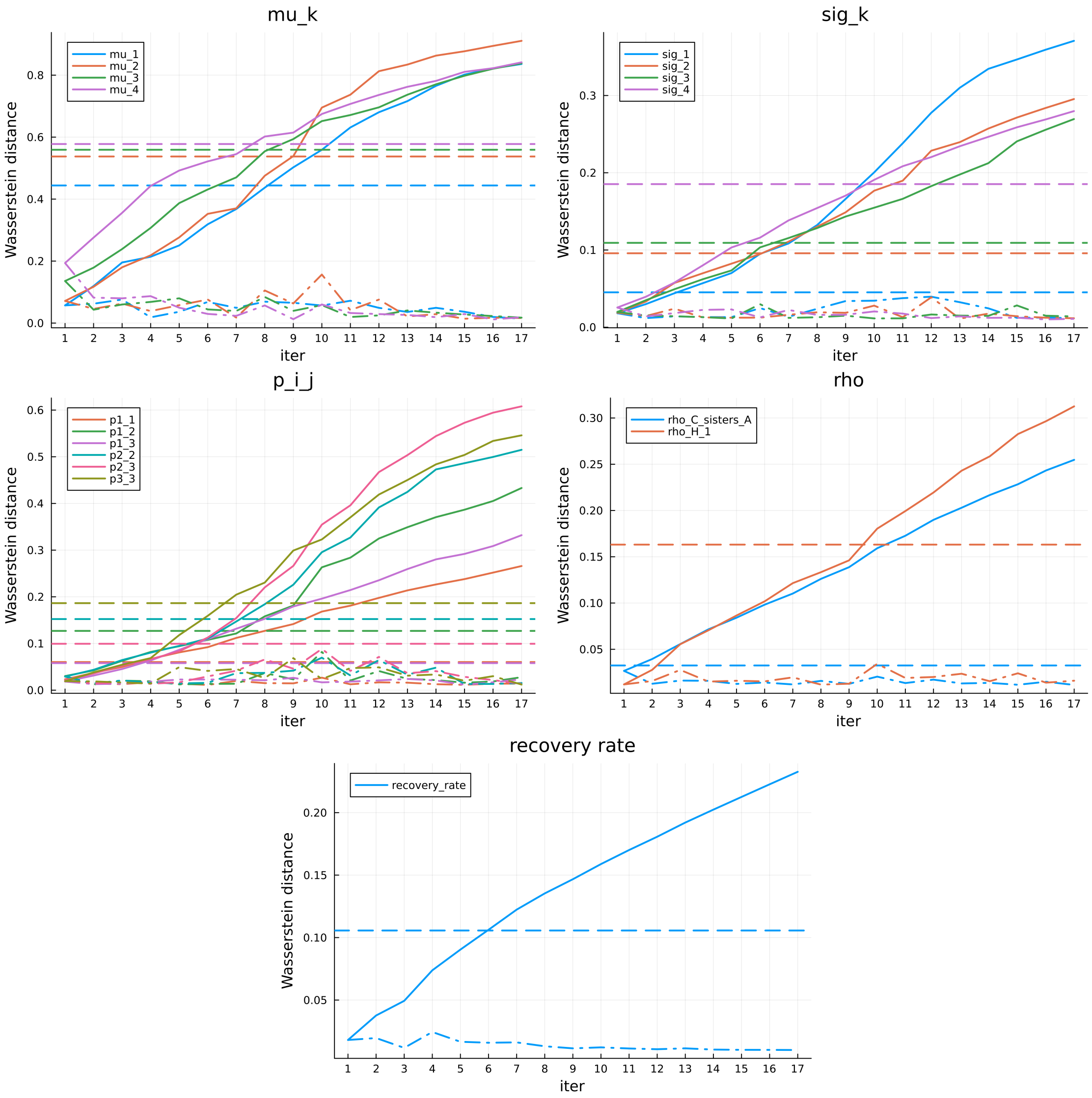

Figure D.8: Wasserstein distances 1) between the marginal posterior distributions and the priors (horizontal dashed line), 2) between each intermediate marginal distribution (dash-doted lines) and 3) cumulated (solid lines), for each model parameter.

#### D.5 Quality of the fits

Finally, to verify that we get good results with our ABC-SMC inference method, we should evaluate the quality of the fits. In our case, a good fit to the data consist of finding parameters values such that the model simulations lead to computed summary statistics which are close to those computed based on the experimental observations.

Even if the distance between our experimental observations and the model simulations is based on linear combinations of the original 340 summary statistics (after having realized a PCA), we can go back to the original summary statistics and evaluate to which extent each of them is close to the experimental value, when sampling from the posterior.

To do that, we consider our $N = 2,000$ particles which approximate our posterior distribution and, for each of them, run a simulation to compute the associated summary statistics. We therefore get an approximation of the posterior distribution of each of our 340 summary statistics, that we compare to the summary statistics computed based on the experimental observations.

The results are displayed in Fig. D.9 for six of them, and the results for all 340 summary statistics are available on: `/post_treatment/ABC_SMC/summary_stats/`.

On these figures, we also display some uncertainty we have concerning the experimental summary statistics. This uncertainty is computed by calculting 100 times the summary statistics, each time on the dataset where we remove 10% of the data.

Overall, we get a good agreement for most of the summary statistics. By analyzing in more detail the summary statistics for which there are more discrepancies, we might potentially determine areas for improving (and complexifying) the model. Yet, it cannot be excluded that we could get better fit to the summary statistics by finding a better way to construct our distance to the observations. For now, we based this latter on a dimension reduction technique, namely the PCA. Non-linear methods could also be explored, as well as subset selection methods, as presented in the review from Blum et al. [S2]. Then, before complexifying the model, it would be worth to investigate whether we could find better way to construct our distance to the observations, to integrate more information from the data. Only then, we should consider further complexifying the model.

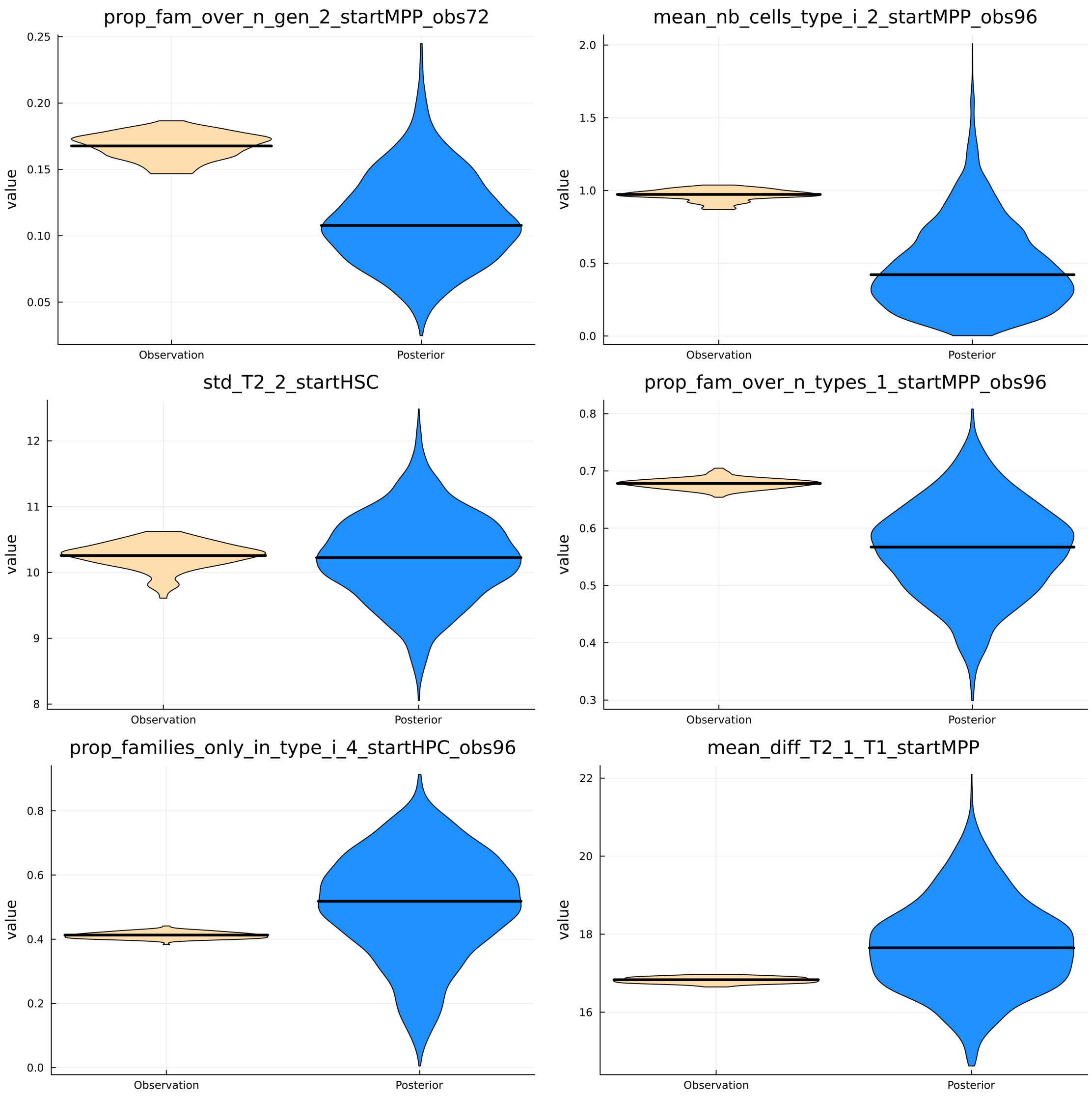

Figure D.9: For several summary statistics, comparison of the value based on the experimental observations (on the left, yellow color, the distribution represents the uncertainty, obtained when randomly sampling 90% of the observations for computing the summary statistic) to those obtained from the mathematical model, when sampling from the posterior distributions (on the right).

### E Considering more scenarios

In this part, we consider more complex scenarios $(C, H)$ with $H \in \{0, 1, 2\}$ and $C \in \{0, 1, \cdots, 5\}$ (§ E.1- E.3) as well as a more parsimonious - minimal - scenario (§ E.4).
Here, to estimate the posterior probability of each scenario, we will use an ABC rejection sampling method, as done in § D.1.

#### E.1 Generating a reference table for the ABC method

For the model selection step, we executed 10,982,900 simulations, each time sampling a parameter vector from the prior.
We first examine the distribution of distances calculated between each simulation and the experimental observations (Figure E.1). We then choose to retain only the top 0.1% of simulations. These will be used to approximate the posterior probability of each our scenario.

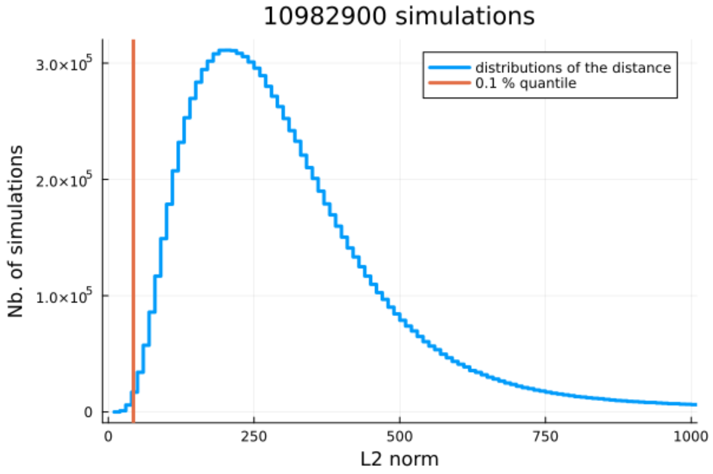

Figure E.1: Distribution of distances to experimental observations across the 10 million+ simulations. The orange vertical line indicates the tolerance associated with the 0.1% percentile. Scenarios (and parameters) leading to simulations with a distance less than this tolerance will be retained to approximate the posterior probabilities of each scenario. Only one in a thousand simulations is thus retained.

In terms of model selection, various scenarios are studied based on the values of the tuple $(H, C)$. The objective is to approximate the posterior distribution of this tuple using an ABC rejection sampling method, consisting of simulating a high number of time the process, and retaining only those for which the computed summary statistics were closes to the ones calculated based on the experiments.

#### E.2 Results

Figure E.2 shows the prior probability of each scenario and their posterior one. The model with the highest posterior probability is the one with:

- Identical homogeneity in fate for the three cell types (HSC, MPP, and HPC) $(H = 1)$
- Identical concordance among all sister cells $(C = 1)$

Rather than looking at the joint prior and posterior distributions of the tuple $(H, C)$, we can also consider the marginal distributions, as shown in Figure E.3. Regarding homogeneity in fate (Figure E.3b), scenarios without this hypothesis are clearly rejected. The scenario with different values of $\rho_{H,a}$ according to mother cell type $a$ shows a higher posterior probability than its prior but is still lower than the configuration where homogeneity would be the same no mater the cell type. Models where $H = 2$ are less parsimonious, with two additional degrees of freedom compared to those where $H = 1$.

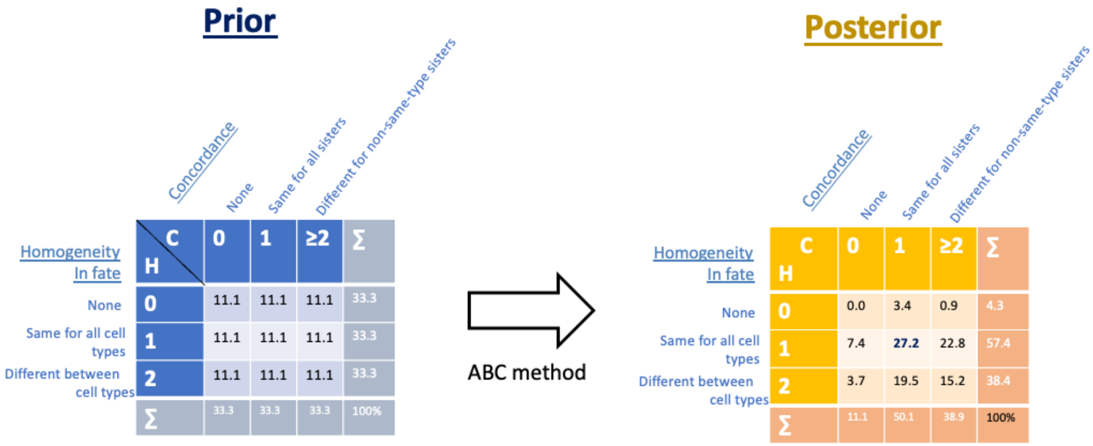

Figure E.2: Prior (left in blue) vs posterior (right in yellow) probabilities (in %) of different scenarios. A scenario is defined by an hypothesis of homogeneity in fate ($H$), and of concordance ($C$). This figure shows how probabilities are updated after confrontation with experimental data. Models without homogeneity ($H = 0$) are found to be very unlikely. The posterior probability of the best scenario is highlighted in bold.

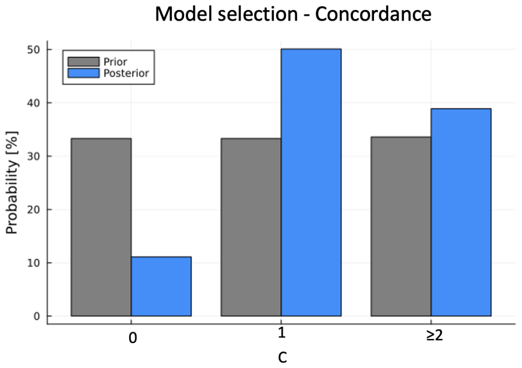

(a) Comparison between the models according to the concordance scenarios $C$.

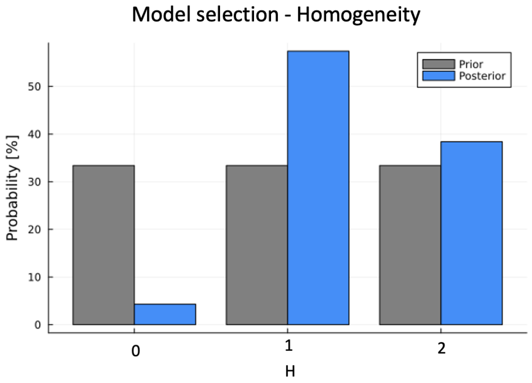

(b) Comparison between the models according to the Homogeneity scenarios $H$.

Figure E.3: Marginal probabilities of concordance (left) or homogeneity (right) scenarios *a priori* (in gray) and *a posteriori* (in blue)

Regarding concordance, scenarios that do not integrate it have only a 11.1% chance of leading to results close to those obtained experimentally (Figure E.3a). The hypothesis of identical concordance for all sister cells ($C = 1$) appears more likely than the hypothesis that this concordance would differ according to certain criteria ($C \geq 2$).

#### E.3 What about the more complex models?

We present in Fig. E.4 and E.5 the results when considering also the scenarios implying more complex hypotheses concerning the concordance.

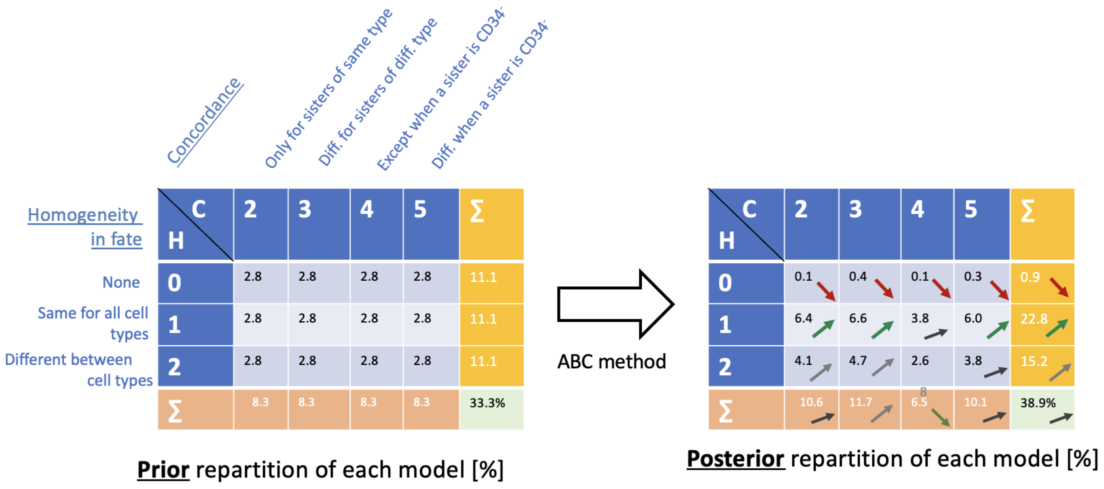

Figure E.4: Prior (left) vs posterior (right) probabilities (in %) of each scenarios implying more complex hypotheses for the concordance.

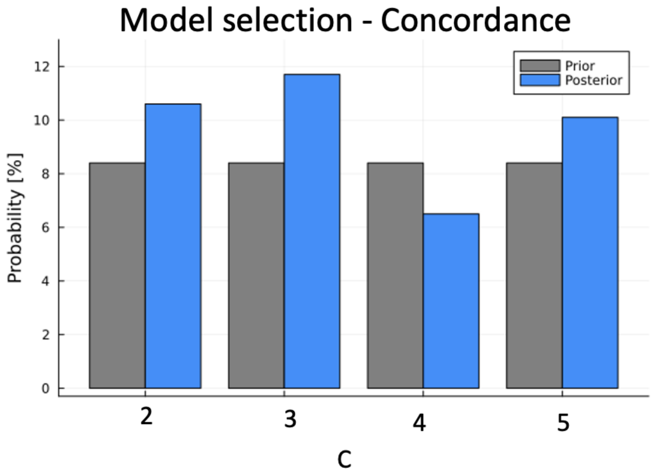

Figure E.5: Marginal probabilities of the more complex concordance scenarios *a priori* (in gray) and *a posteriori* (in blue)

#### E.4 Minimal scenario

We also tested whether an even simpler scenario could be appropriate. In this scenario, all cell divisions (except the first division) follow the same distribution. For any cell $j$ of type $p_j$, the division time follows a lognormal distribution:

$$T_j \mid p_j \sim \mathcal{LN}(\mu, \sigma).$$

Hence, the division time is independent of the cell type, i.e.,

$$\mu_1 = \mu_2 = \mu_3 = \mu_4 = \mu,$$

and

$$\sigma_1 = \sigma_2 = \sigma_3 = \sigma_4 = \sigma.$$

We consider the same priors as in Tab. B.1, namely $\sigma \sim \mathcal{U}([0, 0.5])$ and $\mu \sim \mathcal{U}([1.5, 4.0])$. In addition to the 1,733,106 simulations previously performed for the four main scenarios (see § D.1), we ran 443,000 simulations under this minimal scenario. Among the 2,176,106 simulations in total, we retain only those for which the distance to the observations is below a threshold $\epsilon$. This threshold corresponds to the 0.1 percentile of the 2,176,106 computed distances (see Fig. E.6).

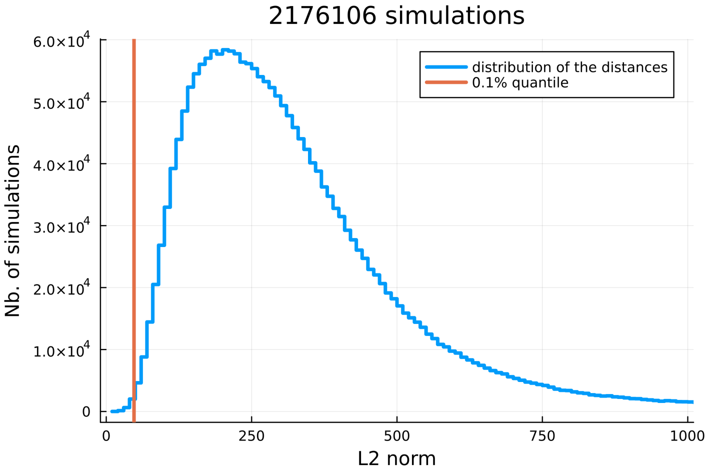

Figure E.6: Distribution of distances to experimental observations across the $\sim 2$ millions simulations (four main scenarios + the minimal scenario). The orange vertical line indicates the tolerance associated with the 0.1% percentile. Scenarios (and parameters) leading to simulations with a distance less than this tolerance will be retained to approximate the posterior probabilities of each scenario.

Among the selected parameter vectors, we compute the frequency of each of the five models; these frequencies approximate the posterior probability of each scenario. We obtain:

- $\mathbb{P}[H = 1, C = 1 \mid \mathcal{D}] \approx 67.62\%$
- $\mathbb{P}[H = 1, C = 0 \mid \mathcal{D}] \approx 23.01\%$
- $\mathbb{P}[H = 0, C = 1 \mid \mathcal{D}] \approx 7.71\%$
- $\mathbb{P}[H = 0, C = 0 \mid \mathcal{D}] \approx 0.14\%$
- $\mathbb{P}[\text{Minimal scenario} \mid \mathcal{D}] \approx 1.52\%$

Including the minimal scenario does not alter the conclusions obtained in § D.1. The minimal scenario is assigned a low posterior probability (1.52%), despite its parsimony, and can therefore be discarded. Its probability is slightly higher than $\mathbb{P}[H = 0, C = 0 \mid \mathcal{D}]$, which is expected given its reduced number of parameters to estimate, but both remain negligible.
